# Rules of engagement for condensins and cohesins guide mitotic chromosome formation

**DOI:** 10.1101/2024.04.18.590027

**Authors:** Kumiko Samejima, Johan H. Gibcus, Sameer Abraham, Fernanda Cisneros-Soberanis, Itaru Samejima, Alison J. Beckett, Nina Pučeková, Maria Alba Abad, Bethan Medina-Pritchard, James R. Paulson, Linfeng Xie, A. Arockia Jeyaprakash, Ian A. Prior, Leonid A. Mirny, Job Dekker, Anton Goloborodko, William C. Earnshaw

## Abstract

During mitosis, interphase chromatin is rapidly converted into rod-shaped mitotic chromosomes. Using Hi-C, imaging, proteomics and polymer modeling, we determine how the activity and interplay between loop-extruding SMC motors accomplishes this dramatic transition. Our work reveals rules of engagement for SMC complexes that are critical for allowing cells to refold interphase chromatin into mitotic chromosomes. We find that condensin disassembles interphase chromatin loop organization by evicting or displacing extrusive cohesin. In contrast, condensin bypasses cohesive cohesins, thereby maintaining sister chromatid cohesion while separating the sisters. Studies of mitotic chromosomes formed by cohesin, condensin II and condensin I alone or in combination allow us to develop new models of mitotic chromosome conformation. In these models, loops are consecutive and not overlapping, implying that condensins do not freely pass one another but stall upon encountering each other. The dynamics of Hi-C interactions and chromosome morphology reveal that during prophase loops are extruded in vivo at ∼1-3 kb/sec by condensins as they form a disordered discontinuous helical scaffold within individual chromatids.

## INTRODUCTION

The SMC protein complexes cohesin and condensin are key determinants of chromatin organization in interphase and in mitosis (*1–4*). During prophase, a cohesin-dominated interphase organization with topologically associating domains (TADs) and compartments transitions into a condensin-dominated mitotic loop array (*5*, *6*). Therefore, encounters between cohesins and condensins are likely to be frequent during this time. When bacterial SMC complexes collide, they can bypass each other (*7*), as modeling and Hi-C studies (*8*) suggest, consistent with single-molecule studies of yeast condensin (*9*). In contrast, the “rules of engagement” for when condensins encounter cohesins and each other during mitotic chromosome formation in metazoa remain underexplored.

SMC complexes generate loops via ATP-dependent loop extrusion (other mechanisms may also contribute to loop formation) (*1*, *10–14*). Throughout interphase, extruding cohesin complexes organize the genome into loops that are potentially important for regulating gene expression (*15*). Also, cohesive cohesin complexes establish cohesion during S-phase, and hold sisters together until anaphase (*16–19*). Aside from its role in maintaining the pairing of sister chromatid arms in early mitosis (*20–22*) cohesin has also been suggested to antagonize chromatid axis formation in Xenopus extracts (*23*).

Most cohesin is released from chromosome arms during prophase (*24*, *25*). Also, at this time, condensin II in the nucleus is activated by CDK1 and Plk1 (*26–28*), associates stably with the chromatin, and begins to extrude a sequence-non-specific loop array. Thus, a dynamic exchange of SMC complexes occurs throughout prophase. After nuclear envelope breakdown, cytoplasmic condensin I associates with chromosomes, forming loops nested within the larger condensin II loops (*6*, *29*). The resulting densely packed “bottlebrush-like” arrays of consecutive loops produce the classical rod-shape of mitotic chromosomes (*30*) with paired sister chromatids aligned along their length (*31–33*).

Here we describe a series of single- and multiple-mutant degron cell lines that enable us to examine the action of specific combinations of cohesin, condensin I or condensin II complexes in chromosome formation during highly synchronous mitotic entry. Hi-C and microscopy data have (1) revealed that cohesive, and not extrusive, cohesin has a significant effect on the structure of mitotic chromosomes; (2) determined how SMC complexes engage with each other during chromosome formation; (3) allowed us to create models for chromosome organization formed by SMC complexes acting singly and in combination, and (4) allowed us to determine the speed of loop extrusion by condensins in vivo.

## RESULTS

### Substantial levels of cohesin remain on prometaphase chromosomes

We quantified the temporal relationships between chromosome condensation, nuclear envelope breakdown (NEB), accessibility of cytoplasmic factors to chromatin, and the amount of SMC complexes bound to chromatin during mitotic entry and progression. We used DT40 CDK1^as^ cells that undergo a highly synchronous mitotic entry following release from a G_2_ arrest with 1NM-PP1, which selectively inhibits CDK1^as^ by preventing ATP binding (*6*, *34*) (Fig. S1A,B). Cells expressing Halo-Lamin B1 and 3x GFP-NES were used to monitor nuclear envelope integrity (Fig. 1A,B).

**Fig. 1.**
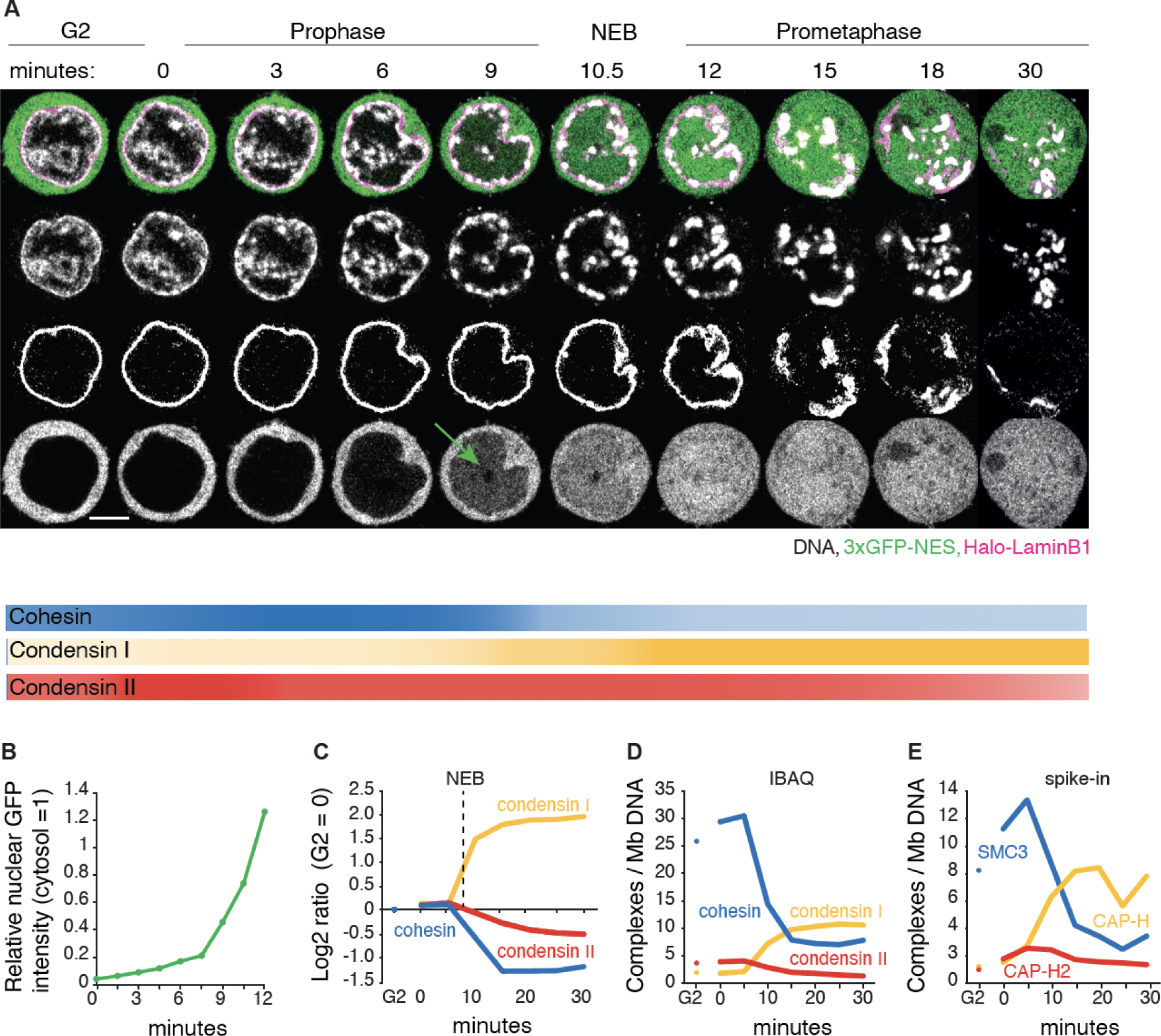
Dynamics of chromatin bound cohesin and condensin during mitotic entry. (**A**) Representative live cell imaging of a DT40 CDK1^as^_Halo-lamin B1_3xGFP-NES cell released from G_2_ block with 1NM-PP1. DNA:grey; Halo-lamin B1: magenta; 3xGFP-NES: green. Seven z-sections (1 µm interval) were taken every 1.5 min. A single section is shown for each timepoint. Scale bar = 5 µm. 3XGFP enters nuclei a few minutes prior to visible nuclear lamina disruption (NEB, t= 10-11 min). Intensity of lines under the images illustrates the relative amount of the indicated complexes on chromatin at each time point. (**B**) Relative nuclear GFP fluorescence intensity (cytosolic GFP intensity = 1) from experiment of Fig. 1A. (**C**) Chromatin enrichment for proteomics (ChEP) analysis of WT CDK1^as^ cells (SILAC analysis). Log_2_ SILAC ratio normalized against G_2_ is shown for cohesin (average of SMC1, SMC3, and RAD21), condensin I (CAP-H, CAP-G, and CAP-D2) and condensin II complexes (CAP-H2, CAP-G2, and CAP-D3). t= 0 is after completion of 1NM-PP1 washout, n= 6. (**D**) Estimated number of chromatin-associated cohesin, condensin I and condensin II complexes (per Mb DNA) during mitotic entry in wild type CDK1^as^ cells. Average iBAQ number from ChEP analysis for subunit as listed in C was normalized relative to values for Histone H4, n= 6. (**E**) Absolute quantification of SMC3 (cohesin subunit), CAP-H (condensin I subunit) and CAP-H2 (condensin II subunit) on chromatin (per Mb DNA). Protein numbers are calculated following ChEP analysis of corresponding Halo-tagged cell lines normalized using purified spike-in Halo-Histone H4 protein (n= 4).

Live cell imaging revealed that chromatin condensation started about 3 minutes after release from G_2_ block (t= 0 min), in early prophase. The condensing chromatin concentrated at the inner surface of the nuclear envelope and around nucleoli, where rod-shaped chromosomes became distinct at t= ∼9 min (late prophase). At t= 10-12 min, the nuclear rim of lamin B1 became discontinuous (NEB), and individual chromosomes were released (prometaphase). Interestingly, fragments of lamin B1 polymer remained associated with the prometaphase chromosomes until at least t= 20 min, consistent with our previous chromatin-enrichment for proteomics (ChEP) study (*35*, *36*)

Chromatin-bound cohesin detected using ChEP was highly abundant in G_2_ (Fig. 1C-E, Fig. S2A). Bound cohesin levels increased slightly in early prophase, then decreased approximately 3-fold during late prophase and after NEB. Condensin II was nuclear during G_2_, with the amount of chromatin-associated condensin II increasing slightly in early prophase, then gradually decreasing by about 50% from prophase to late prometaphase (Fig. 1C-E, Fig. S2B). Condensin I, which is predominantly cytosolic during interphase (*37*, *38*), accumulated on chromatin gradually during late prophase and then rapidly after NEB, ultimately reaching a level about 5x that of condensin II. This behavior of condensin I paralleled that of 3xGFP-NES, which is too large to diffuse through nuclear pores. We observed a gradual increase of GFP intensity in the nucleus several minutes prior to NEB which dramatically increased during and after NEB (Fig. 1A: 9 min, and 1B) (*35*).

Quantitative spike-in-normalized proteomic analysis revealed that late prometaphase chromosomes contained substantial amounts of residual cohesin and newly bound condensin I (∼3-6 and ∼10 complexes per megabase, respectively - Fig. 1D,E). Note that this analysis cannot distinguish between cohesin on the arms and at centromeres, nor distinguish extruding and cohesive cohesin. Below we examine the spatial localization of cohesin on prometaphase chromosomes.

Taken together, these data show that condensin I may access chromatin as early as mid-late prophase, and that a substantial amount of cohesin remains associated with chromatin till late prometaphase (Fig. 1). The co-occurrence of the various SMC complexes raises the possibility that their interactions, including potential clashes, influence mitotic chromosome formation and the removal of interphase chromatin structures. Thus, cohesin could have a previously unsuspected role in mitotic chromosome formation.

### Condensin drives the disassembly of interphase chromatin structures

Using Hi-C, we examined mechanisms driving the dramatic loss of interphase chromatin structures during mitotic entry. As noted in a previous study (*6*) and analyzed quantitatively here, loss of interphase features is condensin dependent. Features lost include TADs, dots and stripes formed by loop-extruding cohesin and global compartments. Here, we studied this phenomenon more quantitatively during prophase using new CDK1^as^ cell lines expressing OsTIR1 plus a series of single and multiple homozygous auxin-inducible degron alleles of condensin I and II (Table S1). This enabled us to mechanistically examine the role of condensin in this process during prophase and test our hypotheses using polymer simulations.

Mitotic cells depleted of SMC2 (i.e. lacking both condensin I and II) entered mitosis with normal kinetics (Fig. S1D), but rod-shaped chromosomes did not form even at late prometaphase (Fig. 2A), consistent with previous data (*6*, *14*, *39–41*). Nonetheless, the chromatin underwent a rapid reduction in volume after NEB. Measuring maximum projection images of those chromosomes in nocodazole-treated cells indicated that the area of all SMC2-depleted prometaphase chromosomes (t= 30 min) decreased approximately 2-fold relative to interphase - corresponding to an approximately 3-fold decrease in volume (Fig. 2A, Table S2). This level of compaction is comparable to the volume decrease observed in normal mitotic cells (*42–44*), and is consistent with our previous conclusion that condensins are not essential for volume compaction of chromosomes in mitosis (*40*). This confirms that mitotic chromosome formation involves condensin-dependent rod formation and condensin-independent volume compaction (*6*, *40*, *45*).

**Fig. 2.**
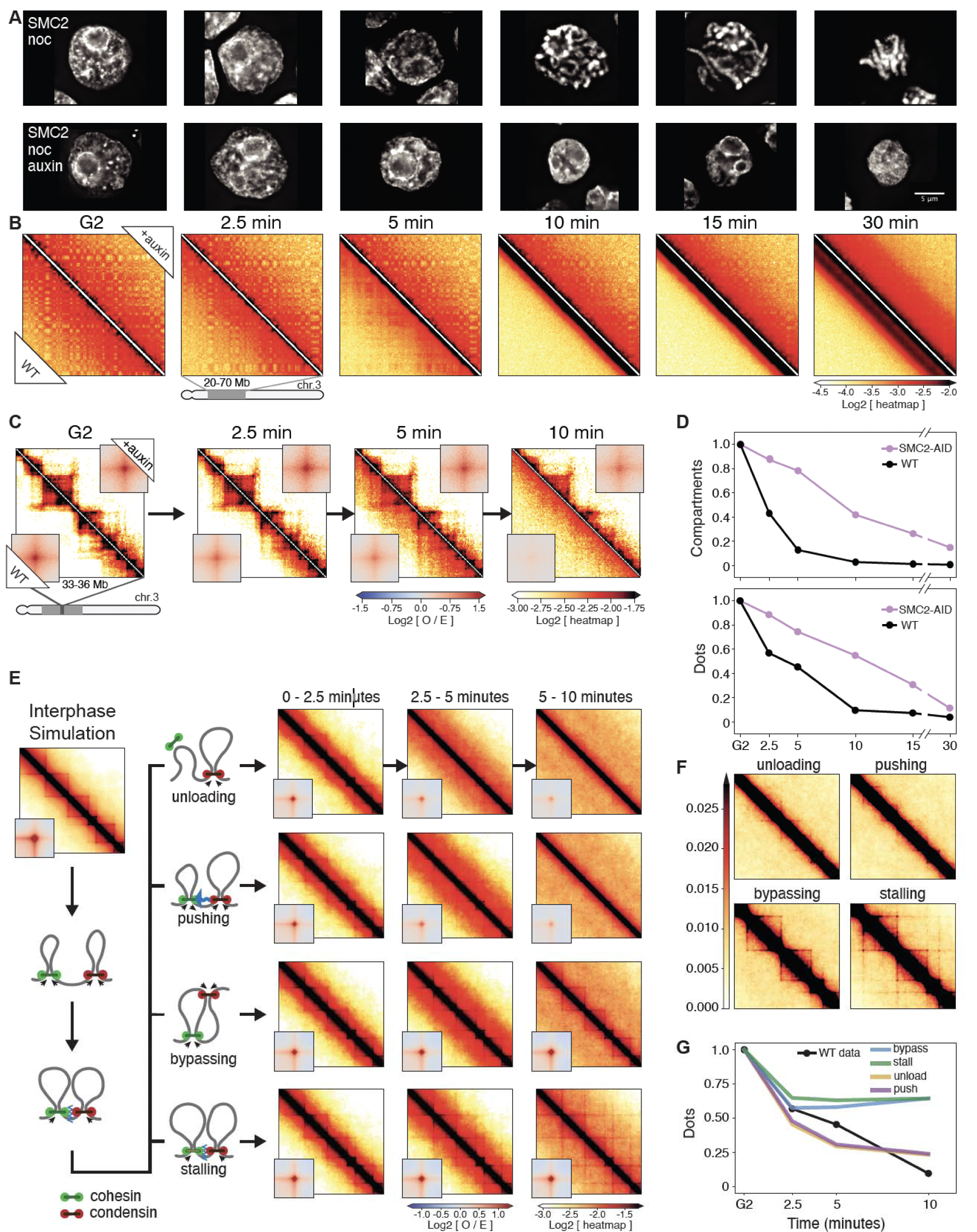
Condensin facilitates disassembly of interphase chromatin organization during mitotic entry. (**A**) Representative images of DAPI-stained SMC2-AID cells used to prepare Hi-C samples. Nocodazole was added 30 minutes prior to 1NM-PP1 washout. Top row: cells from control culture (no auxin treatment). Bottom row: cells from culture treated with auxin during the G_2_ block and release into mitosis. Scale bar = 5 µm. (**B**) Hi-C contact maps of wild-type cells (WT) and SMC2-AID cells treated with auxin (+auxin) from cell populations shown in panel A. A region from chromosome 3 (position 20-70 Mb) is shown. Bottom triangle of each Hi-C map displays Hi-C data obtained with control cells; top triangle displays Hi-C data obtained with auxin treated cells. The checkerboard pattern, readily observed in G_2_ cells, reflects compartmentalization. Compartments disappeared quickly in WT cells but remained in SMC2-depleted mitotic cells. (**C**) Same as in panel B, but zoomed in for chromosome 3 (region 33-36 Mb). Insets show the distance-normalized contact enrichment at detected dots genome-wide. TADs and dots persist in SMC2-depleted mitotic cells. (**D**) Quantification of features shown in panel B (compartments) and panel C (dots). Compartment and dot strength were normalized to their values in G_2_-arrested cells, which was set at 1. (**E**) Outline of four possible simulated scenarios of collisions between cohesin and condensins in prophase (left) and the corresponding simulated Hi-C maps (right, on log scale). (**F**) Same simulated Hi-C maps as in panel E, but in linear scale to better emphasize cohesin-dependent features including lines and dots. (**G**) Quantification of dots in panel E as predicted by the 4 simulated scenarios shown in panel E and in comparison to WT experimental Hi-C data. Dot strength was normalized to G_2_, which was set at 1.

Quantitative analysis of chromosome organization at high temporal resolution (2.5’, 5’, 7.5’, 10’, 15’, 30’ after synchronous release from G_2_) reveals that condensins have an active role in the disassembly of cohesin-dependent interphase features. In WT cells, the strength of compartments (Fig. 2B,D), dots (i.e., CTCF-CTCF loops), and TADs (Fig. 2C,D) decayed rapidly in the first 10 minutes after release from the CDK1^as^ block and became undetectable by the onset of NEB. In contrast, in condensin-depleted cells, interphase chromatin features, including TADs, dots and compartments, persisted until late prometaphase (Fig. 2C,D). Thus, condensin can disrupt compartmental associations, possibly through a mechanism analogous to that reported for how cohesin can weaken compartmentalization through loop extrusion (*46–48*). Additionally, condensin disrupts cohesin-mediated loops. The prolonged presence of “dots” in Hi-C maps till later in prometaphase in condensin-depleted cells reveals that CTCF must remain bound to its cognate sites with cohesin remaining localized to those sites. ChEP data confirm that the loss of CTCF and cohesin from chromosomes is delayed in the absence of condensin (Fig. S2A,C).

We used polymer simulations to explore four possible mechanisms by which condensins could interact with loop-extruding cohesins: when a cohesin and condensin complex meet each other, cohesin is (1) unloaded, (2) pushed along the chromosome by condensin, (3) bypassed by condensin, or (4) the two complexes stall until cohesin unbinds. We simulated these 4 scenarios and quantified the loss of cohesin-mediated features. Our analysis clearly revealed that only unloading and pushing can displace CTCF-stalled cohesins that form dots and recapitulate the fast loss of interphase features (Fig. 2E,F). The slower condensin-independent loss of TADs and dots, observed in SMC2-depleted cells, can be explained by a “background” pathway. This could involve the gradual loss of CTCF and cohesin seen in the proteomics (ChEP) data (Fig S2A,C). In wild type cells, the much faster loss driven by condensin obscures the slower-acting “background” pathway.

We conclude that the rapid disassembly of interphase chromosome folding during mitotic entry is condensin driven and can be explained by disruption of cohesin loops by condensins. In addition to stalling and bypassing observed in prior studies (*9*), our data and simulations suggest that interactions between the complexes can lead to selective unloading and/or pushing of extrusive cohesin. Selective depletion of either condensin I or II shows that condensin II is the most likely driver of this interphase disassembly (Fig. S4).

### Cohesin localizes between sister chromatids away from condensin

We visualized cohesin and condensin in DT40 CDK1^as^ cell lines expressing knock-in SMC3-Clover plus SMC2-Halo. Strikingly, cohesin on prometaphase chromosomes did not co-localize with condensin. Although the cohesin was localized all along the whole length of the chromosomes, it was concentrated between the sister chromatids, away from the condensin axis of each chromatid (Fig. 3A - see also Rhodes et al. (*49*)). This distribution is consistent with the remaining cohesin mediating inter-sister cohesion (i.e. cohesive cohesin) and forming links between the distal portions of chromatin loops, the bases of which are located in the chromatid interior, where the condensin is concentrated (Fig. 3A).

**Fig. 3.**
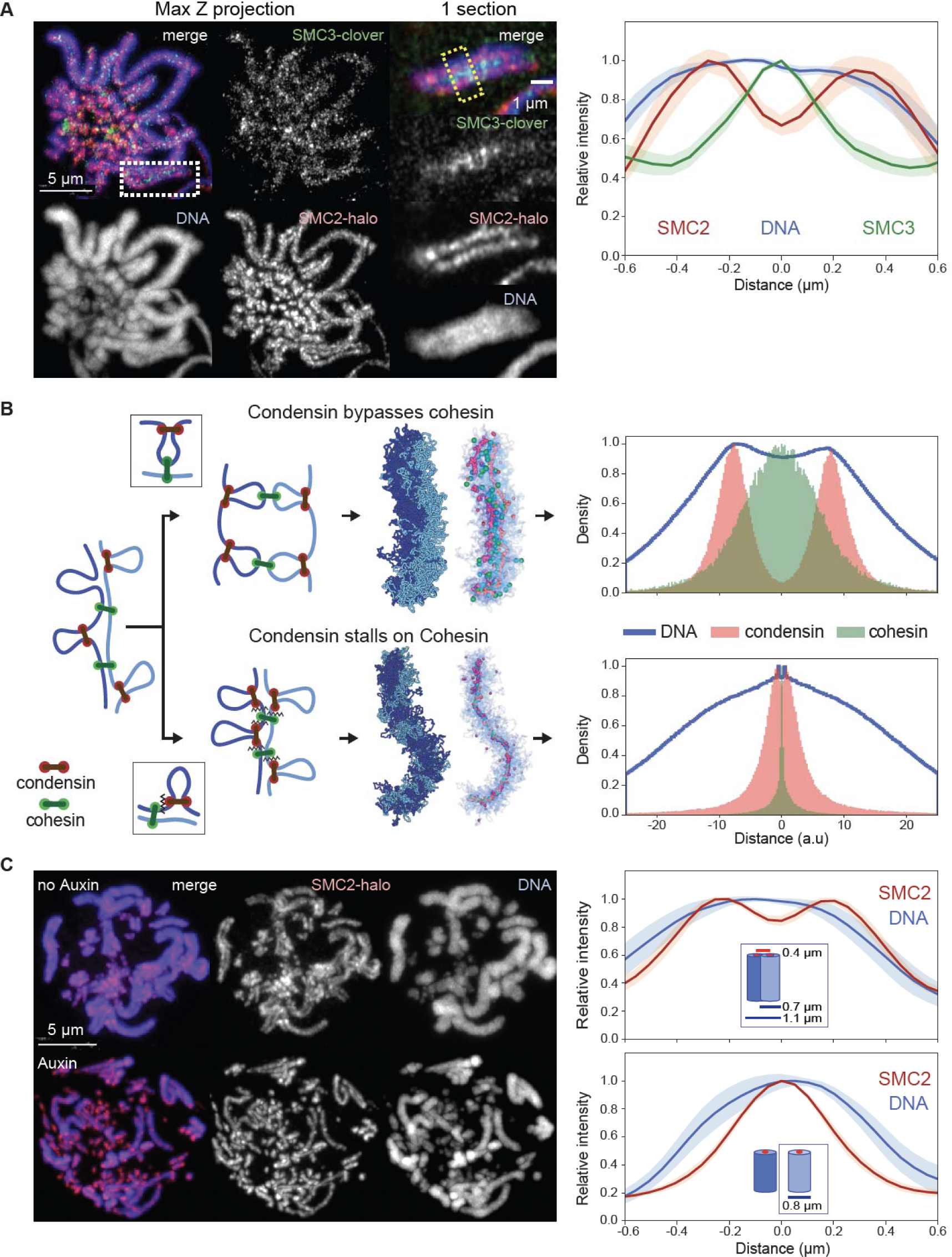
Condensin bypasses cohesin to establish separated but cohesed sister chromatids. (**A**) Localization of cohesin (SMC3) and condensin (SMC2) in prometaphase SMC3-clover/SMC2-Halo CDK1^as^ cells 30 minutes after release from G_2_ block. JFX646-Halo (1/1,000) was added to visualize SMC2-Halo. To remove chromatin-unbound SMC3, cells were fixed with 4% formaldehyde plus 0.5 % Triton. (left), Maximum projection of the full stack of z-sections. (right), Zoom of region in the white rectangle (single z-section). Line scans across chromosomes in the single z-section images (yellow box) were used to quantify the relative fluorescence intensities of SMC3-clover, SMC2-Halo, and DNA (plot to the right). Plots were centered around the position with the highest SMC3 intensity. Data obtained with 10 chromosomes each for two replicate experiments were averaged. Shaded envelopes around main lines represent standard deviations. (**B**) Polymer models of chromatid compaction of pairs of cohesed sister chromatids through condensin-mediated loop extrusion. Left: two mechanisms of interactions between condensin and cohesive cohesins are modeled (top: condensins bypassing cohesive cohesins; bottom: condensins stalling at cohesive cohesins). Polymers of sisters are shown in shades of blue, cohesive cohesins in green, and extruding condensins in red. Middle: simulated outcomes of configurations of sister chromatids obtained with bypassing (top) or stalling models. Right: histograms present the localization of cohesin, condensin, and DNA for cross-sections of pairs of sister chromatids (as in panel A, right). (**C**, left) Representative images of prometaphase SMC3-AID/SMC2-Halo cells without or with auxin treatment to remove cohesin prior to mitotic entry. Cells at t= 30 min after release from G_2_ block were fixed with formaldehyde and SMC2-Halo was visualized with JFX549-Halo (1/10,000). DNA was stained with Hoechst 33542. Max projections of z sections are shown; scale bar = 5 µm. (right) Line-scan quantification of relative fluorescence intensities of DNA and SMC2 across pairs of sister chromatids (no auxin) or single chromatids (+ auxin). Positions with lowest SMC2 intensity (no auxin, marks the point where sister chromatids touch) or highest (auxin, the midpoint of a single chromatid) were aligned in the middle. n = 10 chromosomes for each condition in two replicate experiments. Shadow colors show standard deviation. The calculation of the chromatid/chromosome width is described in the Method section.

### Condensin must bypass cohesive cohesins

Sister chromatid cohesion is established during S-phase ((*50*) and below) and maintained thereafter by stably bound cohesive cohesins. Interactions between cohesive cohesins and condensins, which occur following condensin activation much later in prophase, determine the emerging chromosome morphology and localization of SMCs. To explore possible mechanisms that can lead to localization of cohesive cohesin between sister chromatids and away from condensin scaffolds, we explored possible modes of interaction between condensin and cohesive cohesin. We developed a model with two polymers representing the sister chromatids in G_2_ linked by cohesive cohesin complexes (for simplicity we did not include extrusive cohesins) (*30*). Once condensins start extruding loops as cells enter prophase, they encounter cohesive cohesins (Fig. 3B). Our models explored two possible interactions between condensins and cohesive cohesins: 1) bypassing, in which condensins can step over cohesins, and 2) stalling and/or pushing, in which condensins accumulate at cohesive cohesins or push them ahead of them.

The two mechanisms gave rise to strikingly different organizations of sister chromatids, their loops, and localizations of condensins and cohesins. Firstly, if condensins cannot bypass cohesive cohesin, condensins stall at those sites, resulting in tight colocalization of the two complexes. As a result, the two sister chromatids would share a single axis containing both SMC complexes, in clear disagreement with microscopy (Fig. 3B). In contrast, if condensins can bypass cohesive cohesins, the two sister chromatids would individualize, and each would form a bottlebrush of loops with condensins enriched along its axis and distal cohesins linking loops of the two sisters. Cohesins, in this case, were positioned inside condensin loops, away from loop bases, and at the interface where the two bottlebrushes were connected (Fig. 3B). This distribution is in striking agreement with cohesin and condensin localisation seen in cells (Fig. 3A).

Together, these analyses reveal a complex interplay between cohesins and condensins. First, we showed that condensins actively erase interphase features by mediating either unloading or pushing loop-extruding cohesin. Second, the ability of condensin to bypass cohesive cohesin is essential for the formation of individual cohesed sister chromatids. While SMC complex bypassing has been observed in vitro (*9*) and in bacteria (*8*), such activity in eukaryotes implies that condensin distinguishes between the two types of cohesin complexes: removing or pushing one while bypassing the other.

### Cohesin depletion significantly alters mitotic chromosome organization

Given the complex interplay between condensin and cohesin complexes during mitosis, we determined how mitotic chromosome formation was affected by the depletion of cohesin in G_2_ arrested SMC3-AID/CDK1^as^ cells, prior to synchronous mitotic entry. Following the depletion of cohesin, sister chromatids appeared separated by prometaphase (Fig. 3C), and quantification of chromatin -associated proteins by ChEP confirmed that the core cohesin subunits (SMC1/3 and Rad21) were depleted from chromatin (Fig S2A). In contrast, amounts of other chromatin-associated key scaffold proteins, including condensins, TopoIIα, and KIF4A, were similar to those of control cells (Fig. S2C) and the proteins showed a normal concentration along the axis of each sister chromatid (Fig. S7) (*45*). Importantly, cohesin depletion in late G_2_ had no effect on the kinetics of mitotic entry, as defined by the timely occurrence of NEB (Fig S1D).

We performed Hi-C with control and SMC3-depleted cells during mitotic entry (Fig. 4A, 4B). Consistent with previously reported results in mammalian cells, G_2_-arrested DT40 cells depleted of SMC3 lost cohesin-mediated features (TADs, dots, and stripes), and exhibited much stronger compartmentalization (*46*, *47*)(Fig. 4B, left panel; 5B, middle panels). Global chromatin organization can be characterized by the contact frequency (*P*(*s*)) as a function of genomic separation (*s*). In G_2_, *P*(*s*) changed upon SMC3 depletion: the characteristic “shoulder” at ∼100 kb indicative of the presence of cohesin-mediated loops (*51*, *52*)) was lost. Furthermore, the slope of *P*(*s*)∼*s*^−1^ reveals the fractal globule folding of chromatin evident for more than two decades of genomic separation (*s*= 10^4^-10^7^ bp) (Fig. 4C, Fig. S11B).

**Fig. 4.**
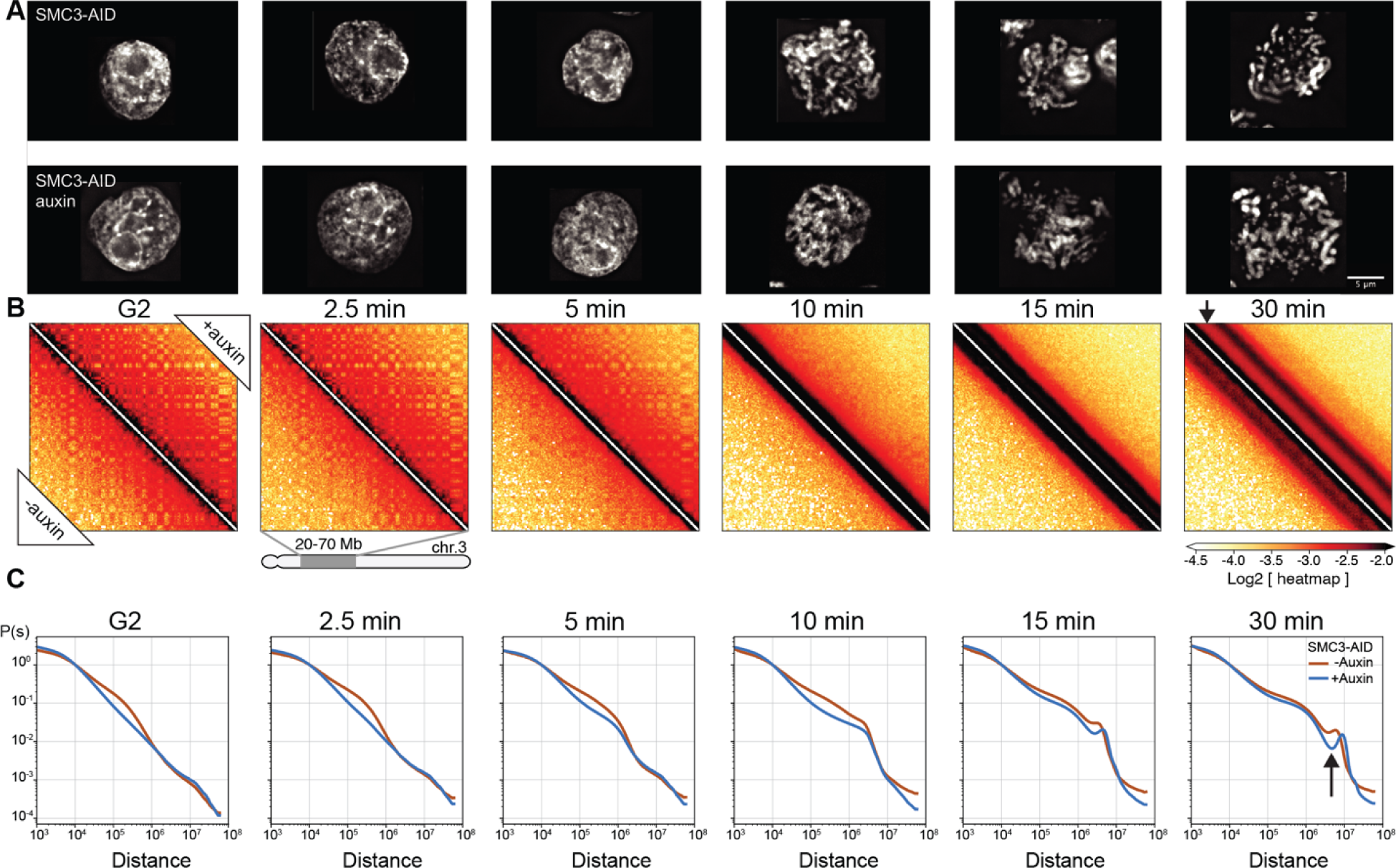
Cohesin impedes spiralization of mitotic chromosomes. (**A**) Representative images of DAPI-stained SMC3-AID cells used to prepare Hi-C samples. Top row: cells from control culture (no auxin treatment). Bottom row: cells from culture treated with auxin during the G_2_ block and release into mitosis. Scale bar = 5 µm. (**B**) Hi-C contact maps for SMC3-AID CDK1^as^ cells treated as in panel A. Chromosome 3, position 20-70 Mb is shown. Compartmentalization (checkerboard pattern in Hi-C maps) was stronger in G_2_ cells, and the second diagonal band appeared sharper, and positioned at larger genomic distance in prometaphase, in SMC3-depleted CDK1^as^ cells (black arrow) compared to those of non-depleted control cells. (**C**) Quantification of Hi-C data shown in panel B: contact frequency *P* plotted as a function of genomic separation (*s*). *P*(*s*) curves reveal position and prominence of the second diagonal band visible in Hi-C maps (arrow).

**Fig. 5.**
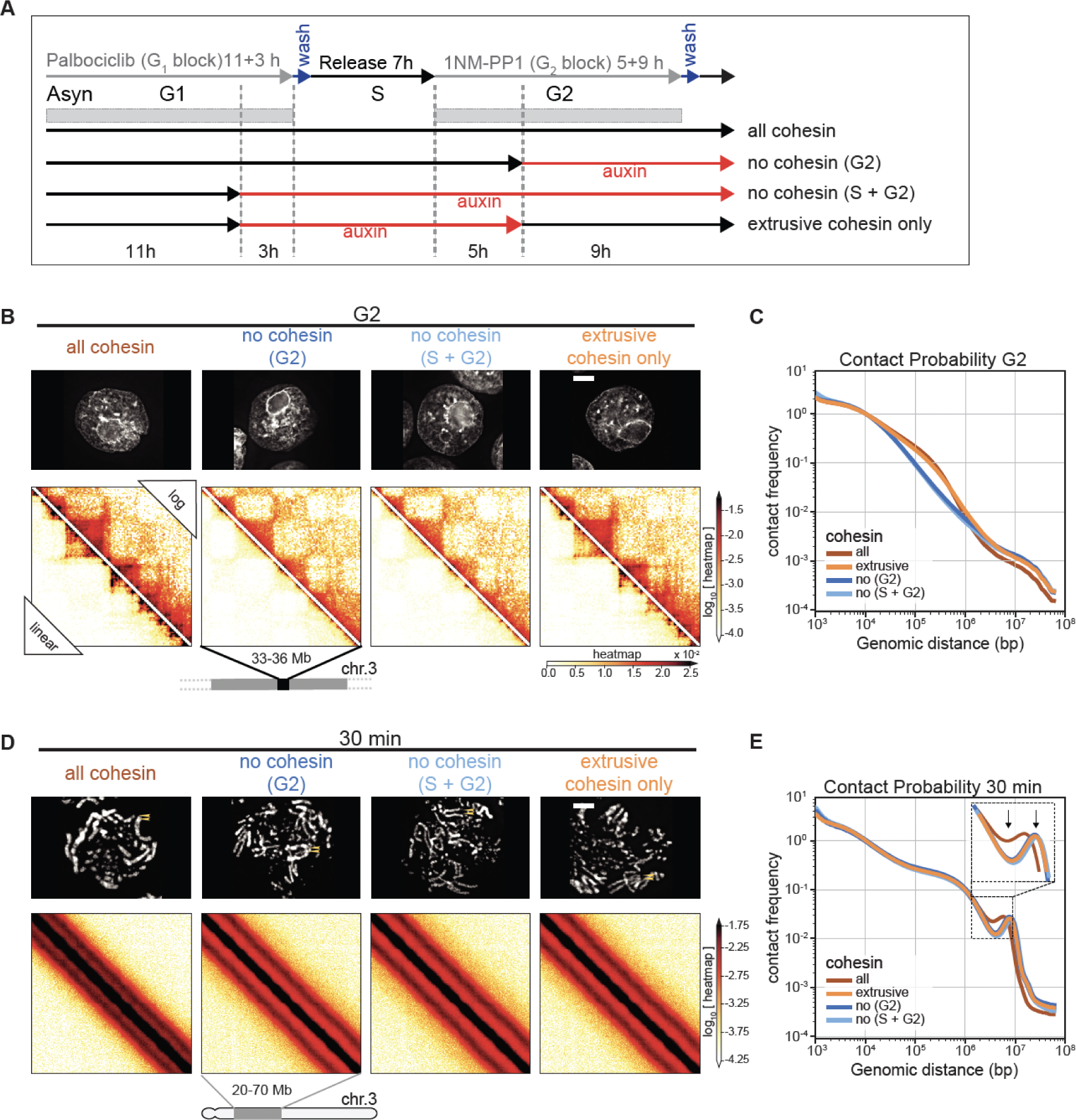
Cohesive and not extrusive cohesin impedes spiralization of mitotic chromosomes. (**A**) Double synchronization procedure for SMC3-AID CDK1^as^ cells. Cells were collected and cross-linked in G_2_ or at 5, 15, 30 minutes after release from G_2_ block. FACS-sorted cells were used for subsequent analysis. GFP-positive cells (labeled “all cohesin” and “extrusive cohesin only”) contained those respective SMC3 populations. In GFP -negative cells (“no cohesin, G_2_” and “no cohesin, S + G_2_”) SMC3 was depleted in the indicated cell cycle phases. (**B**, **D**) Images of DAPI-stained SMC3-AID cells used to prepare Hi-C samples and corresponding Hi-C interaction maps at G_2_ (B) and t= 30 min (D). Scale bar = 5 µm. Hi-C interaction maps are shown in log (top right) and linear scales (bottom left). (**C**, **E**) Contact frequency *P*(*s*) vs. genomic separation (*s*) for maps shown in B and D. Inset in (**E**) shows the magnified view of the region of *P*(*s*) corresponding to the second diagonal band visible in Hi-C interaction maps. Arrows point the difference in the position and prominence of the second diagonal band.

By prometaphase, striking differences in Hi-C interaction maps revealed that cohesin modifies the chromatin organization of the condensed chromosomes. The maps from wild type prometaphase cells revealed a second diagonal, reflecting a helical loop arrangement (*6*, *53*) (Fig 4C, lower left of each panel). In cohesin-depleted cells, this feature changed in two ways (Fig 4C, upper right of each panel). First, the second diagonal was substantially more prominent. This confirms that elevated interactions along the second diagonal are not due to inter-sister interactions and rather reflect helical organization of individual chromatids. Second, the position of the second diagonal moved to larger genomic distances (from ∼6 to ∼8 Mb) (Fig. 4C). Together, both observations suggest that the helical pattern is altered by the presence of cohesin-mediated interactions.

Cohesin appears to “squeeze” mitotic chromosomes, causing both a lengthening and lateral compression of sister chromatids that can be observed by light microscopy. In control DT40 cells, sister chromatid arms remained tightly paired during early mitosis and individual sisters separate only just before anaphase onset. Consistently, SMC3-depleted sister chromatids were already separated at the earliest stages of prometaphase (Fig 4A, t= 10 min), yet remained loosely associated, probably as a result of residual catenations (*54–57*). SMC3-depleted late prometaphase chromosomes (t= 30 min) were both wider and shorter than chromosomes in control cells (Fig. 4A, 6C-E), consistent with the increased size of each helical turn observed in Hi-C (Fig. 4C). Wild-type chromosomes with two tightly cohesed sister chromatids had an overall width of 1.1 µm, consistent with a sister chromatid width of 0.55 µm. Paradoxically, the diameter of individual chromatids is actually 0.8 µm when measured in the absence of cohesin (Fig. 3C, 6C). Thus, as cohesive cohesin holds sister chromatids together, it compresses the space between them by about 0.3-0.5 µm.

**Fig. 6.**
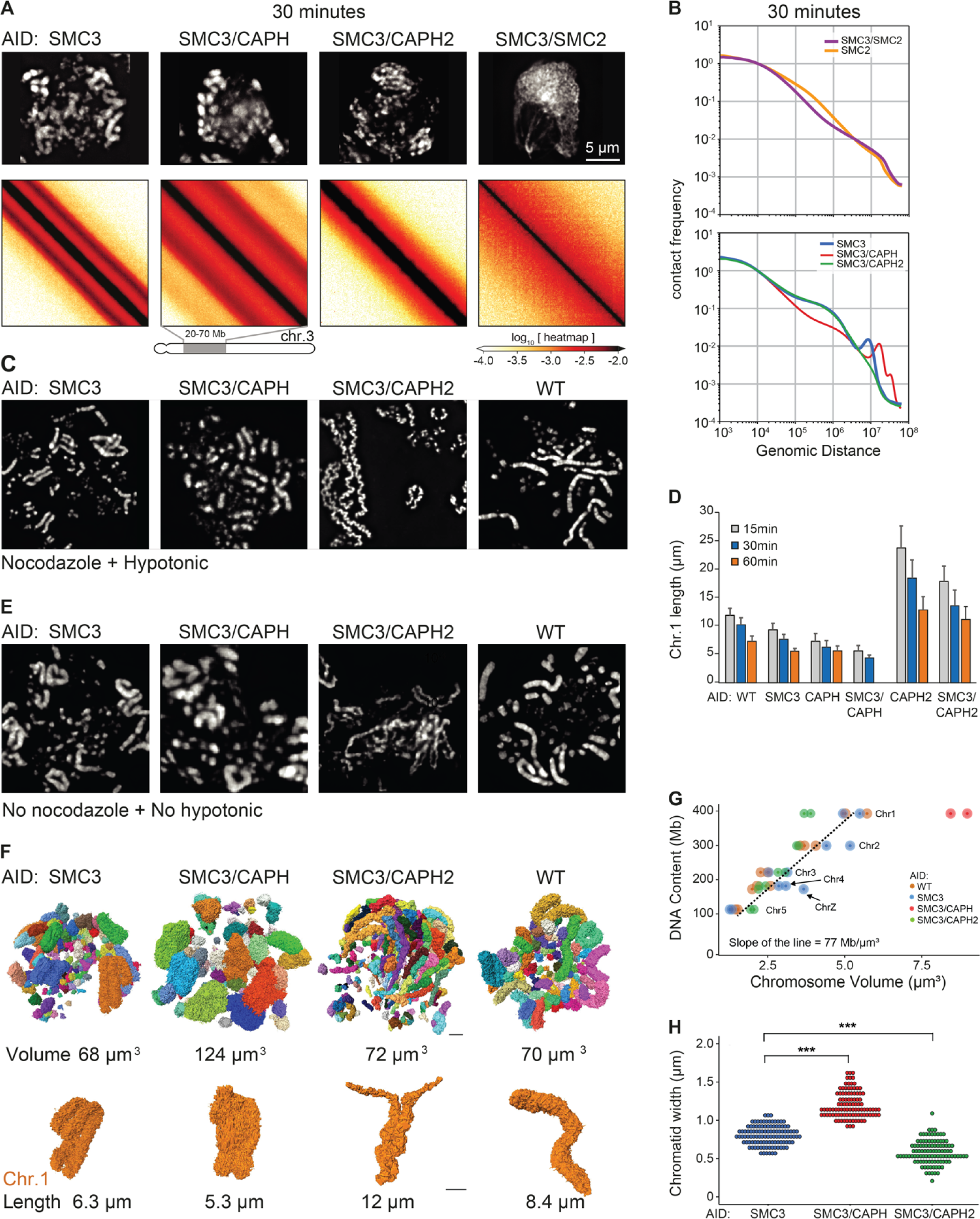
Structure of chromosomes assembled by single SMC complexes. (**A**) SMC3-AID, SMC3-AID/CAP-H-AID, SMC3-AID/CAP-H2-AID, SMC3-AID/SMC2-AID cells treated with auxin during G_2_ arrest were released into mitosis and DAPI-stained cells imaged at t= 30 min (prometaphase). Scale bar = 5 µm. Hi-C was performed on the same cultures at t= 30 min. Hi-C interaction maps for a portion of chromosome 3 is shown for each cell line. (**B**) Plots showing contact frequency *P* as a function of genomic separation (*s*) for Hi-C data shown in A, as well as for SMC2-depleted cells. (**C**) Chromosome spreads of DAPI-stained WT and SMC3-AID, SMC3-AID/CAP-H-AID, SMC3-AID/CAP-H2-AID cells plus auxin. Cells in G_2_ block were treated with 0.5 µg/ml nocodazole for 30 minutes prior to release into mitosis for an additional 30 min. Cells were harvested and hypotonically swollen with 75 mM KCl for 10 minutes prior to ice-cold methanol-acetic acid (3:1) fixation and spreading. (**D**) Length of the longest chromosome (Chr 1) of each cell line treated with auxin in the previous G_2_ measured using cells processed at t= 30 min post release from G_2_ (prometaphase) as shown in panel C. n ≥16 chromosomes or chromatids measured in 3 independent experiments. Average and standard deviation are shown. (**E**) As in (C) but cells were rinsed with PBS prior to ice-cold methanol-acetic acid (3:1) fixation and spreading (no nocodazole, no hypotonic treatment). (**F**) 3D Reconstruction of prometaphase chromosomes of WT, SMC3-AID, SMC3-AID/CAP-H-AID and SMC3-AID/CAP-H2-AID cells treated with auxin in G_2_, and then released into mitosis (t= 30 minutes). Each image represents a 3D reconstruction of an entire mitotic cell obtained by SBF-SEM. Each chromosome is represented as a different color. Chromosome 1 of each cell line is shown in orange. The total volume of the chromosomes and the length of chromosome 1 are indicated. Scale bar = 2 μm. (**G**) Correlation between DNA content vs. chromosome volume, derived from images shown in panel G. Chromosomes 1-5 and Z of each cell line are annotated in the graph. The slope of the line is calculated for WT, SMC3-AID, and SMC3-AID/CAP-H2-AID. (**H**) Chromatid width quantification of large chromosomes in each mutant. Each point represents a measurement, and 10 measurements were taken per chromosome. n= 10 chromosomes, *** p<0.001.

Together, these data show that cohesin has a previously unappreciated role in modulating condensin-driven mitotic chromosome architecture.

### Cohesive, but not extrusive cohesin, shapes mitotic chromosomes

Given the unexpected role of cohesin in mitotic chromosome architecture, we next sought to determine whether this role is carried out by cohesive or extrusive forms of cohesin, or both. In yeast, cohesion is only established during S-phase (*50*). Hypothesizing that this is also the case in DT40 cells (as we confirm below), we established a double synchronization protocol to obtain G_2_ and mitotic cells with chromosomes carrying both cohesive and extrusive cohesin, no cohesin, or only extrusive cohesin (Fig. 5A). We blocked SMC3-AID/CDK1^as^ cells at the G_1_/S boundary with the CDK4/6 inhibitor palbociclib for 11 hours (one cell cycle) followed by an additional 3 hours in the presence or absence of auxin. Subsequent palbociclib washout yielded two cultures of S-phase cells; one with cohesin and one without. Subsequent addition of 1NM-PP1 blocked all cells at the G_2_/M boundary. Addition or removal of auxin during this G_2_ arrest and subsequent release into mitosis yielded four cell populations with distinct cohesin states (Fig. 5A):

1. Cohesin present during S and subsequent mitosis;
2. Cohesin present during S and absent during subsequent G_2_ / mitosis;
3. Cohesin absent during S and during subsequent G_2_ / mitosis;
4. Cohesin absent during S and restored during subsequent G_2_ / mitosis;

All 4 cultures progressed through S phase, entered G_2_ with essentially identical kinetics, and efficiently and synchronously entered mitosis, forming rod-shaped chromatids after 1NM-PP1 washout (see Fig. 5D, S5D). This confirms earlier observations that cohesin is not essential for normal S phase progression, entry into mitosis or for the formation of rod-shaped chromosomes.

Control cultures 1-3 all behaved as predicted (Fig. 4). In G_2_, TADs and dots (CTCF-CTCF loops) were readily observed in Hi-C maps in the presence of cohesin, and absent when cohesin was depleted (Fig. 5B,C). Hi-C confirmed that extruding cohesin was indeed restored onto chromatin after auxin wash-off in G_2_, as the chromosomes of condition 4 had robust TADs and dots (Fig. 5B). As expected from prior published studies with yeast (*50*), cells in condition 4 (no cohesin present during replication) failed to establish sister chromatid cohesion during G_2_ / mitosis when cohesin was restored after S-phase (Fig. 5D,E, S5D). In condition 4, TADs and dots were restored when cohesin was expressed during G_2_ (Fig. 5B). Thus, extrusive cohesin, rather than cohesive cohesin, is important for formation of TADs, dots and stripes in G_2_.

By late prometaphase (at t= 30 min), Hi-C data obtained from the three cultures that lack cohesive cohesin (cultures 2, 3, 4) were indistinguishable regardless of whether extrusive cohesin was present, and showed the same differences from control cells (culture 1). Specifically, the second diagonal ran at a larger distance compared to control cells (similar to results in Fig. 4). It also appeared much more pronounced with a deeper dip midway between the first and second diagonal (Fig. 5D,E). Similarly, microscopy of these prometaphase chromosomes showed that in all cultures lacking cohesive cohesin, the separated sister chromatids were shorter and wider than chromosomes in control cells, regardless of extrusive cohesin re-loading in G_2_ (Fig S5).

Thus, cohesive, and not extrusive, cohesin alters the helical conformation and dimensions of prometaphase chromosomes, presumably by linking sister chromatids and limiting the ability of each sister to form an independent helical condensin loop array.

### Roles of individual condensin complexes in mitotic chromosome assembly

Cohesin depletion further allowed us to uncover the individual activities of condensin I and II in mitotic chromosome assembly. We therefore created three additional cell lines to disentangle the roles of these individual SMC complexes in chromosome assembly. SMC5/6 complexes were not included here as their depletion in mitosis was reported to have no effect on mitotic chromosome structure (*58*).

1. Condensin II only - SMC3-AID/CAP-H-AID cells allow co-depletion of cohesin and condensin I;
2. Condensin I only - SMC3-AID/CAP-H2-AID cells allow co-depletion of cohesin and condensin II;
3. No cohesin or condensin - SMC3-AID/SMC2-AID cells allow co-depletion of cohesin and both condensins.
4. Condensin I & II, no cohesin - SMC3-AID cells, described above, provided a baseline with both condensin I and II active, and cohesin depleted.

Previous studies had also examined cell lines lacking condensin I, II, but all in the presence of cohesin (*6*, *37*, *59–62*). Here we report important additional insights from cell lines expressing only a single SMC complex. For all cell lines, we performed light and electron microscopy and Hi-C analysis of chromosomes from G_2_ to mid-mitosis.

In G_2_, microscopy (DNA staining) and Hi-C data for all the above cell lines lacking cohesin, were indistinguishable from one another (Fig. S6). Hi-C maps showed loss of cohesin-mediated features (TADs, dots, and stripes), P(s) plots lacked the characteristic shoulder at s∼100 kb that normally results from the presence of cohesin-mediated loops (Fig. S11). Thus, these observations confirm and extend earlier data indicating that condensin I and II lack major roles in G_2_ chromatin organization (*6*, *63*).

In contrast, mitotic chromosome morphologies and Hi-C data obtained with cells after co-depletion of different SMC complexes in G_2_ exhibited striking differences (Fig. 6, A and B). Mitotic chromosomes without any condensins lacked a compact rod-shaped morphology and, at late prometaphase, appeared clumped together with the centromeric chromatin often pulled out by the force of kinetochore-microtubule interactions (*40*). This morphology was unaltered by the presence or absence of cohesin (e.g. SMC2-AID versus SMC3-AID/SMC2-AID). Hi-C analysis of SMC2-AID cells and SMC3-AID/SMC2-AID cells revealed an absence of the second diagonal band in prometaphase (Fig. 6B).

Strikingly, the contact frequency *P*(*s*) curve for SMC2-AID cells (in the presence of cohesin, but without condensin) shows the presence of cohesin-mediated loops, seen as a characteristic shoulder at s∼ 100 kb. This declines gradually, but persists deep into prometaphase (t= 30 min) (Fig. 6B). In contrast, SMC3-AID/SMC2-AID cells (no cohesin, no condensin) lack this feature. These data show that cohesin-mediated loops persist in the absence of condensin. The eventual loss of TADs and dots in prometaphase in the absence of condensin (see Fig. 2D) can then be explained by random relocalization of remaining cohesin loops, presumably because of loss of CTCF binding (see Discussion).

Next, we examined prometaphase chromosomes formed by condensin I only or condensin II only. Chromosomes formed by condensin I have been reported to be long and “wiggly” compared to wild-type (*46*, *64*). This phenotype was exaggerated in chromosome spreads when cohesin was also absent, with the chromatids forming wiggly fibers of relatively constant diameter (Fig. 6C - SMC3-AID/CAPH2-AID). The chromosome morphology was dramatically different under more native conditions: chromosomes formed by condensin I only when fixed without nocodazole or hypotonic treatment lacked the “wiggles” and were disorganized with highly variable width (Fig. 6E), resembling chromosomes in 3-dimensional reconstructions of whole cells imaged by electron microscopy (Fig. 6F). Thus, (and often overlooked in other studies) the structural parameters of chromosomes observed by traditional microscopy methods are extremely sensitive to the exact protocols used for sample preparation. Despite this variability, we note that chromosomes assembled by condensin I only are reproducibly shorter than the same chromosomes assembled in the presence of cohesin (Fig. 6D, Fig. S7A).

Chromosomes formed by condensin II only without cohesin were rod-shaped and exhibited a relatively “dumpy” and fragile appearance by light microscopy (SMC3-AID/CAPH-AID in Fig. 6A,C,E). As previously described by others (*37*, *59*, *60*), in chromosome spreads without condensin I, chromosomes tended to be shorter and fatter than chromosomes with both condensins (CAPH-AID vs WT and SMC3-AID/CAPH-AID vs SMC3-AID in Fig. 6D). Consistently, the additional removal of cohesin made chromosomes even shorter and less well defined in shape (CAPH-AID vs SMC3-AID/CAPH-AID in Fig. 6D, S7A).

The lack of cohesin also exaggerated differences in the Hi-C data for condensin I-only and condensin II-only chromosomes. Hi-C data from condensin I -only chromosomes lack the second diagonal, indicating the absence of the helical organization (Fig. 6A, S11D). This is consistent with our previous conclusion that condensin II is required to form a helical array of chromatin loops (*6*). Hi-C data indicate that chromosomes with only condensin I fold into an array of ∼100 kb loops reflected by the shoulder on the *P*(*s*) curve for s∼100 kb (Fig. 6B, S11D). This is followed by a region of steady decay between 2-8 Mb with a slope of −1.5, indicative of the array adopting a random walk (*65*). This behavior is consistent with the microscopy that reveals long wiggly chromosomes in native spreads (Fig. 6A,E,F).

The unremarkable morphology of condensin II-only chromosomes by light microscopy (Fig. 6A,C,E) sharply contrasted with astounding Hi-C maps in which the second diagonal band was flanked by a robust third diagonal, and a fainter 4^th^ diagonal band. The position of the second diagonal appeared around 16 Mb, while for chromosomes formed by condensin I and II the second diagonal appeared at around 8 Mb. The re-location of the second diagonal band to larger genomic distances for condensin II-only chromosomes revealed that each helical gyre contains more DNA, consistent with the shorter and wider morphology of these chromosomes, as compared to the same chromosomes in the presence of cohesin (Fig. 6D,H).

The second diagonal in Hi-C has been explained by interactions between adjacent gyres of chromatids in a helical configuration (*6*). Remarkably, the third and fourth diagonals were positioned at multiples of 16 Mb: 32 and ∼40-50 Mb, suggesting that chromatin in one gyre may interact with chromatin two or three gyres above or below. Careful inspection revealed that a very weak third diagonal band was also visible in Hi-C data from cells lacking condensin I but containing both condensin II and cohesin (Fig. S11E). This strongly suggests that the additional diagonals are caused by condensin II-mediated chromatin loops, and that the presence of cohesin weakens these helical features in Hi-C analysis.

### Measurement of chromosome parameters for polymer modeling

We used polymer modeling to gain mechanistic insights into the roles of individual SMC complexes in mitotic chromosome folding. In order to create accurate polymer models to explain our Hi-C data, it was necessary to first determine precise volumes and shapes for the various mutant chromosomes studied here. Such measurements cannot be obtained from light microscopy due to (1) variations in chromosome morphology as a result of the spreading protocol, (2) differences in cell fixation and (3) the limited resolution of the light microscope. We used Serial Block Face Scanning Electron Microscopy (SBF-SEM) coupled with a labeling method that selectively enhances the contrast of DNA (Methods).

The total volume of all chromosomes in WT, SMC3-AID, and SMC3-AID/CAPH2-AID cells was similar, with values of 68 µm^3^, 72 µm^3^, and 70 µm^3^, respectively. Chromosomes 1 - 5 and Z can be unambiguously identified in WT and SMC3-AID cells. Knowing the volume and DNA content of each yields an average DNA density of 77 Mb/µm^3^ (Fig. 6G, Movies S1-S8). However, the fuzzy-looking chromosomes in SMC3-AID/CAPH-AID mutants had a significantly larger volume of 124 µm^3^ (Fig. 6F), corresponding to an average DNA density of 44 Mb/µm^3^ (Fig. 6G).

The combination of light and electron microscopy measurements described here together with Hi-C observations allowed us to build accurate polymer models of the organization of chromatin within individual chromatids.

### Structural models for the roles of condensins in mitotic chromosome formation

We modeled the folding of chromatids formed by individual condensin complexes, using our SMC3-AID/CAPH-AID and SMC3-AID/CAPH2-AID mutant cell lines. This eliminated complications arising from interactions between sister chromatids. We focused on the 30 minute time point (late prometaphase/metaphase) as it represents mature mitotic chromatids.

Our model was built on four simple principles (Fig. 7A):

i. The nucleosome fiber is modeled as a 100 Mb chain of connected 10 nm/200 bp beads that move by random Brownian motion.
ii. This fiber is folded by condensins into an array of consecutive loops separated by small gaps.
iii. The resulting bottle-brush of chromatin loops is tightly packed into a cylindrical chromatid body that effectively models condensin-independent condensation of mitotic chromatin (*40*, *66*, *67*).
iv. This chain of loops is weakly “nudged” to follow a helical path by pinning two ends to the opposite caps of the cylinder and fixing the number of turns that it makes around the axis of the cylinder.

**Fig. 7.**
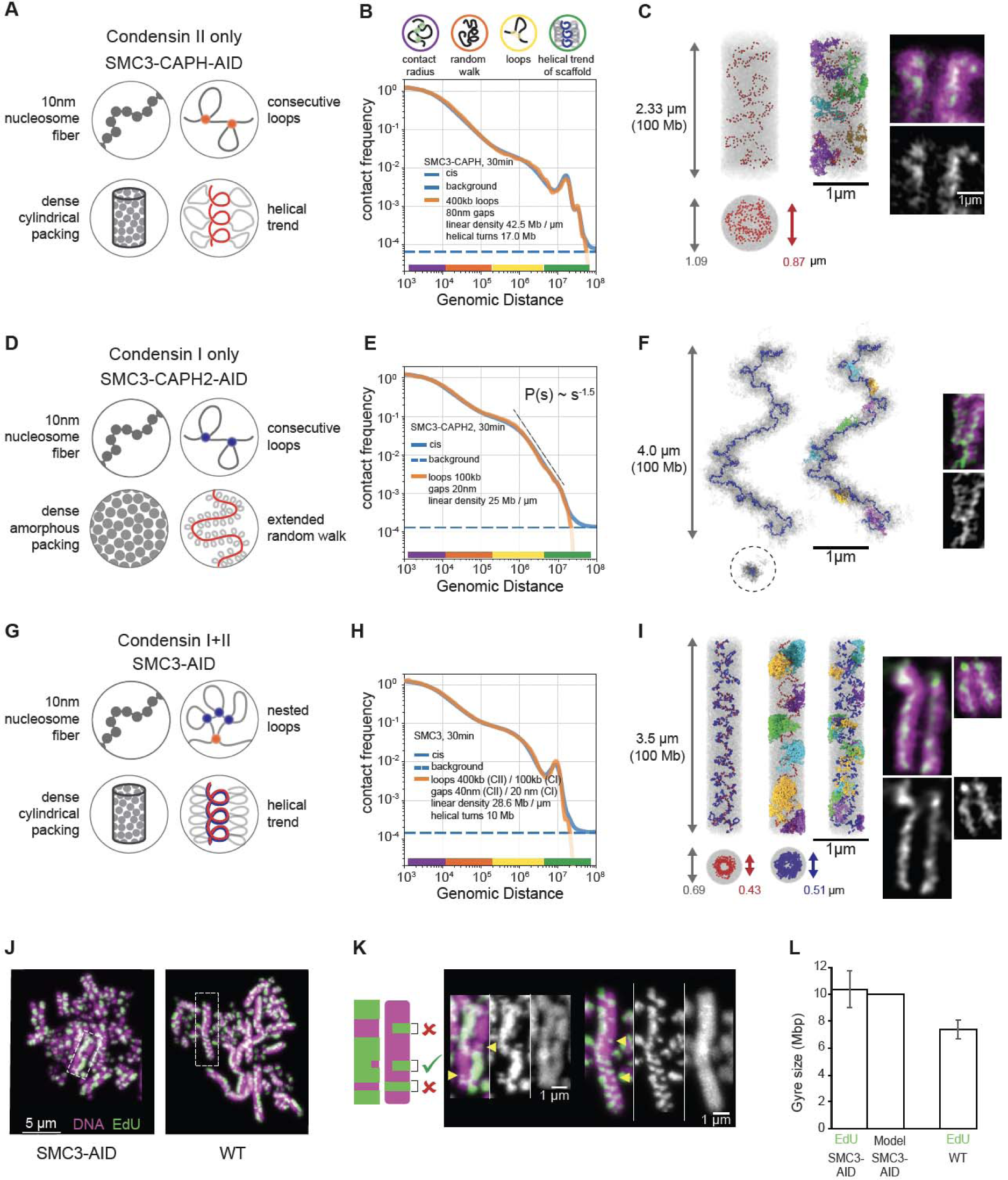
Polymer modeling reveals the internal organization of chromosomes built by single condensin complexes. (A-C) Model of SMC3-AID/CAPH-AID (condensin II-only) prometaphase (t=30 min) chromosomes. (**A**) The four modeling assumptions. (**B**) Contact frequency *P* as a function of genomic separation *s* for the best-fitting model (orange line, model parameters are listed in the plot), in comparison to experimental data (blue line). Dotted line indicates upper limit of the background interaction frequency in experiments, estimated from the average inter-chromosomal interaction frequency. The colored circles (top) and color bars (bottom) indicate levels of genomic organization and their characteristic sizes in base pairs. (**C**) Left: Simulated chromosome conformation in one modeling replicate in longitudinal projection (top) and a cross-section (bottom). DNA is shown in gray, condensins II shown as red spheres. Gray and red arrows indicate diameter of the chromatid and the condensin scaffold respectively. Middle: Same, but with a few selected loops stained in different colors. Right: A microscopy image of SMC3-AID/CAP-H-AID/Halo-CAP-H2 chromosome. Halo-JFX549 (green) is added to the medium >30 min prior to 1NM-PP1 washout to stain Halo-tagged proteins. Cells were treated with 1NM-PP1 for 13 h and fixed with formaldehyde 30 minutes after 1NM-PP1 washout, and DNA was stained with Hoechst (magenta). Scale bar = 1 µm. (**D-F**) Same as (A-C) but for SMC3-AID/CAPH2-AID (condensin I-only) prometaphase chromosomes. In (F), the two left images show positions of condensins I as blue spheres. In the middle image, a few selected loops are colored. Right: Microscopy images of SMC3-AID/CAP-H2-AID/SMC2-Halo chromosomes, stained with DAPI and Halo. (**G-I**) Same as (A-C) but for SMC3-AID (condensin I+II) chromosomes. In (I), SMC3-AID/SMC2-Halo (left) shows DNA plus all condensins, SMC3-AID/Halo-CAP-H2 (right) shows DNA plus only condensin II. (**J**) SMC3-AID and WT chromosome spreads in which EdU is incorporated into one sister chromatid after two cycles of DNA replication. The harlequin appearance is caused by sister-chromatid exchanges. EdU (green), DNA (magenta). Scale bar = 5 µm. (**K**) Enlarged images from (J). EdU (green), DNA (magenta). Scale bar = 1 µm. Cartoon shows the criteria used to select partial exchanges to measure the height of an EdU-labeled gyre. Arrow heads indicate gyres matching criteria (**L**) Size of gyre calculated from EdU height in wild type and SMC3-AID (cohesin depleted) sister-chromatid exchanges and the average gyre size used for modeling. n=4, total 92 measurements (SMC3-AID), n=3, total 81 measurements (WT).

The resulting models for chromatids formed by a single condensin complex have three key free parameters (Fig. S9A): (1) average loop size (bp), (2) gap size between condensin complexes (nm), and (3) the linear density of chromatin along the cylinder axis (Mb/µm). Two other parameters (Fig. S9A) were determined from experimental data directly: (1) the length of a turn in Mb (evident from the position of the second diagonal in Hi-C maps), and (2) the volume density of chromatin (measured by SBF-SEM, Fig. 6G). This framework produces a class of individual structural models, whose morphology and contact frequency *P*(*s*) curves are sensitive to changes of each parameter (Fig. S9A). We explored the space of these parameters (see Methods, Fig. S9B), matching the *P*(*s*) curve predicted by each model against that from Hi-C data (Methods).

For chromosomes formed by condensin II only (SMC3-AID/CAPH-AID cells), our best-fit model reproduced the *P*(*s*) from Hi-C across four and a half decades of genomic separation (Fig. 7B). In the model, condensins II formed 400 kb loops, with adjacent loop anchors separated by 80 nm gaps, and organized into an irregular helix (Fig. 7C) with a pitch of 400 nm containing 17 Mb. Individual chromosomes generated by this model reproduce Hi-C equally well, yet clearly differ in the position of the scaffold subunits and the folding of individual loops, reinforcing the notion that there is no single structure of mitotic chromosomes (Fig. S9C,D).

Intriguingly, *P*(*s*) predicted from the best-fit models reproduced both the third diagonal at 34 Mb and a weak fourth diagonal at ∼40-50 Mb that were also observed in Hi-C. Inspection of the model indicated that the third and the fourth diagonals represented contacts between large loops separated by two and three helical turns (Fig. S9E). This explains why these extra diagonals run at multiples of the distance of the second diagonal. Thus, helical bottlebrush chromosomes can naturally produce multiple periodic diagonals visible in Hi-C, which were not explicitly imposed in our models.

We next modeled chromatids formed by only condensin I (SMC3-AID/CAPH2-AID cells). These chromatids do not have a well-defined cylindrical (rod shape) morphology, but rather form “wiggling noodles” with variable width (Fig. 6C,E,F). Consistently, their long-distance (>1 Mb) Hi-C contacts show no periodic patterns (i.e., additional diagonals) and the contact frequency *P*(*s*) curve instead decays as s^−1.5^ for 2 Mb < s < 8 Mb, suggesting that the bottlebrush of loops follows a random-walk trajectory at these length scales. Guided by these observations, we did not impose a cylindrical constraint (iii, above) and instead allowed the chromatid to fold into a stretched random walk with a specified end-to-end linear distance (Fig. S10A). To model non-cylindrical chromosomes with variable width, we replaced the cylindrical constraint with periodic boundary conditions. This way we could still impose the high chromatin density as measured by SBF-SEM, but not make any specific assumption on the shape of the chromatid (Fig. 7D).

The best-fit model accurately reproduced the experimental Hi-C data across over four decades of genomic separation (Fig. 7E, S10B, Methods) and was consistent with microscopy images. In this model, condensin I folds chromosomes into bottle-brushes of ∼100 kb loops, separated by gaps of 20 nm (Fig. S10B). At the scale of multiple loops, this bottle-brush spontaneously folded into a random walk without spiralization and with a 3D end-to-end separation of 4 µm for a 100Mb chromatid (Fig. 7F, S10B,C).

These principles of chromosome folding by condensins II and I individually allowed us to reconstruct the internal structure of chromatids folded by the combined action of the two condensins in the absence of cohesin (SMC3-AID). This model included the same assumptions (i)-(iv) as the condensin II-only model (above), yet additionally, condensins I formed a second layer of shorter loops, nested into longer condensin II-mediated loops as a result of collisions between condensin complexes (Fig. 7G-I, Methods) (*6*). Surprisingly, with only minor changes in the gap size, combining the parameters obtained from the two independent models for each condensin complex described above yielded a model that accurately reproduced experimental Hi-C from cells expressing both condensins (SMC3-AID) across more than four decades of genomic separation (Fig. 7H, S10D, Methods). Furthermore, these loop sizes (400 kb for condensin II, 100 kb for condensin I) were consistent with the results of quantitative proteomics (Fig. 1D; 2-3 condensins II and ∼10 condensins I per Mb). This suggests that the two condensin complexes function additively despite acting on the same chromatin substrate at the same time.

Two approaches allowed independent validation of the structural predictions of the models for overall chromosome shape and internal structure. Both came from the agreement between model predictions and light and electron microscopy measurements that were not used to construct the models. Firstly, the chromatid width, length, and 3D distance between chromatid ends observed in chromosome spreads and in intact cells correspond closely to the corresponding distances predicted by the models (Table S3). Secondly, it has been reported that gyre size can be estimated by measuring the dimensions of a class of sister chromatid exchange events that can be detected by EdU labeling ((*53*) and Methods, Fig. 7J-K). Using this approach allowed us to obtain independent measurements of gyre sizes. Remarkably, the height of these partial exchange events measured experimentally corresponds nearly exactly to the pitch of the helix in the model calculations for SMC3-depleted cells and to the genomic spacing of the second diagonal in Hi-C of SMC3-depleted and wild-type cells (Fig. 7J-K, Fig. S8, Tables S4 and S5).

Together our new models of chromatin folding within individual chromatids gave several novel insights: (i) the condensin scaffold is discontinuous with gaps between loops; (ii) longer chromatids formed by condensin I are best reproduced by a randomly folded and weakly stretched bottlebrush; (iii) the strong second diagonals detected in Hi-C experiments on cell populations can be explained by a relatively weak and irregular condensin II-mediated spiraling of individual chromosomes that is consistent with microscopy data where a regular spiral is not detected; and (iv) condensin I and II complexes act additively and in parallel.

## DISCUSSION

During prophase, two types of cohesin and two types of condensin act on chromosomes. Our study reveals how collisions between these complexes are resolved, defining three “rules of engagement” when an actively extruding condensin complex runs into another SMC complex: bypassing, collision-facilitated removal, and stopping/blocking. These rules can be mediated by steric interactions, specific protein-protein interactions (as is the case for CTCF-cohesin interactions), and/or the physical state of the chromatin template itself (e.g. tension) (*68*). Application of such rules explains how the local action of SMC complexes and their interactions leads to the formation of condensed mitotic chromatids that are cohesed via interactions between their loops. Furthermore, as we discuss below, our temporal analysis in DT40 cells entering highly synchronous mitosis also allows us for the first time to calculate the speed of loop extrusion by condensin complexes in living vertebrate cells.

### Rules of engagement when Condensin and Cohesin encounter each other

#### Rule 1: Condensins bypass cohesive cohesins

Our data provide strong evidence for the ability of condensins to bypass cohesive cohesins in vivo. Cohesins that connect arms of sister chromatids end up at the tips of condensin-extruded loops by t= 30 min into prometaphase. Since those cohesins were loaded during S-phase, before condensin-mediated extrusion begins in prophase, they must be bypassed to end up at the tips of the condensin loops, as shown by our simulations. If condensins failed to bypass cohesive cohesins but instead either pushed or stalled on them, then cohesins would accumulate at the bases of condensin loops. The result would be a structure with two sister chromatids connected at the bases of the condensin loops by cohesins, thus forming a single axis, contrary to what is observed. However, a single axis could form transiently if condensins pause before bypassing cohesive cohesins.

The ability of loop-extruding SMCs to bypass each other (*9*) or large DNA-bound obstacles (*68*) has been observed in single-molecule experiments, and bypassing of bacterial SMCs was revealed using engineered bacterial chromosomes (*69*). In contrast, extrusive cohesins appear not to bypass cohesive cohesins in *S. cerevisae* (*70*). Here we demonstrate that vertebrate condensins can bypass other SMC complexes, and obstacles they tether. For example, when condensins bypass cohesive cohesin, they are effectively bypassing the entire other sister chromatid.

#### Rule 2. Condensins remove extrusive cohesins

In contrast to the situation with cohesive cohesin, encounters between condensin and extrusive cohesin are apparently resolved by removal of cohesin (e.g., the prophase pathway (*24*, *25*)). Our Hi-C data indicate that cohesin loops are normally removed by late prometaphase. At 30 minutes, *P*(*s*) obtained for CAPH-AID cells (which lack condensin I-mediated loops that would obscure the detection of cohesin loops) is identical to the *P*(*s*) of SMC3-CAPH-AID (which also lack cohesins) at s < 1Mb (Fig. S12). This loss of cohesin-mediated loops is condensin II-dependent, since 100 kb cohesin loops are readily seen in the *P*(*s*) of prometaphase SMC2-AID cells at 30 minutes (Fig. S12, black arrow).

The cohesin loops that remain until late prometaphase in the absence of condensins (SMC2-AID cells) are not positioned at CTCF sites, since we do not see an enrichment of CTCF-CTCF interactions (dot score at 30 minutes, Fig.2D, S4C). The most likely explanation is that another condensin-independent pathway removes positioning cues for cohesin loops. This background process is likely driven by progressive unloading of CTCF (*71*). The dominant rapid condensin-mediated removal of cohesin acts in parallel with this background process.

Modeling suggests that condensin removes cohesin loops by either pushing cohesin aside (i.e., hijacking DNA from cohesin loops into condensin loops) or by facilitating unloading of cohesin, which does not rebind. Neither pushing of roadblocks nor facilitated unloading of other SMCs by condensins have been observed in in vitro single-molecule experiments (*68*). Furthermore, it appears that pushing would require some force, and condensins are relatively weak motors (*72*, *73*), making pushing a less likely scenario. In support of unloading, our ChEP data show that cohesin is lost more rapidly during prophase in the presence of condensin. It thus appears that two processes disassemble interphase cohesin-mediated structures: condensin-mediated unloading of cohesins, and a condensin-independent loss of CTCF.

#### Rule 3. When condensins encounter one another, they stall

Although our experiments show clearly that condensin II must be able to bypass cohesive cohesin, two lines of evidence argue that condensin II complexes do not bypass each other and instead stall when they encounter each other. First, the average loop size in condensin II-only chromosomes stabilizes and stops growing after 5 minutes in prophase (see below), suggesting that they extrude all available chromatin into loops and then stall (Fig. S11C).

Second, microscopy shows that in condensin II-only chromatids, condensin is concentrated in the interior of the chromatid (Fig. 7C). While models with consecutive loops, i.e. non-bypassing condensins, can reproduce such a localized condensin scaffold distribution (Fig. 7C, S7B), models where loops overlap, i.e. extruding condensins freely bypass other condensins, end up with condensins dispersed evenly throughout the chromatid (Fig. S9F-H).

### In vivo estimation of the extrusion speed

Our system with highly synchronous entry into prophase in the presence of a single extruder, alongside models that infer loop sizes from Hi-C data, provides a unique opportunity to measure the dynamics of loop growth in mitosis and estimate the speed of extrusion of individual condensins in vivo.

Characteristic loop sizes can be estimated from the *P*(*s*) curves. While in interphase, where the loop density is low, the loop size corresponds to genomic distance at the peak of the *P*(*s*) derivative (*51*, *52*), its position in the dense mitotic loop array is harder to infer. Our models show that the loop size defined in a model has a characteristic position somewhat to the left of the peak on its *P*(*s*) derivative (Fig. S9A, arrowhead). This allows us to estimate loop sizes for time points for which the full model is not available.

For chromosomes formed only by condensin II, at t= 2.5 min, P(s) reveals a lower density of loops and suggests an average loop size of ∼200-300 kb (Fig. S11C). We thus estimate the speed of extrusion for individual condensin II complexes as 200-300 kb/2.5 min, giving an extrusion speed in vivo of ∼1.3-2 kb/sec. At t= 5 min, the loop size is 400 kb (Fig. S11C), again yielding an extrusion speed of ∼1.3 kb/sec. These values are approximations because we cannot know the exact time when condensin II is activated during reversal of the 1NM-PP1 block. In an independent approach, by measuring the change in loop size between t= 2.5 and 5 minutes in a single time course we arrive at (400 minus 200-300) = 100-200 kb in 2.5 min (i.e., an extrusion speed of 0.5-1.3 kb/sec). This is likely to be an underestimate, as loops likely reached their maximum size between t= 2.5 and 5 minutes.

Microscopy provides a third way to estimate the speed of extrusion by condensin II. By t∼10 minutes, chromosomes in SMC3-AID/CAPH-AID cells acquire the flexible rod-like shape that is characteristic of a dense loop array (Fig. S6C). Simulations show that this shape requires adjacent extruders to meet each other with most of the chromatin extruded into loops. Theory shows that to close most gaps, condensins need to extrude ∼4-5 times the average loop size (Methods). Using the average loop size from the *P*(*s*) at t= 10 minutes (400 kb), we calculate an extrusion speed of 400 kb*(4-5) / 600 sec=2.5-3 kb/sec.

Altogether, our analysis in SMC3-AID/CAPH-AID cells yields an extrusion velocity of condensin II as 1-3 kb/sec in living DT40 cells.

This approach also allows us to quantify loop extrusion by condensin I. In condensin-I only chromosomes, the *P*(*s*) reveals loop formation between t= 5 and 10 minutes, i.e., even before nuclear envelope breakdown (Fig. S11D). Together with the ChEP data, this argues for the presence of active nuclear condensin I at these early time points (*60*). Comparison of the *P*(*s*) for t= 2.5 minutes (no loops) and 7.5 minutes (∼300-400 kb loops) (Fig. S11D), reveals that these nuclear condensins I extrude loops at ∼1 kb/sec.

We can also directly observe the formation of nested loops by condensin I in the presence of condensin II in SMC3-AID cells. We observe formation of large (400 kb) loops before NEB (by t= 10 minutes, largely by condensin II) (Fig. S11B). The loops then abruptly become smaller (∼100 kb) upon NEB when the bulk of condensin I gets access to chromosomes (Fig. S11B). This drop in the loop size strongly supports a nested loop organization of the mitotic chromosome where each ∼400 kb condensin II loop is split into several ∼100 kb condensin I loops.

### Spiraling dynamics

Analysis of the *P*(*s*) curves at different time points yields information on the period of the chromatid spiral as revealed by the second diagonal, i.e., the first peak on the *P*(*s*) curve (Fig. S9A, S11A,B,D). In wild type cells, the period grows from 4.0 to 6.1 Mb between t= 15 and 30 minutes (∼2.3 kb/sec). This growth of the spiral correlates with an overall shortening of the chromosome (Fig. 6D). In SMC3-depleted cells, the growth of the spiral over this period rises to ∼3.5 kb/sec. For chromosomes built solely by condensin II, the period changes from ∼6.6 Mb to 16.4 Mb between 15 and 30 minutes. This yields the growth rate of 650 kb/min = 10.9 kb/sec over 15 minutes, i.e. almost an order of magnitude higher than the speed of condensin loop extrusion. Moreover, the *P*(*s*) between these two time points shows almost no change in average loop size. Therefore, the growth of the helical spiral, though dependent on condensin II, appears to be driven by processes other than extrusion of the 400 Kb loops that has been completed by the end of prophase. This additional role of condensin II in driving spiraling is restrained by cohesive cohesin and condensin I (Table S5).

In summary, the present study combines genetics, microscopy, proteomics, Hi-C and polymer modeling to define rules of engagement that dictate the outcome when condensins encounter other condensins and extrusive or cohesive cohesins during mitotic chromosome formation. These rules allow chromosomes to transition from largely cohesin-organized interphase chromatin to condensin-compacted rod-shaped paired sister chromatids. Cohesive cohesin tethering loops between sister chromatids limits the ability of the loop array in each chromatid to adopt a more regular helical folding. We find that mitotic chromosomes are disorderly helices with chromatin loops distributed throughout the body of the chromosome and organized by a discontinuous scaffold.

## Acknowledgments

We thank Christos Spanos for help with mass spectrometry analysis, David Kelly and Toni McHugh for help with light microscopy and Martin Waterfall for help with flow cytometry.Sorting of GFP positive and negative cells was performed in the Flow Cytometry facility, Institute of Immunology & Infection Research, Edinburgh with assistance of Dr. Martin Waterfall.

## Funding

Austrian Science Fund (FWF) grant SFB F 8804-B “Meiosis” (AG)

European Union (ERC, CHROMSEG, 101054950) (AAJ)

Howard Hughes Medical Institute (JD)

Medical Research Council (MRC, United Kingdom; MR/X001245/1) (AAJ)

National Human Genome Research Institute grant HG003143 (JD)

National Institutes of Health Common Fund grant DK107980, HG011536 (JD, LM)

National Institute of General Medical Sciences grant GM114190 (LM)

NSF Physics of Living Systems grant 1504942 (LM)

Wellcome grant 107022 (WCE)

Wellcome grant 221044 (WCE).

Wellcome grant 203149 (The Wellcome Centre for Cell Biology)

## Author contributions

KS generated cell lines, established cell synchronization protocols, performed synchronized time courses and performed imaging analyses. JHG and SA performed Hi-C analyses. FSC, AB and IP performed and FSC and NP analyzed electron microscopy. SA and AG performed polymer simulations. IS performed chromatin enriched proteomics analyses using spike-in protein isolated by MAA, BMP and JA. LX and JRP prepared 1NM-PPl. KS, JHG, SA, FCS, IS, LAM, JD, AG, and WCE designed the project, analyzed data, and contributed to writing the manuscript.

## Data and materials availability

Data produced in this paper is part of NCBI BioProject number PRJNA1091327. Hi-C data has been submitted to GEO and will be publicly available upon publication (accession number: GSE262525). EM data has been submitted to the Electron Microscopy Public Image Archive: EMPIAR-11919

## Supplementary Materials

Materials and Methods

Supplementary Text

Figs. S1 to S12

Tables S1 to S7

References (*68–113*)

Movies S1 to S8

