## Supplemental Materials for "Rules of engagement for condensins and cohesins guide mitotic chromosome formation"

MATERIALS AND METHODS:

Cell culture

Cell Culture, transfection to establish stable cell lines, and drug treatments.

Type Chicken DT40 (B-cell lymphoma) cells were cultured in RPMI 1640 medium supplemented with 10% fetal bovine serum and 1% chicken serum in a humidified 39˚C incubator with 5% CO_2_ in air. When specified, PIPES-NaOH (0.5 M, pH 7) was added to the culture medium at final concentration 25 µM.

Stable transfection was achieved by two different methods. For random integration of exogenous DNA, a Biorad electroporator was employed following a previously described protocol (*74*). To insert tags (AID-GFP/Clover, Halo, GFP), the Neon transfection system (Neon1, Thermofisher Scientific) was utilized. Typically, 2-4 million cells suspended in R-buffer (Thermofisher Scientific) were mixed with 2 µg of a rescue plasmid containing a desired tag and a resistance cassette, flanked by ~500 bp homology arms, and 6 µg of a plasmid encoding hCas9 and guide RNA (pX300, addgene 42230). The mixture was then electroporated at setting 24. Approximately 24 h later, serially diluted cells were plated into 6 X 96 well dishes containing the corresponding antibiotics (final concentration 1 mg/ml hygromycin B, 1.5 mg/ml Geneticin).

Indole-3-acetic acid, auxin (57330, Merk, stock 100 mM in ethanol, final conc. 125-150 µM) was added to the culture to induce rapid degradation of AID-tagged proteins. To depolymerize microtubules, nocodazole (Sigma-aldrich, stock 1mg/ml in DMSO, final conc. 0.5 µg/ml) or colcemid (Thermofisher Scientific, stock 10 µg/ml, final conc. 100 ng/ml) was added to some cultures 30 minutes prior to 1NM-PP1 washout or just after washout, respectively. Custom made 1NM-PP1 (stock 10 mM in DMSO, final conc. 2 µM) was added to cultures to accumulate cells at the G_2_/M boundary. Palbociclib (CDK4/6 inhibitor, stock 1mM in water, final conc. 240 µM) was added to cultures to accumulate cells at the G_1_/S boundary. To prevent mitotic exit, the cell-permeable proteasome inhibitor MG132 was added to WT cultures that were harvested 60 or 90 minutes after release from G_2_ arrest (*75*)

Cell lines

All the cell lines used in this study are CDK1^as^ cells. The establishment of DT40 CDK1^as^ cells and expression of OsTIR1 were described previously (*6*). A list of cell lines, genomic modifications, rescue constructs, and the sequences of guide RNAs used in this study can be found in Supplemental Tables 6 and 7. Knock-in constructs contain tags such as AID-Clover/GFP, Clover, or Halo, resistance cassettes (hygromycin or geneticin) with loxP sequences, and 2x ~500 bp homology arms. These plasmids were assembled by Dr. Kumiko Samejima (using restriction digests) or by the Edinburgh Genome Foundry (*76*). Plasmids encoding a guide RNA and hCas9 were assembled by inserting double-stranded oligos into a plasmid pX330-U6-Chimeric_BB-CBh-hSpCas9 which was a gift from Feng Zhang (Addgene plasmid # 42230, (*77*, *78*). Plasmids pMK290 and pAID2.3C were kind gifts from Professor Kanemaki (NIG, Japan). A detailed protocol, all the plasmids (rescue constructs, guide RNAs/Cas9), and cell lines are available upon reasonable request to Dr. Kumiko Samejima. The expression of exogenous proteins and/or homozygous integration of tagged proteins by CRISPR/Cas9 technologies were confirmed by western blot analysis, by genomic DNA sequencing, or by both. Throughout this manuscript, the AID tag refers to the miniAID tag (*79*). For AID cell line names, "protein name-AID" was used for simplicity (excluding GFP/Clover). Unless otherwise specified (e.g. "no auxin") all cells expressing AID-tagged proteins were treated with auxin.

Cell Synchronization and time course experiments

CDK1^as^ synchronization and time course experiments

G_2_ synchronization and time-course experiments were performed as described previously (*6*). In brief, asynchronously growing DT40 CDK1^as^ cells were treated with 1NM-PP1 for 13 hours. As the doubling time of the CDK1^as^ cells is ~10 h, almost all (if not all) cells accumulate at G_2_/M boundary after this treatment. To deplete an AID-tagged protein just prior to mitotic entry, auxin was added to the culture for the final 3 hours of 1NM-PP1 treatment and maintained until harvest. Cells were collected just before 1NM-PP1 washout (G_2_) and typically at 2.5 minutes, 5 minutes, 7.5 minutes, 10 minutes, 15 minutes, and 30 minutes after 1NM-PP1 washout. Cells were fixed in media with 1% formaldehyde (Thermo Fisher Scientific) for 10 minutes at room temperature. Next, 2.5 M glycine was added (final concentration 125 µM) to quench excess formaldehyde for 5 minutes at room temperature and subsequently for 15 minutes on ice. Then, those cells were collected by centrifugation and rinsed with PBS once (or TBS for ChEP analysis). Cell pellets were either frozen on dry ice for Hi-C sample preparation or with liquid nitrogen for protein analysis and kept in -80˚C. Characteristics and treatments for each set of cells that served as input for Hi-C can be found in Table S1. For microscopy analysis, small amounts of cells were taken from formaldehyde-fixed cell pellets and stored in methanol/acetic acid (3:1) at -20˚C. Those cells were applied on slides, air-dried at room temperature before DNA staining with DAPI in Vectashield Plus antifade mounting medium (H-2000, Vectashield). Images were taken with a wide-field Deltavision Elite microscopy system (Olympus iX71 Inverted Fluorescence Microscope, SoftWoRx software 7.0.0 (Applied Precision Inc, Image Solutions UK Ltd), PCO edge 4.2 sCMOS camera, a X 100 UPLX-Apochromat objective/1.45 Oil). After deconvolution, 3D datasets were visualized and analyzed using Fiji. Representative images of chromatin morphologies (Fig. S1A) and the ratio of cell cycle stages present at each time point of WT CDK1^as^ cells (n=5, >100 cells counted/time point, Fig S1B) confirmed that cells reproducibly entered mitosis in a synchronous manner. Depletion of AID-tagged proteins from auxin-treated cells was confirmed by flow cytometry analysis (GFP/clover protein) and/or mass spectrometry analysis. When AID-tagged protein depletion was not homogeneous in culture, formaldehyde-fixed cells were subjected to a cell sorter (BD FACSAria) to collect GFP/Clover-negative cells with the same level of (background) fluorescence as wild-type cells that do not express GFP/Clover.

Double synchronization experiment

SMC3-AID cells were treated with Palbociclib for 14 h to synchronize them at the G_1_/S boundary. Auxin was added for the last 3 h to the cultures of “extrusion only” and “no cohesin (S+G_2_)” conditions to deplete SMC3 throughout S phase. After Palbociclib washout, cells were allowed to proceed through S phase for 7 h in the presence or absence of auxin in the medium supplemented with 25 µM PIPES. Next, 1NM-PP1 was added to the culture for the second synchronization at the G_2_/M boundary. After 3 h incubation with 1NM-PP1, auxin was added to the “no cohesin (G_2_)” culture and auxin was washed out from “extrusive cohesin only” cells. All cells were further treated with 1NM-PP1 for 9 h. Cells were collected at the end of 1NM-PP1 treatment (G_2_) and at 5 min, 15 min, and 30 minutes after 1NM-PP1 washout. Treatments for protein analysis, Hi-C and microscopy were identical to the CDK1-as synchronization protocol (previous section). Additionally, small amounts of cells were released into medium containing colcemid (final conc. 100 ng/ml) and collected at t= 30 min. Those cells were treated with 75 mM KCl for 10 minutes at room temperature (RT) prior to ice-cold methanol/acetic acid (3:1) fixation to prepare conventional chromosome spreads. The expression level of SMC3-AID-clover proteins was detected by flow cytometry analysis. Clover-positive or negative formaldehyde-fixed cells were collected using a cell sorter (BD FACSAria) to remove any dead cells and cells that still expressed detectable levels of SMC3 for no cohesin samples.

Measuring the timings of nuclear envelope breakdown

DNA fragments encoding 3x superfolder GFP or mCherry (Addgene plasmids 75385 and 75387 digested with BamHI/XhoI) and double-stranded oligos encoding BP-NLS or NES from Gg cAMP-dependent kinase inhibitor alpha were ligated into pcDNA3 (digested with BamHI/ApaI). These plasmids were randomly integrated into the genome of wild type cells, SMC2-AID cells, and SMC3-AID cells. The expression of GFP or mCherry was confirmed by flow cytometry/microscopy analysis. Wild type cells expressing Halo-laminB1 were described previously (*80*).

Live cell imaging of 3xGFP-NES Halo-Lamin B1 WT cells and its analysis

3xGFP-NES Halo-Lamin B1 WT cells were treated with 1NM-PP1 for ~13 hours in normal media, then transferred to polylysine-coated glass bottom dishes (p35G-1.5-10-C, MatTek) and incubated with SiR-DNA650 (1/1000, Spirochrome) and Halo-JF549 (1/20,000) for ~1 hour. Just prior to image acquisition, the cells were rinsed twice with live cell imaging media (Leibovitz L-15 medium supplemented with 10% FBS and 1% Chicken serum). Images were acquired every 1.5 minutes using Airyscan mode on a Zeiss LSM 880 confocal microscope with a 100x alpha Plan-Apochromat objective/1.4 Oil DIC. Seven sections were taken (1 µm interval). 3D datasets were visualized and analyzed using Fiji. Presented images show a single section of 3D data stacks. The average intensity of GFP signal within a circle drawn at nuclei, cytoplasm, and outside of cells (background) of the single section are measured. The ratio of the intensity (nuclei - background) divided by the intensity (cytoplasm – background) is shown.

Fixed cell imaging of GFP or mCherry-NES expressing cell lines and its analysis

3XGFP-NES Halo-lamin B1 WT, 3XmCherry-NES SMC2-AID, 3XmCherry-NES SMC3-AID cells were synchronized with 1NM-PP1 for 10 hours and then further treated with 1NM-PP1 in the presence or absence of auxin for 3 hours. After 1NM-PP1 washout, cells were fixed with a 4% formaldehyde solution at the corresponding time points and DNA was stained with DAPI in Vectashield Plus. Images were acquired at a 0.2 µm interval using a wide-field Deltavision Elite microscopy system and processed as above. Images show a single section of 3D data stacks. The ratio of GFP/mCherry signal intensity was measured and calculated as described above in the previous section. For each of 2 replicate experiments, twenty cells were measured for each time point.

Microscopy

(Native) chromosome spreads, DNA, Topoisomerase IIalpha and SMC2 staining, fixed cell imaging with Deltavision microscopy, and measurement of Chromosome 1 length and telomere to telomere length

Chromosome spreads were prepared with cells 30 minutes after release from the 1NM-PP1-block at G_2_. To prepare native chromosome spreads, prometaphase cells were rinsed with PBS and then fixed with methanol/acetic acid (3:1) at -20˚C. Fixed cells were dropped onto coverslips and air-dried prior to DNA staining with DAPI in Vectashield Plus. For conventional chromosome spreads, cells were treated with nocodazole (0.5 µg/ml) starting from 30 minutes prior to 1NM-PP1 washout. The cells were then collected by centrifugation, suspended in 75 mM KCl, and kept at room temperature for 10 minutes before being fixed with methanol/acetic acid (3:1) at -20˚C. Fixed cells were dropped onto coverslips and air-dried prior to DNA staining with DAPI in Vectashield Plus or subjected to indirect immunofluorescence with antibodies against Topo IIα (custom-made antibody in Guinea pig 2B2, (*81*), 1/1000 dilution) and SMC2 (custom-made antibody made with rabbit 997, (*82*), 1/500 dilution) in TEEN buffer (1mM Triethanolamine:HCl pH 8, 0.2 mM NaEDTA, 25 mM NaCl) for 1 hour. Coverslips were rinsed with KB-buffer (10 mM Tris:HCl pH 7.7, 150 mM NaCl, 0.1% BSA) for 3 x 5 minutes, incubated with anti-guinea pig and anti-rabbit secondary antibodies for 30 minutes, and then rinsed with KB-buffer for 3 x 5 minutes prior to DNA staining with DAPI in Vectashield Plus.

DNA Images were acquired at 0.2 µm intervals using a wide-field Deltavision Elite microscopy system, processed as above and analyzed using Fiji. Images with Topoisomerase IIalpha, SMC2, DNA staining were taken using Airyscan mode on a Zeiss LSM 880 confocal microscope, with a 100x alpha Plan-Apochromat objective/1.4 Oil DIC. Whole cells were imaged with a 0.13 µm interval. After airyscan processing, 3D datasets were visualized and analyzed using Fiji. Images show either a single section of 3D data stacks or max projection as stated in Figure legends.

The length of chromosome 1 was measured by drawing a line along the center of the longest chromosomes within conventional chromosome spreads using Fiji. Telomere-to-telomere length was measured by drawing a straight line between the ends of the longest chromosomes.

Imaging of Halo/GFP-tagged protein in formaldehyde-fixed cells with an airyscan microscope, and measurements of sister chromatid widths

To achieve synchronized mitotic entry, WT and AID-cells were treated with 1NM-PP1 for 10 hours followed by an additional 3 hours with auxin. More than 30 minutes prior to 1NM-PP1 washout Halo dyes were added to the medium in the following dilutions:, Halo-JF646 at 1/1000 dilution (Fig 3A), Halo-JF549 at 1/10,000 dilution (Fig 3C), or Halo-JFX549 at 1/10,000 dilution (Fig 7C, F, I Fig S7B) s. Thirty minutes after 1 NM-PP1 washout, the cells attached to polylysine-coated coverslips were rinsed with pre-warmed PBS and fixed with 4% formaldehyde, except for the SMC3-clover/SMC2-Halo cells in which cells were fixed with 4% formaldehyde in the presence of 0.5% Triton (Fig 3A). DNA was stained with DAPI in Vectashield Plus. Images were taken using Airyscan mode on a Zeiss LSM 880 or 980 confocal microscope, with a 100x alpha Plan-Apochromat objective/1.4 Oil DIC. Whole cells were imaged with a 0.13 µm interval. After airyscan processing, 3D datasets were visualized and analyzed using Fiji. Images show either a single section of 3D data stacks or max projection as stated in Figure legends.

The width of individual sister chromatids or chromosomes (paired sister chromatids) was measured by drawing a straight line perpendicular to the long axis of the sister chromatids on single section images where the intensities of SMC2-Halo were highest. The width of the single sister chromatids in the SMC3-AID cells with auxin or the chromosomes in the SMC3-AID cells without auxin was determined where the intensity of the DNA signal dropped to 50% of the maximum intensity on the line. In case of SMC3-AID cells without auxin, the distance between the SMC3-Halo and the outside edge of sister chromatid (DNA stain) was also measured to estimate the width of sister chromatids.

EdU incorporation to label single sister chromatids and measurement of gyre size

DT40 cells were treated with Palbociclib (final concentration 240 µM) for 11-14 hours. The cells were then rinsed three times with fresh media to wash out Palbociclib, and EdU was added to the medium (final concentration 0.1 µM). After ~11 hours, 1NM-PP1 was added to the medium for 5 hours. Subsequently, auxin was added to the medium for an additional 3 hours to deplete AID-tagged proteins. After washout of 1NM-PP1, cells were further treated with colcemid (0.1 µg/ml) for 30 minutes. The cells collected by centrifugation were subjected to 75 mM KCl for 10 minutes and then fixed with ice-cold methanol/acetic acid (3:1). Cells were then dropped onto coverslips and air-dried. Incorporated EdU was visualized with the Click-it EdU Imaging Kit Plus Alexa Fluor 488 (Thermo Fisher Scientific) following the manufacturer’s protocol. DNA was stained with Hoechst 33342 and sealed with Prolong Gold Antifade Mountant (Thermo Fisher Scientific). Images were taken using Airyscan mode on a Zeiss LSM 980 confocal microscope, with a 100x alpha Plan-Apochromat objective/1.4 Oil DIC (Z: 0.13 µm interval). After Airyscan processing, 3D datasets were visualized and analyzed using Fiji.

To measure the size of single gyres, EdU incorporated areas which satisfy the following three criteria were selected (see (*53*)): (1) the EdU signal does not cover the whole width of a sister chromatid, (2) some EdU signal can be seen on the other sister chromatid, (3) and the length of chromosome 1 can be measured within the same chromosome spread (See Fig 7K and (*53*)). The height of these single gyres was measured by drawing a straight line parallel to the axis of the sister chromatids. The length between the points was measured where the intensity of the EdU signal dropped to 50% of the maximum intensity on the line. The amount of genomic DNA within the single gyre was calculated considering the length of chromosome 1 (196 Mbp) in the same chromosome spread. It was noticed that even though these chromosome spreads were prepared from synchronized cell culture (30 minutes after release from G_2_ block), the measurement of single gyres tends to be more frequent with chromosome spreads which exhibit shorter than average length of chromosome 1. Therefore, a regression analysis between the length of chromosome 1 and the calculated gyre size (Mbp) was performed for each experimental dataset rather than simply calculating the average size. A functional linear regression model and the average length of chromosome 1 (WT: 10.18 µm, SMC3-AID: 7.67 µm, Fig 6D) were applied to obtain the final gyre size in WT and SMC3-AID cells.

Proteomics

Chromatin Enrichment for Proteomics (ChEP) and protein quantification based on iBAQ number provided by mass spectrometry

Cells were fixed with 1% formaldehyde for 10 minutes. To inactivate the formaldehyde, 1/20 volume of 2.5 M glycine was added and incubated for 5 minutes before harvesting cells. The fixed cells were washed with TBS (50 mM Tris pH 7.5, 150 mM NaCl), and snap-frozen in liquid nitrogen for storage at −80°C. Once thawed on ice, cells were processed according to the ChEP protocol (*36*, *80*, *83*). In brief, formaldehyde-crosslinked cells were lysed in lysis buffer (25 mM Tris pH 7.5, 0.1% Triton X-100, 85 mM KCl). Chromatin was extracted with SDS buffer (50 mM Tris pH 7.5, 10 mM EDTA, 4% SDS), and was washed twice under denaturing conditions (6 M Urea and 1% SDS), followed by a wash with SDS buffer. The DNA content of the chromatin fractions was measured using a Qubit with HS DNA QuantIT (Thermo Fisher Scientific) according to the manufacturer’s instructions.

ChEP chromatin was processed for mass spectrometry by in-gel trypsin digestion. A detailed procedure is described in (*35*). Following digestion, samples were acidified (pH<3) and spun onto StageTips as described (*84*). Peptides were then eluted in 40 μL of 80% acetonitrile in 0.1% TFA and concentrated down to 1 μL by vacuum centrifugation (Concentrator 5301, Eppendorf, UK). The peptide samples were then prepared for LC-MS/MS analysis by diluting each one to 5 μL by 0.1% TFA.

LC-MS analyses were performed on a Q Exactive and on an Orbitrap Exploris™ 480 Mass Spectrometer (both from Thermo Fisher Scientific, UK) both coupled on-line, to Ultimate 3000 HPLCs (Dionex, Thermo Fisher Scientific, UK). Peptides were separated on a 50 cm (2 µm particle size) EASY-Spray column (Thermo Scientific, UK), which was assembled on an EASY-Spray source (Thermo Scientific, UK) and operated constantly at 50oC. Mobile phase A consisted of 0.1% formic acid in LC-MS grade water and mobile phase B consisted of 80% acetonitrile and 0.1% formic acid. Peptides were loaded onto the column at a flow rate of 0.3 μL min-1 and eluted at a flow rate of 0.25 μL min-1 according to the following gradient: 2 to 40% mobile phase B in 180 min and then to 95% in 11 min. Mobile phase B was retained at 95% for 5 minutes and returned back to 2% a minute after until the end of the run (220 min).

For the samples on the Q Exactive, Data Dependent Acquisition (DDA) was used, and the instrument parameters were the same as described (*80*). On Orbitrap Exploris™ 480, for DDA, survey scans were performed at 120,000 resolution with scan range of 350-1500 m/z, normalized AGC target of 3.0E106 and injection time of 50ms. MS2 scans were performed with an isolation window of 1.4 Thomson, and with orbitrap resolution of 15,000. We used HCD fragmentation with normalized collision energy that was set to 30% (*85*), normalized AGC target at 8.0E104, and maximum injection time at 60ms. The cycle time was set at 3 seconds and only ions with charges between 2 and 7 were chosen for MS2.

For Data Independent Acquisition (DIA) Survey scans were recorded at 120,000 resolution (scan range 350-1650 m/z) with an ion target of 5.0e6, and injection time of 20 ms. MS2 was performed in the orbitrap at 30,000 resolution with a scan range of 200-2000 m/z, maximum injection time of 55 ms and AGC target of 3.0E6 ions. We used HCD fragmentation with stepped collision energy of 25.5, 27 and 30. We used variable isolation windows throughout the scan range ranging from 10.5 to 50.5 m/z. Shorter isolation windows (10.5-18.5 m/z) were applied from 400-800 m/z and then gradually increased to 50.5 m/z until the end of the scan range. The default charge state was set to 3. Data for both survey and MS/MS scans were acquired in profile mode.

All mass spectrometry raw files are available upon request and will be deposited to the ProteomeXchange Consortium (http://proteomecentral.proteomexchange.org) via the PRIDE partner repository. The raw files were processed by MaxQuant version 2.4.9.0 (*86*) and peptide searches were conducted against the chicken reference proteome set (UP000000539) of UniProt database (Release 2024_01) with additional sequences from our in-house database of chicken proteins, using the Andromeda search engine (*87*).

Quantification of key proteins associated with chromatin is based on ChEP SILAC analysis data of WT cells with no auxin (n=3), WT cells with auxin (n=3), SMC2-AID cells plus auxin (n=3), and SMC3-AID plus auxin (n=2-4) cells. Note that the n differs because CAP-H, CAP-H2, CAP-G2 proteins were undetected in some of the SMC3-AID samples. Data were normalized against the values of Histone H4. Log 2 ratio against G_2_ samples of the same cells with no auxin (shown as “con” in Fig S2A) (Fig 1C. Fig S2).

The number of condensin I, II, and cohesin complexes on chromatin (per Mb DNA) shown in Fig 1D was calculated as follows. The average iBAQ number of corresponding subunits (CAP-G, CAP-H, CAP-D2 for condensin I; CAP-G2, CAP-H2, CAP-D3 for condensin II; and SMC1, SMC3, RAD21 for cohesin) and Histone H4 were obtained from WT cells (n=3 no auxin, n=3 with auxin). The number of condensin I, II, and cohesin complexes on chromatin (per Mb DNA) was deduced by assuming 2 Histone H4 proteins are equivalent to 180 bp of DNA.

Chromatin Enrichment for Proteomics (ChEP) and protein quantification with halo-tagged spike-in control protein by mass spectrometry

Absolute quantification of SMC3, CAP-H, and CAP-H2 on chromatin was performed with a heavy amino acid labeled Halo tagged Histone H4 proteins as a spike-in control. Halo-H4 fusion protein was expressed in E. coli and purified as described below. The ChEP chromatin with the spike-in was processed by filter-aided sample preparation (FASP) (*88*) to generate tryptic peptides which were analyzed by a mass spectrometer by data-independent acquisition (DIA). The output spectra were analyzed by DIA-NN (version 1.8.1) (*89*, *90*)**.**

The number of corresponding proteins (SMC3, CAP-H, CAP-H2) on chromatin are calculated as follows. First, total MS1 intensities of peptides originating from each protein (SMC3, CAP-H, CAP-H2, Halo tag, histone H4) are obtained. As MS1 intensities are influenced by the composition of peptides, the MS1 (precursor ions) intensity values are adjusted using the Halo-tag protein as a standard protein. The Histone H4 to Halo tag adjustment value was obtained using recombinant Halo-Histone H4. The SMC3, CAP-H, CAP-H2 to Halo tag adjustment values were obtained using the ChEP data of SMC3-Halo, CAP-H-Halo, Halo-CAP-H2 cells, respectively. Finally, the number of SMC3, CAP-H, CAP-H2 on chromatin (per Mb DNA) was deduced by assuming 2 Histone H4 proteins are equivalent to 180 bp of DNA.

Expression and purification His_6_-Halo-HistoneH4 of protein and absolute quantification of Halo-tagged proteins

A plasmid encoding the His_6_-Halo-GgHistoneH4 protein was constructed by inserting a synthesized Histone H4 gene into a pH6HTN His6Halotag T7 vector (Promega), which has a TEV cleavage site after the Halo tag. The plasmid was then transformed into an E. coli mutant strain BL21 (DE3) ΔlysA ΔargA, generously provided by Professor Ronald Hay of the University of Dundee, UK (*91*). The E. coli cells were cultured in M9 medium supplemented with 0.2% glucose, 0.1% thiamine, 100 μg/mL ampicillin, 1 mM MgSO4, 50 μg/mL 13C6, 15N4-arginine, 50 μg/mL 13C6, 15N2-lysine, and a non-essential amino acid mix. Protein expression was induced by adding IPTG to a final concentration of 0.1 mM and incubating at 37°C for 3-4 hours. The E. coli pellets were then snap-frozen in liquid nitrogen and stored at -80°C. The His_6_-Halo-HistoneH4 protein (shown in Fig S1C) was subsequently purified from the inclusion bodies using a Dounce glass/glass homogenizer as described before (*92*). Briefly, inclusion bodies were solubilized by soaking in 500 ml of DMSO and resuspension in unfolding buffer containing 7 M Guanidine, 20 mM Tris pH 7.5, and 10 mM DTT. After solubilization, a three-step dialysis against Urea buffer (7 M Urea, 100 mM NaCl, 10 mM Tris pH 8.0, 1 mM EDTA, and 5 mM β-mercaptoethanol) was performed. The sample was then loaded onto a 5 ml HiTrap SP cation exchange column (Cytiva) and eluted using a linear gradient from 100 mM to 1 M NaCl in 7 M Urea, 10 mM Tris pH 8.0, 1 mM EDTA, and 1 mM DTT. The fractions containing the pure histone were pooled, aliquoted, and snap-frozen before storage at -80°C. For mass spectrometry analysis to achieve absolute quantification, 0.16-0.4 µg of purified His6-Halo-HistoneH4 protein was mixed with 30 µg of ChEP sample of SMC3-Halo, CAP-H-Halo, or Halo-CAP-H2 cells.

Electron microscopy (EM)

EM sample preparation, SBF-SEM imaging, and data acquisition

DT40 cells were attached to a MatTek dish coated with poly-L-lysine and fixed with 2.5% EM grade Glutaraldehyde in 0.1 M sodium cacodylate buffer for 1 hour (pH 7.4). They were then blocked for 15 minutes using a blocking buffer (10 mM glycine, 10 mM potassium cyanide in 0.1 M sodium cacodylate buffer). Cellular DNA was stained with DRAQ5 (10 µM) in 0.1 M cacodylate buffer for 30 minutes, followed by bathing the cells in 2.5 mM diaminobenzidine tetrahydrochloride (Sigma) in 0.1 M sodium cacodylate buffer.

To identify the cells and proceed with photooxidation, the fixed cells were placed on a wide-field DeltaVision Elite (Applied Precision) microscope with a PCO edge 4.2 sCMOS camera and a 100× NA 1.45 UPLX Apochromat objective with oil immersion (refractive index = 1.514) and a Cy5 filter set (640 ± 30) using SoftWoRx 3.6 (Applied Precision) software. The cells were photooxidized by continuous epi-fluorescence illumination for 2 minutes.

Next, the cells were rinsed 5 times for 2 minutes each with 0.1 M sodium cacodylate buffer and then stained for 1 hour with 2% osmium tetroxide and 1.5% potassium ferrocyanide. Subsequently, 1% tannic acid was added to the cells as a mordant for 20 minutes at room temperature, followed by the second osmification step adding 2% osmium tetroxide for 1 hour. Finally, Walton’s lead aspartate (0.02 M lead nitrate in 0.03 M L-aspartic acid, adjusted to pH 5.5) was added. The cells were rinsed 5 times for 2 minutes each with ddH2O between each step. The cells were dehydrated in a graded ethanol series of 30%, 50%, 70%, and 90% in ddH2O for 5 minutes each, followed by 2 rounds of 5 minutes in 100% ethanol, infiltrated with Agar 100 Hard Premix resin (Agar Scientific) at a 1:1 ratio (resin:100% ethanol), and then resin-only for 30 minutes. The cells were embedded in 2 mm of 100% fresh resin and cured for 48 hours at 60°C.

To prepare the blocks, the resin was removed from the MatTek dish using pliers. The photo-oxidized cells and the coordinates on the coverslip were identified using a stereoscopic microscope, and the block was cut with a junior hacksaw and glued to a cryopin (cell side up) using silver epoxy resin. Then, the block was trimmed using a microtome to select the region of interest. The samples were covered with 10 nm gold/platinum before SBF-SEM imaging.

SBF-SEM images were acquired using the Gatan 3View serial block-face system (Gatan, Pleasanton, CA) installed on a FEI Quanta 250 FEG scanning electron microscope (FEI Company, Hillsboro, OR). Images were collected with the following parameters: a pixel size of 0.004 X 0.004 X 0.06 nm, a dwell time of 4 seconds, magnification 5,400, with a chamber pressure of 70 Pa at 3.2 kV.

3D Reconstruction, modeling and segmentation of SBF-SEM images

The electron microscopy images were reconstructed and annotated using AMIRA software (FEI). The selection of the chromosomes was done by setting up a threshold to differentiate between the cytoplasm and the chromosomes. Then, interactive thresholding masks and the Magic Wand tool were used in the segmentation environment to select the chromosomes. The chromosomes were separated using the "separate objects" tool with a 3D interpretation and a neighborhood criterion of 26 connected elements. Chromosomes were identified based on size, length, and centromere position; only chromosome 1 to 5 and chromosome Z could be identified. The Label Analysis module was used to measure geometrical values such as the volume of all identified chromosomes. Surface renders were generated using unconstrained smoothing at level 5. Chromosome length and width were calculated using two methods: 1) using the skeleton tool to draw a line along the chromosomes and measure the length, and 2) using the measuring tool to draw 10 lines along and across the chromosomes.

Hi-C protocol

Chromosome conformation capture (Hi-C 2.0) was performed as described previously (*93*). Briefly, ~5x106 crosslinked, frozen cells were thawed on ice and lysed for 15 minutes in ice-cold lysis buffer (10 mM Tris-HCl pH8.0, 10 mM NaCl, 0.2% Igepal CA-630) in the presence of Halt protease inhibitors (Thermo Fisher, 78429). Cells were disrupted with pestle A for 2x 30 strokes, centrifuged at 2500 xg, washed, and resuspended in NEBuffer 3.1 (NEB, B9200S). Chromatin was solubilized in 0.1% SDS at 65°C for 10 minutes, quenched by 1% Triton X-100 (Sigma, 93443), and digested with 400 units of DpnII (NEB, R0543M) overnight at 37°C in a total volume of 475 µl. DpnII was inactivated at 65°C for 20 minutes. Digested overhangs were filled in in the presence of 250 nM biotin-14-dATP (Invitrogen, 19524-016) using the large Klenow fragment of DNA polymerase I, (NEB, M0210) for 4 hours at 23°C. Blunted ends were ligated in situ with 50 units of T4 DNA ligase (Invitrogen, 100004817) for 4 hours at 16°C (1200 µl total volume). Cross-links of ligated chromatin were reversed overnight at 65°C using 2 batches of 50 μl of 10 mg/ml proteinase K (Invitrogen, 25530-031). DNA was isolated with 1:1 phenol:chloroform, followed by desalting with an Amicon Ultra-0.5 Centrifugal Filter (EMD Millipore, UFC500396) and 30 minutes of RNase A incubation (Thermo Scientific, EN0531). Biotin was removed from unligated ends by incubation with 15 units of T4 DNA polymerase in the presence of 25 nM dATP/dGTP in NEBuffer 2.1 at 20°C for 4 hours. DNA was sheared using an E220 evolution sonicator (Covaris) and size selected with Agencourt AMPure® XP (Beckman Coulter) to 150- 350 bps. DNA ends were repaired in a mixture of T4 polynucleotide kinase (25 units; NEB, M0201), T4 DNA polymerase (7.5 units; NEB, M0203L) and the large Klenow fragment of DNA polymerase I (2.5 units; NEB, M0210) at 20°C for 30 minutes. The enzymes were inactivated for 20 minutes at 75°C before A-tailing with dATP using 15 units of exonuclease deficient Klenow (Klenow 3’ → 5’ exo-) (NEB, M0212L) at 37°C for 30 minutes. Biotinylated DNA was bound to 12 µl of streptavidin-coated myOne C1 beads (Life Technologies, 650.01). Illumina paired-end adapters were ligated to bead-bound DNA using T4 DNA ligase for 2 hours at room temperature. To determine the minimal number of PCR cycles for Hi-C library generation, a PCR titration was performed before the production PCR (primers PE1.0 and PE2.0). Primers were separated from the 350-550 base pair library using Ampure size selection prior to 50 bp paired-end sequencing on an Illumina HiSeq4000 sequencer.

Hi-C data analysis

Mapping sequenced Hi-C reads

Hi-C paired-end fastq files (BioProject number PRJNA1091327, GEO GSE262525) were mapped to bGalGal1.mat.broiler.GRCg7b (NCBI RefSeq assembly GCF_016699485.2) using the distiller-nf pipeline ([https://github.com/open2c/distiller-nf/](https://github.com/open2c/distiller-nf/blob/master/distiller.nf)), as described previously (*94*, *95*). Briefly, reads were aligned with bwa mem (*96*), and resulting alignments were parsed into Hi-C pairs with the pairtools package (*97*). We then removed PCR duplicates and kept pairs of uniquely mapped alignments. Pairs with phred scores ≥ 30 for both sides were aggregated into binned contact matrices in the cooler format (*98*). Contact matrices were normalized using iterative correction balancing after excluding the first two diagonals (*99*). Bins with extreme coverage were excluded using the MADmax filter on genomic coverage (*47*). Multi-cooler files and their mapping statistics were submitted to GEO (GSE262525).

Contact frequency (*P*(*s*)) curves from pairs of interactions.

To calculate the functions of contact frequency *P*(*s*) vs genomic separation s from Hi-C pairs at bp-level resolution, we used pairtools. Briefly, we split all genomic distances *s* between 1 kb and 1 Gb into bins of exponentially increasing widths, with 128 bins per 10-fold change in s. For every such separation bin, we identified the number of observed cis-interactions within this range of separations (numerator) and the number of all loci pairs separated by such distances in cis (denominator). We then used the built-in functionality of pairtools to smooth the numerator and denominator separately with a Gaussian kernel in the log10-s space with the width of log10_sigma = 0.03. The contact frequency *P*(*s*) was calculated as the ratio of smoothed numerator and denominator. Finally, for these analyses, we only used pairs of loci located on chromosomal arms that were longer than 100 Mb.

Contact frequency (*P*(*s*)) curves for binned data

Separately, we also computed the *P*(*s*) curves from binned and balanced Hi-C maps at all used binning resolutions using cooltools (v0.5.035) (*100*). The *P*(*s*) curves produced by cooltools were used to normalize Hi-C maps for distance-dependent decay of contact frequency (i.e. to calculate observed-over-expected signal, OE) and enable compartment detection, insulation, and loop analysis.

Generating a high-resolution “merged G_2_” dataset

The limited sequencing depth of individual libraries did not allow de-novo loop calling independently for each sample. Therefore, we merged 9 Hi-C libraries from all samples at the G_2_ time point that still contained cohesin (i.e., we excluded all SMC3-AID libraries that were induced with auxin addition):

1)	SMC3-AID-G2-S1-R1-spna

2)	SMC3-CAPH-AID-G2-S2-R1-spna

3)	SMC3-CAPH2-AID-30min-S2-R1-spna

4)	SMC3-AID-G2-S4-R1-AN-SPnAa

5)	SMC3-AID-G2-S4-R1-NN-SPnaa

6)	CAPH-AID-G2-S3-R5-spnA

7)	CAPH2-AID-G2-S3-R5-spnA

8)	SMC2-AID-G2-S4-R5-spNA

9)	WT-G2-S4-R5-spna

We generated “merged G_2_” coolers at 1kb, 5kb, 10kb, and 100 kb resolutions and balanced them using the *merge_coolers* and *balance_cooler* functions of the *cooler* library.

Loop analysis

Loops (dots) were called from the merged G_2_ cooler at both 5 kbp and 10 kbp resolution using the modified HICCUPS approach via *cooltools.dots* (<https://cooltools.readthedocs.io/en/latest/notebooks/dots.html>) with the following settings: max_loci_separation=1Mbp, clustering_radius=30kbp, tile_size=5Mbp, lambda_bin_fdr=0.15, max_nans_tolerated=2. Loop calls from both resolutions were combined into a single file containing all loop locations.

We used KDTrees (*scipy.spatial.cKDTree.query_pairs*) to identify and remove dots within a radius of 30kbp (same as clustering radius using the dot calling). For each subset of duplicates, one instance was arbitrarily selected to serve as the representative of the subset.

To quantify the strength of these dots in other conditions and time points, we then applied *cooltools.pileup* to extract chunks of Hi-C maps around each dot (referred to as Hi-C snippets), at a resolution of 1 kb (flank_bp=50 kb). We then averaged these snippets to create an average Hi-C contact map for each dot, also known as a “pileup”. The dot enrichment was calculated as the mean distance-normalized signal (observed/expected) within a 3x3 pixel window around the pileup’s center.

Insulation analysis (TADs)

The following protocol was used to create TAD pileups for each Hi-C library:

1. Insulating boundaries were called at 10 kb resolution on the "merged G_2_" cooler using cooltools.api.insulation.calculate_insulation_score (window= 200 kb). Only galGal7b chromosome arms larger than 10 Mb were considered. Boundaries with strength < 0.1 were filtered out to avoid spurious calls.
2. TADs were defined as regions between adjacent strong boundaries. A list of TADs was generated, and only TADs between 100-700 kb in size were selected for further analysis.

The TADs detected in the "merged G_2_" cooler in steps 1-2 were then visualized for each individual library in steps 3-5:

3. For each Hi-C library, cooltools.pileup was used to create a stack of regions (referred to as snippets) extending +/- 1 Mb around the center of each TAD. Choosing a 2 Mb window guaranteed that flanks would be sufficiently large for TADs of all selected sizes.
4. The stack of snippets was then iterated through, and for each snip, a region of +/- the size of the associated TAD was extracted. Each extracted region was rescaled to a fixed size (100x100 pixels) using mirnylib.numutils.zoomArray.
5. The scaled snippets were averaged to produce the "average TAD pileup", representing the contact footprint of an "average" TAD for each condition and time point.

Compartment analysis

The strength of compartmentalization in the G_2_ libraries was computed in two steps.

First, we extracted the first eigenvector (EV1) of all cis Hi-C contact maps from the “merged G_2_” cooler. The largest eigenvector of the Hi-C matrix was computed using *cooltools.eigs_cis* at 100 kb and oriented to have a positive correlation with the GC content of the galGal7b reference. When computing the eigenvector, only galGal7b chromosome arms larger than 10 Mb were retained. Only “cis” eigenvectors were calculated. Separately, the first 3 eigenvectors were derived from each cooler, but not used in this section’s analyses. From this analysis, EV1 was submitted to GEO (GSE262525).

Second, we generated saddleplots for each Hi-C library, providing 100 kb coolers for individual libraries, their associated *P*(*s*), and the “merged G_2_” eigenvector as input (see (*101*)). This saddleplot showed interactions between 10 groups of genomic bins, sorted and grouped according to their eigenvector values; bins with eigenvector values below the 2.5th percentile and above the 97.5th percentile were ignored. In these plots, AA interactions are shown in the bottom right corner, BB interactions are in the top left corner, and AB interactions are in the top right corner. Compartmentalization scores were then calculated as the ratio of these corner pixels: (AA+BB)/(AB+BA). All scores were normalized to the G_2_ value for the condition set.

Simulations of cohesin displacement by condensins

To model the transition between loosely packed interphase chromatin (formed by cohesin) and the condensed mitotic state (formed by condensin II) during prophase, we combined the polymer models of interphase (*1*) and mitotic loop extrusion (*30*). Simulations were performed using the open2c/polychrom library (*102*), similar to previous studies (*103*). This library utilizes OpenMM (*104*) for GPU-accelerated molecular dynamics.

We represented chromatin as a chain of 1 kb monomers (10 nm diameter), with each simulation modeling 20 Mb and incorporating three system copies. Lengths are expressed in monomer units, and energies in kT. The model incorporated the following forces:

-	Bead Connectivity: Harmonic bonds (*polychrom.forces.harmonic_bonds* with bondLength=1.0, bondWiggleDistance=0.1) linked monomers into a polymer chain.

-	Polymer Stiffness: Angle forces (*polychrom.forces.angle_force* with k=1.5) provided polymer rigidity.

-	Excluded Volume: Repulsive forces (*polychrom.forces.polynomial_repulsive* with trunc=1.5, radiusMult=1.05) prevented spatial overlap, simulating real polymer behavior.

Periodic boundary conditions maintained a density of 0.224 monomers per cubic nanometer, and a variable Langevin thermostat (collision rate: 0.01 [1/picosecond], error_tol=0.01) controlled temperature.

Our kinetic polymer simulation consists of two stages. First, we modeled interphase chromatin with active cohesin loop extrusion, optimizing parameters to generate a steady-state ensemble of polymer conformations representing a population of cells. Second, we selected 400 conformations from this interphase ensemble. Each conformation served as the starting point for simulations modeling the initial interphase-to-mitotic transition within a single nucleus. Averaging these 400 simulations approximates the population-level behavior observable in our time-resolved Hi-C data.

Polymer model of late interphase/prophase cohesin extrusion

Our prophase simulations with loop extrusion follow the framework established in (*48*). Simulations are initialized with random walk conformations, and we assess equilibration by determining the convergence of contact probability scaling curves. This convergence occurred around 3.5e8 molecular dynamics steps.

The dynamics of cohesin loop extrusion in late interphase and prophase were modeled via a parallel 1-dimensional non-equilibrium Monte-Carlo simulation with the following rules:

-  	**1D extrusion dynamics**: the polymer chain is represented as a 1-D lattice. Here, a loop-extruding factor (LEF) was conceived and implemented as a walker with a pair of legs that move probabilistically in opposite directions. The location of the legs represents either end of the loop that the LEF extrudes. LEFs load at randomly chosen lattice sites. They extrude loops for a period of time before unbinding, a process that continues throughout the simulation.

-	**Collisions and pausing**: When the legs of two LEFs collide, they stall until one LEF unbinds.

-	**CTCF barriers**: CTCF sites act as bidirectional barriers. Upon encountering a CTCF barrier, LEFs have a 90% capture probability. Captured LEFs remained at CTCF barriers until unbinding from chromatin. The CTCF barriers were located at the following positions on each of the 3 lattices:

[0, 55, 201, 239, 516, 551, 666, 717, 806, 1366, 1714, 1905, 2050, 2214, 2386, 2456, 2586, 2713, 2829, 3088, 3315, 3875, 4196, 4480, 4593, 4845, 4972, 5008, 5131, 5228, 5437, 5593, 5626, 5686, 5786, 5840, 5895, 6198, 6250, 6427, 6602, 6818, 6950, 7632, 7898, 8175, 8328, 8423, 8646, 9028, 9395, 9505, 9571, 9770, 9920, 10182, 10231, 10422, 10684, 10853, 10912, 10991, 11287, 11412, 11570, 11631, 12107, 12203, 12388, 12459, 12729, 13180, 13499, 13554, 13634, 13769, 13930, 13959, 14143, 14241, 14315, 14463, 14497, 14619, 14650, 14800, 15100, 15177, 15321, 15460, 15492, 15520, 15620, 15692, 15747, 15837, 16245, 16415, 16569, 16737, 16844, 17029, 17181, 17189, 17254, 17684, 17725, 17837, 17885, 17961, 18072, 18146, 18261, 18544, 18721, 18950, 19184, 19258, 19414, 19528, 19570, 19610, 19753, 19809, 19999]

LEFs from the 1D simulation were then implemented as additional harmonic bonds within the 3D polymer model. These bonds were updated in the 3D simulation to track with the LEFs positions in the 1D simulation.

The loop extrusion process can be characterized by processivity (λ, the amount of chromatin extruded by LEF over its lifetime in the absence of obstacles), separation (d, average spacing between extruders), and extrusion speed (*48*). From this previous work, we adopted established optimal values for processivity and separation (λ = 100 kb, d = 300 kb) . To determine the extrusion speed, we relied on the previous modeling (*1*, *48*)) and iFRAP data (*105*). These studies estimated the average cohesin loop size at ~100kb and the residence time at ~10 minutes. Together, this implied an effective cohesin extrusion velocity of 10 kb/min, which we used in our interphase model.

Finally, we found the conversion ratio between the time units in simulations and the real world. Live-cell imaging data (*106*) showed that a 515 kb chromatin region has a Rouse time of ~120 minutes. Our own polymer simulations without cohesin yielded a Rouse time of 6e6 MD steps. Matching the time scales of observed and simulated polymer diffusion yields the time unit conversion ratio. Together, this allowed us to map the real-world cohesins that extruded 100 kb loops at 10 kb/min to model LEFs with λ= 100 kb and LEF step intervals of 10^4 MD steps.

Polymer model of prophase condensin extrusion

We selected 400 conformations from the interphase ensemble. Each served as the starting point for a simulation modeling the initial interphase-to-mitotic transition within a single nucleus. On top of active cohesin LEFs, we introduced extra condensin loop extruders to these conformations, using a method similar to cohesin extruders (1D Monte-Carlo coupled to the 3D polymer). Key differences between condensin and cohesin LEFs:

-	**Loading and stable binding**: Condensins loaded synchronously and remain bound throughout the simulation.

-	**CTCF Interaction:** Condensins bypassed CTCF barriers. This is a reasonable assumption given experimental evidence that CTCF-bound elements do not block loop extrusion by condensins (*107*).

Cohesin extrusion persisted during this phase. We explored various potential interaction mechanisms upon collision between condensin and cohesin LEF legs (as described in the main text):

-	**Stalling:** Identical to interactions between two condensins or two cohesins (remain stalled until unbinding).

-	**Bypass:** condensin LEF continued past the cohesin LEF.

-	**Push:** condensin LEF displaced the cohesin LEF.

-	**Unload:** condensin LEF removed the cohesin LEF.

Importantly, these interactions occur regardless of whether the cohesin was trapped at a CTCF barrier. For condensin LEFs, we used the following parameters: infinite lifetime, average separation d= 700 kb, speed of extrusion = 120 kb/min (see below, chapter “Estimation of loop extrusion speed”, and Discussion).

Simulations of cohesion

Our tethered sister chromatid simulations built upon the previously established methodology (*30*). Crucially, unlike the prophase scenario, which aimed to recapitulate the dynamic process of cohesin loop removal, our simulations of cohesion were designed to sample a dynamic steady state. As above, simulations were conducted using *open2c/polychrom* with the same parameters, unless specified. We modeled two chains of 1-kb monomers (10 nm diameter), each representing an 18 Mb chromatin fiber. Polymer bonds and lengths remained identical to the prophase simulations. To induce sister chromatid separation, we omitted periodic boundary conditions and confinement, simulating good solvent conditions.

Loop extrusion mechanics were largely consistent with the prophase simulations described above. Key distinctions lie in the two additional SMC factors acting on the polymers:

-  	Condensin I: we modeled a population of LEFs that represent condensins I. In contrast to Condensin-II, Condensin I LEFs are abundant and highly dynamic. The extrusion parameters were adjusted (λ= 800, d= 50) to reflect this and produce the dense extrusion regime required for mitotic chromosomes (*25*). As with condensins II, condensin-I leg collisions induced stalling.

-  	Cohesive cohesin: we implemented cohesive cohesins as two-legged walkers (LEFs) positioned on separate chromosomes. The legs of cohesin LEFs stepped forward or backward in steps with equal rates, resulting in random diffusion along the chromosome. Within the 3D simulation, these manifested as bonds between the polymer chains. Cohesins bound randomly (average 300 kb spacing) at the simulation start and remained bound throughout the simulation. Collisions between cohesins’ legs also induced stalling.

Bypassing and stalling interactions between condensins and cohesive cohesins were implemented in the same way as those in the prophase simulations described above. To ensure steady-state LEF distributions, simulations ran for 7.5e6 MD steps. Both cohesin and condensin steps occurred every 750 MD steps, and conformations were saved every 7500 MD steps.

Generation of synthetic fluorescence profiles.

Out of the 7.5e6 MD steps in these simulations, 500 conformations from the second half of the simulation were saved and used to extract a radial density profile of the monomers. Three different categories of monomers were considered:

1. Condensin monomers: monomers associated with a condensin LEF at that step of the simulation.
2. Cohesin monomers: monomers associated with cohesive cohesin at that step of the simulation.
3. Generic monomers: all monomers, regardless of whether they are associated with cohesin or condensin.

The radial profile was constructed in the following manner:

1. For each conformation, the spine of each individual sister chromatid is extracted:
   a. The polymer conformation was represented as a graph (using the *networkx* Python package) where monomers were nodes and edges represented the bonds that connected monomers.
   b. In addition to the polymer bonds of each chromatid, bonds were added between monomers at which a condensin LEF resides.
   c. The shortested path between the first and last monomer of each chromatid is calculated (using *networkx.shortest_path*).

2. Monomers from the spines of each sister chromatid were paired to define sections.
   a. The distance of each spine monomer to every monomer on the **other** spine was calculated.
   b. Every spine monomer of the first chromatid was paired with the monomer on the other spine that was closest to it in 3D space.
   c. The above step was repeated for every spine monomer of the second chromatid.
   d. The two lists of pairing were compared, and only pairs present in both lists (i.e. spine monomer pairs that were mutually nearest-neighbor) were considered.
   e. To mimic perpendicular sections of the sister chromatid from experimental fluorescence, a line was drawn through these pairs.

1. For each monomer category (generic, condensin & cohesin), monomer positions were assigned to sections and collapsed onto the lines.
   a. For each line section, the midpoint of the line was assigned as the origin, and all positions were recalculated around that origin.
   b. The projection and rejection of each position vector onto the section line were calculated. The projection referred to the footprint (dot product) between the position vector and the section line vector, and the rejection represented the perpendicular distance of that monomer from the line.
   c. Monomers were assigned to the nearest line section (i.e. the line section that minimized the rejection), and the associated section projection was chosen for that monomer.
   d. Once this was done for every monomer (within a category), a histogram of projection values was computed to represent the density profile obtained from a single conformation.
   e. This process was repeated for all 500 conformations, and the counts associated with each histogram were aggregated and visualized (as shown in the Fig. 3B, right panels)

Polymer models of mitotic chromosomes

The design of molecular dynamics simulations of chromosomes

In order to provide structural interpretation to our Hi-C data of prometaphase chromosomes in key studied mutants, we built polymer models of chromosomes using molecular dynamics simulations. Overall, our approach was roughly based on those from (*5*, *6*), but included several major novel modifications.

The models and molecular dynamics simulations were implemented in HOOMD-blue v. 4.0.0 (*108*), a simulation package that provided a rich collection of simulation methods and a flexible and high-performance Python interface.

In all simulations, we modeled 10 nm ‘beads-on-string’ chromatin fiber of nucleosomes as a chain of particles, each representing one nucleosome with 200 bp of DNA. In HOOMD, simulations use a normalized system of units, where particles have a mass of 1.0 [mass] and a diameter of 1.0 [length], where one unit of length corresponds to 10 nm in real coordinates. The particles were connected by harmonic springs (equilibrium length L_bond_ = 1 [length], stiffness = 10 k_B_T/[length]^2. To prevent spatial overlap and model chromatin's excluded volume, particles interacted via the standard DPD repulsion potential (*109*):

We used a repulsion coefficient A_rep_ = 3.0 k_B_T. This prevented overlap while still allowing some strand passing for efficient topological equilibration.

To model the thermal equilibration of chromatin conformation, we subjected the system to the Dissipative Particle Dynamics thermostat (*109*) (target temperature T=1 [temperature], friction coefficient γ=1.0 [mass / time]). Each simulation contained 500,000 particles (representing 100 Mb of chromatin) and ran for at least 1.5e7 steps of dt= 0.05 [time] to ensure thermal equilibration. Three replicates were conducted for each tested set of simulation parameters. Each simulation replicate was executed for 2 days on one NVIDIA GPU (P100, V100, RTX 6000, or A100), provided by the CLIP cluster of Vienna Biocenter. The resulting sets of particle coordinates were analyzed with Python and visualized with Matplotlib (*110*) and Blender 4.0 Python API (https://www.blender.org).

Simulations of SMC3-AID/CAPH-AID chromosomes

To reconstruct the structure of chromosomes formed by condensin II only (SMC3/CAPH-AID cells) we used the model of chromatin fiber described above (which we refer to as model principle (i)) and imposed three additional model principles:

(ii) Formation of loop arrays by Condensin II:

-	The chromatin fiber was compacted into an array of consecutive loops separated by small gaps (*6*, *12*). The size of each loop was drawn randomly and independently from an exponential distribution with an average of l_loop_.

-	The pair of particles corresponding to the two “anchors” of one loop were connected by a harmonic bond (equilibrium length L_loopbond_ = 1.0 [length], stiffness coefficient = 10 k_B_T/[length]^2).

-	Each pair of consecutive loops had their anchors separated by a short “gap” chain segment containing n_gap_ particles. A harmonic force stretched this gap (equilibrium length, L_gap_ = n_gap_+1 [length], stiffness coefficient = 10 k_B_T/[length]^2).

-	For each computational replicate, the loop array was generated randomly and independently.

(iii) Condensin-Independent Condensation (*66*, *111*, *112*):

-	The resulting bottle-brush of chromatin loops was constrained by a cylindrical wall force. We used the built-in HOOMD-blue Gaussian wall potential (epsilon = 1.0 k_B_T,  sigma = 0.5 [length]).

-	Cylinder radius R_cyl_ and length L_cyl_ were calculated based on the target linear and volume density of chromatin (see below).

-	An extra technical challenge arose because HOOMD applied the bounding potential within the specified cylinder, leading to excess volume compaction. To compensate, we slightly increased the input radius and length of the cylinder, ensuring the confining potential reached 0.3 k_B_T at the target radius and length.

(iv) Formation of a helical chromosome:

To model formation of helical mitotic chromosomes, we used a combination of potentials that weakly “nudged” the chain of loops into a helical path:

-	We arranged the chromosome linearly along the axis of the confining cylinder by pulling its two ends to the opposite caps of the cylinder with a force of 1.0 [mass * length / time2].

-	In the initial conformation, we arranged the particles of loop anchors in a regular helix with a specified number of turns.

-	To maintain the number of helical turns of the chromosome throughout simulations, we added a “spool” potential. This potential, a built-in HOOMD Gaussian wall (epsilon=6 k_B_T, sigma=1.0 [length]), was applied radially from a cylindrical "spool" surface (radius = 2.0 [length]), and excluded backbone particles (i.e., the chain formed by loop anchors and "gap" segments between them) from the axial core of the system.

-	To prevent the helix from unwinding from the tips, we fixed the angular positions of the chromosome's two terminal particles (T1 and T2). We achieved this by adding four extra immovable particles: two on opposite ends of the cylinder axis (A1 and A2), and two more (P1 and P2) at the initial positions of T1 and T2. HOOMD torsional potentials were applied to the quadruplets T1-A1-A2-P1 and T2-A1-A2-P2 (setting k=100 kBT), fixing the angular positions of T1 and T2 without restricting radial motion.

-	To maintain the consistent size of each helical turn, we additionally restricted the angle of one loop anchor per helical turn using the same technique as above.

The resulting model had 5 geometric parameters (Figure S9A):

1. average loop size, l_loop_ [bp]
2. the size of gaps between anchors of consecutive loops, L_gap_ [nm]
3. the linear density of chromatin along the cylinder axis, ρ_L_ [Mb/µm]
4. the length of a turn in Mb, l_turn_ [Mb]
5. the volume density of chromatin, ρ_V_ [Mb/µm^3^].

The last parameter was used during the conversion of 3D coordinates into contact frequencies (see below):

6. “contact radius”, σ_C_ [nm]

The five geometric parameters, together with the genomic length of the simulated chromosome, l_chrom_ [Mb], and principles (i)-(iv) were sufficient to fully define the model, allowing the following geometric parameter to be derived.:

1. the length of the confining cylinder [nm], L_cyl_ = l_chrom_ /ρ_L_
2. the radius of the confining cylinder [nm], R_cyl_ = √(l_chrom_/ρ_V_/L_cyl_/π)
3. the pitch of the helix (i.e. the height of one complete helical turn) [nm], P = l_turn_ / ρ_L_
4. the expected number of loops in the chromosome, N_loops_ = l_chrom_ / l_loop_
5. the length of the chromosomal backbone (i.e., the chain formed by loop anchors and "gap" segments) [nm], L_bb_ = N_loop_ * (L_gap_+L_loopbond_)

Two of these parameters, (4) and (5), can be defined from experimental datasets. Previously, we demonstrated that models with helical backbones produce *in silico* Hi-C maps with 2nd diagonals at genomic separations equal the size of the turn, l_turn_ (*6*). Here, we found that this relationship required a minor correction, so to reproduce our 30-minute Hi-C dataset, where the 2nd diagonal is located at 16.4 Mb, we had to set (4) l_turn_ = 17 [Mb]. We relied on the SBF-SEM measurements (see above) to set the volume density of chromatin (5) ρ_V_ = 44 [Mb/µm^3^], which corresponds to the specific volume of (16.5 nm)^3^ per nucleosome or (1.65 [length])^3^ per simulated particle. The rest of the parameters (1)-(3) and (6) were identified via systematic sweeps and comparisons against Hi-C (see below).

Calculation of the simulated *P*(*s*) curve

To generate *in silico* Hi-C maps from simulated chromosome conformations, we found neighboring particle pairs in 3D using the Scipy KD-tree algorithm (*113*). Since KD-trees use strict distance cutoffs, we proposed a novel scheme to calculate more realistic, distance-dependent contact probabilities. To achieve this, we perturb each particle's position by adding a random shift drawn from a multivariate Gaussian distribution (standard deviation σ_c_ along each axis). Following perturbation, we identify neighboring pairs within a 1.1 [length] distance cutoff. This process is repeated 100 times, and average contact frequencies between all particle pairs are calculated across the replicates. Finally, we calculated *in silico* *P*(*s*) curves from these contact frequencies, mirroring the procedure used for Hi-C pair analysis.

Model selection via comparison of simulated and experimental P(s) curves

To identify simulations best matching experimental Hi-C data on SMC3-AID/CAPH-AID cells at t=30 min, we systematically explored combinations of model parameters (1), (2), (3), and (6). Instead of directly varying linear chromatin density (ρ_L_), we swept over helical pitch (P), which is related to ρ_L_  through P = l_turn_ / ρ_L_.

For each parameter set, we conducted three simulation replicates. We averaged the resulting *P*(*s*) curves and calculated the root mean square (r.m.s.) deviation from the experimental *P*(*s*) in a log10-log10 scale. R.m.s was calculated for genomic distances (*s*) between 3 kb and 40 Mb, with both curves normalized to 1.0 at 3 kb. The top three parameter sets yielded by this procedure had l_loop_=400kb, L_gap_=80-120nm, P=400 nm, σ_C_=48nm.

Simulations of SMC3/CAPH2-depleted chromosomes

To model chromosomes formed solely by condensin I (SMC3-AID/CAPH2-AID cells), we modified our approach. Instead of cylindrical confinement (iii), we linearly extended the chromosome within periodic boundary conditions. The chromosome's ends were fixed in space at a distance L_stretch_ = l_chrom_/ρ_L_. The periodic box length matched L_stretch_, and its width was adjusted to achieve the target volume density (ρ_V_=77Mb/µm^3^, as measured by SBF-SEM).

To reduce the number of simulations and the associated computational costs, we determined the four free parameters (l_loop_, L_gap_, ρ_L_, and σ_c_) in a two-step procedure, leveraging the fact that the shape of *P*(*s*) is independent of ρ_L_ for genomic distances s < 2 Mb (Fig S10A).

Step 1: Defining loop structure. We simulated chromosomes without linear extension, varying l_loop_, L_gap_, and σ_c_.  Comparisons to experimental *P*(*s*) data for SMC3-AID/CAPH2-AID cells (3kb - 2Mb separation range) yielded the best estimates for l_loop_ and L_gap_.

Step 2: Defining linear extension. Using the values from Step 1, we then varied ρ_L_ and σ_c_.  Matching *P*(*s*) behavior in the 3 kb - 15 Mb range allowed us to determine optimal ρ_L_ and σ_c_ values.

The top three parameter sets yielded by this procedure had l_loop_=100kb,  L_gap_=10-40nm, ρ_L_=25 Mb/µm and σ_C_=48-51nm (Fig.S10B-C).

Simulations of SMC3-depleted chromosomes

In our models of chromosomes with both condensin I and II (SMC3-AID cells), we built nested loop structures. Our previous work suggested condensin II forms larger, stable loops subdivided by smaller, more dynamic condensin I loops (*6*). This nested structure was directly supported by our new Hi-C data in SMC3-depleted cells (see below).

We implemented a two-step modeling process. First, we generated a layer of loops with average length l_loop_^root^, as described previously. We then subdivided each of these into smaller loops (exponentially distributed with the average  l_loop_^nested^). For each of these loop arrays, we specified its own length of gap segments, L_gap_^root^ and L_gap_^nested^.

Finally, since SMC3-AID chromosomes exhibit cylindrical shapes and 2nd Hi-C diagonals, we imposed the cylindrical confinement (iii) and helical backbone (iv) features used in our condensin II model.

This model had 8 parameters: l_loop_^root^, l_loop_^nested^, L_gap_^root^, L_gap_^nested^ , ρ_L_ (or, pitch P), l_turn_, ρ_V,_ σ_C_. As previously, we took the target value of volume density from SBF-SEM data (ρ_V_= 77 Mb/µm^3^, Fig.6F-G), and l_turn_=10 Mb to reproduce the 2nd diagonal at 8.4 Mb.

Since a full 6-dimensional parameter sweep is computationally infeasible, we started with a model combining l_loop_^root^ and L_gap_^root^ values from condensin II simulations with l_loop_^nested^ and L_gap_^nested^ from condensin I simulations. Adjusting pitch (P) and σ_C_, we achieved an excellent fit to SMC3-AID Hi-C (t= 30 min) with P= 350 nm or 400 nm and σ_C_= 43 nm (Figure S10D, green curve in S10G).

We then wanted to test if the model based SMC3-AID data could independently support our previous finding of significant gaps between consecutive condensin II loops. Sweeping L_gap_^root^, P, and σ_C_ revealed that 80nm was indeed the global optimum for Hi-C fit (Figure S10D) and that L_gap_^root^ below 40nm do not yield well-matching P(s) curves. However, a second solution emerged with L_gap_^root^=40nm (Figure S10G, yellow curve). This alternative better matched optical microscopy data, predicting a narrower condensin distribution relative to DNA (compare Figure S10E to Figure 7I lower right and Figure S10F to Fig 3C lower right). Therefore, we propose L_gap_^root^=40nm as the solution reconciling both Hi-C and microscopy observations.

Simulations of chromosomes with overlapping condensin loops.

Finally, we explored whether our Hi-C and microscopy data on condensin II-only chromosomes (SMC3/CAPH2-AID) supported models where loop-extruding condensin II complexes could completely bypass each other. To model this scenario, we made two minor adjustments to our condensin II simulation approach:

(i) Independent Loop Placement: To reflect complete condensin II-condensin II bypassing, loop arrays were built using N_loop_ independently positioned loops with exponentially distributed lengths (average l_loop_) (for an example, see Figure S9F).

(iv) Enforced Helicity: With no continuous scaffold, helical conformation was maintained by pinning each loop anchor's angular position using a torsional potential. The target angle for loop i was θ_loop_^i^ = π * (x_left_^i^+x_right_^i^) / l_turn_, (where x_left_^i^ and x_right_^i^ are the genomic positions of the two anchors). Potential stiffness was tuned to k_B_T energy at angular deviation of 1.2 radian.

Through iterative search in a parameter space, we identified one set of parameters yielding a strong Hi-C data match (Figure S9G): l_loop_= 900 kb with one loop per 360 kb of chromatin (in this model, l_chrom_ = 135 Mb, l_turn_= 17.5 Mb).

Our findings suggest that Hi-C data alone may not be sufficient to definitively distinguish between models with and without condensin II bypassing. Importantly, a crucial disagreement arose upon visualizing the 3D conformation of the modeled chromosome. Condensins II did not form a scaffold and appeared distributed across the entire chromosome width (Figure S9H), in stark contrast to microscopy images that clearly showed a distinct scaffold in SMC3-CAPH chromosomes (Figure 7C). This strong contradiction with the microscopy data argued against the possibility of complete loop extrusion bypass by condensin II.

Estimation of loop extrusion speed from chromosome morphology dynamics.

Microscopy images revealed that chromosomes achieve their cylindrical shape in around 10 minutes.  Concurrently, Hi-C analysis of both SMC3-AID and SMC3-AID/CAPH-AID cells shows that the peak in the *P*(*s*) derivative at the loop size reaches its maximum value at the same time point of 10 minutes. This combined evidence suggests that by 10 minutes, most chromatin has been extruded into loops by condensin II complexes, causing gaps between adjacent condensins to reach their near-minimal value.

We have previously demonstrated how gaps within the extruded loop array can be estimated as g=e^-λ/d^ (*12*), where g is the fraction of chromatin within gaps; λ is the extruder processivity, i.e. maximally possible loop size per extruder, λ=v_extrusion_ * time; d is average extruder spacing (d=l_chrom_/N_extruders_). Using this relation, we could calculate extrusion speed (v_extrusion_) from the gap fraction (g) and time: v_extrusion_ = - d * ln(g) / time

To estimate the extrusion speed, we could make two further assumptions. (i) Given the stable DNA binding of condensin II in mitosis (*38*) the size of condensin loops will be limited by their collisions, allowing us to infer d=l_loop_= 400 kb.  (ii) The appearance of bottlebrush-shaped chromosomes in microscopy data by 10 minutes requires DNA in each gap to be 100-300% greater than the minimum 80 nm (assuming 1600 bp / 80 nm conversion). This corresponds to a gap fraction (g) of 0.004-0.012. Together, these numbers gave us an estimate of v_extrusion_ = 4-5 * 400kb / 10 min = 2.5-3 kb/sec.

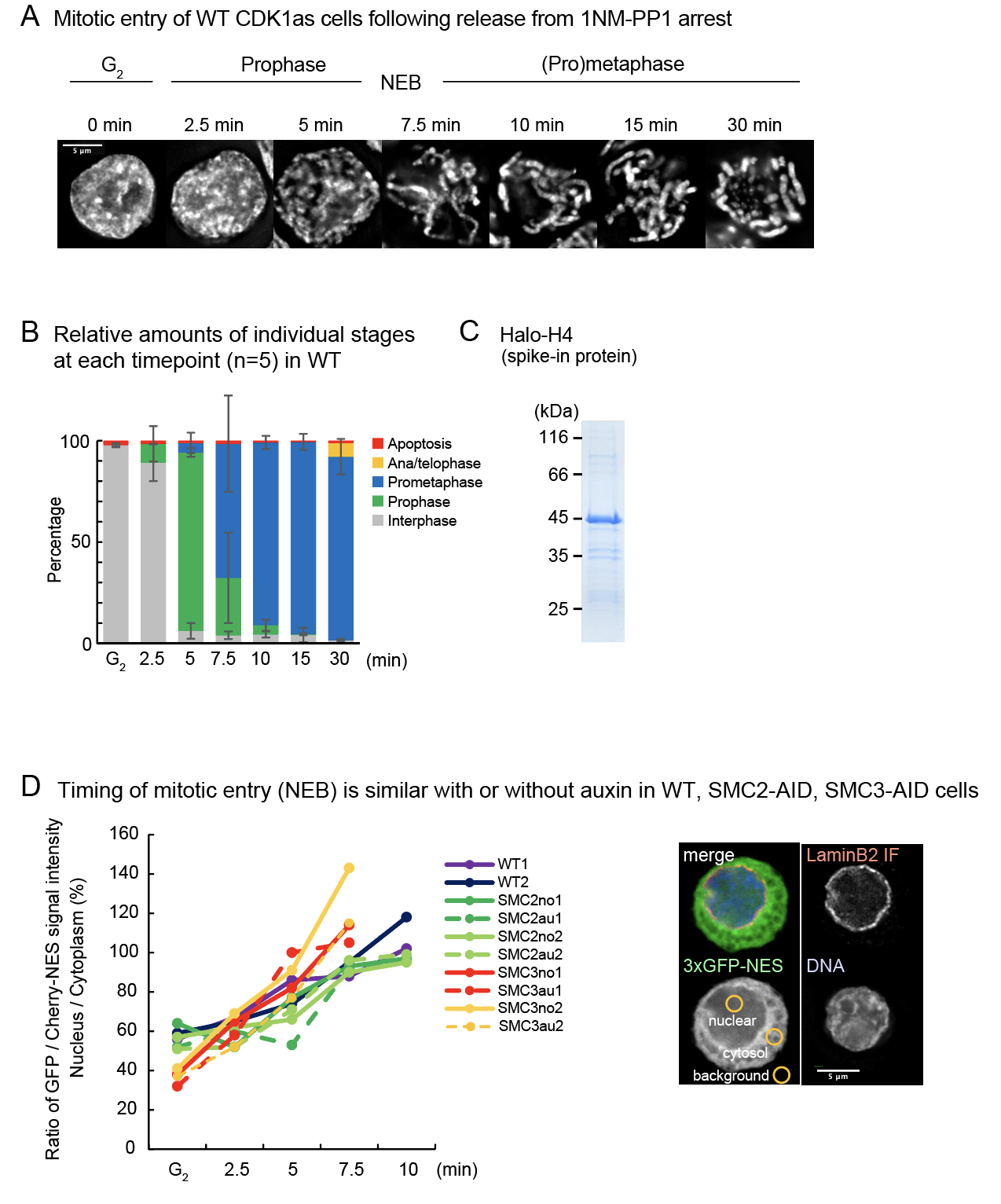

**Fig. S1. Mitotic entry of CDK1^as^ cells following release from 1NM-PP1 arrest.** (**A**) Representative images of DAPI-stained wild-type (WT) CDK1^as^ cells following release from G_2_ arrest. Cells were initially fixed with 1% formaldehyde and then with an ice-cold methanol/acetic acid solution. Scale bar: 5 µm. (**B**) Relative populations of individual mitotic stages at each time point taken from Hi-C samples of WT CDK1^as^ cells. Average and standard deviation (STDEV) are depicted (n = 5). (**C**) Purified Recombinant Halo-tagged Histone H4 protein, labeled with heavy arginine and lysine and used as a spike-in control (**D**) Timing of nuclear envelope breakdown (NEB) in WT, SMC2-AID, and SMC3-AID cells expressing 3xGFP-NES or 3xCherry-NES with or without auxin. Ratio of GFP or Cherry signal average intensity within the nucleus and cytoplasm is shown. A representative image of a G_2_ WT cell stained with Lamin B2 antibody, including representative yellow circles to measure the fluorescent intensities, is shown. The average of 20 cells is depicted for each time point with 2 biological replicates. Scale bar: 5 µm.

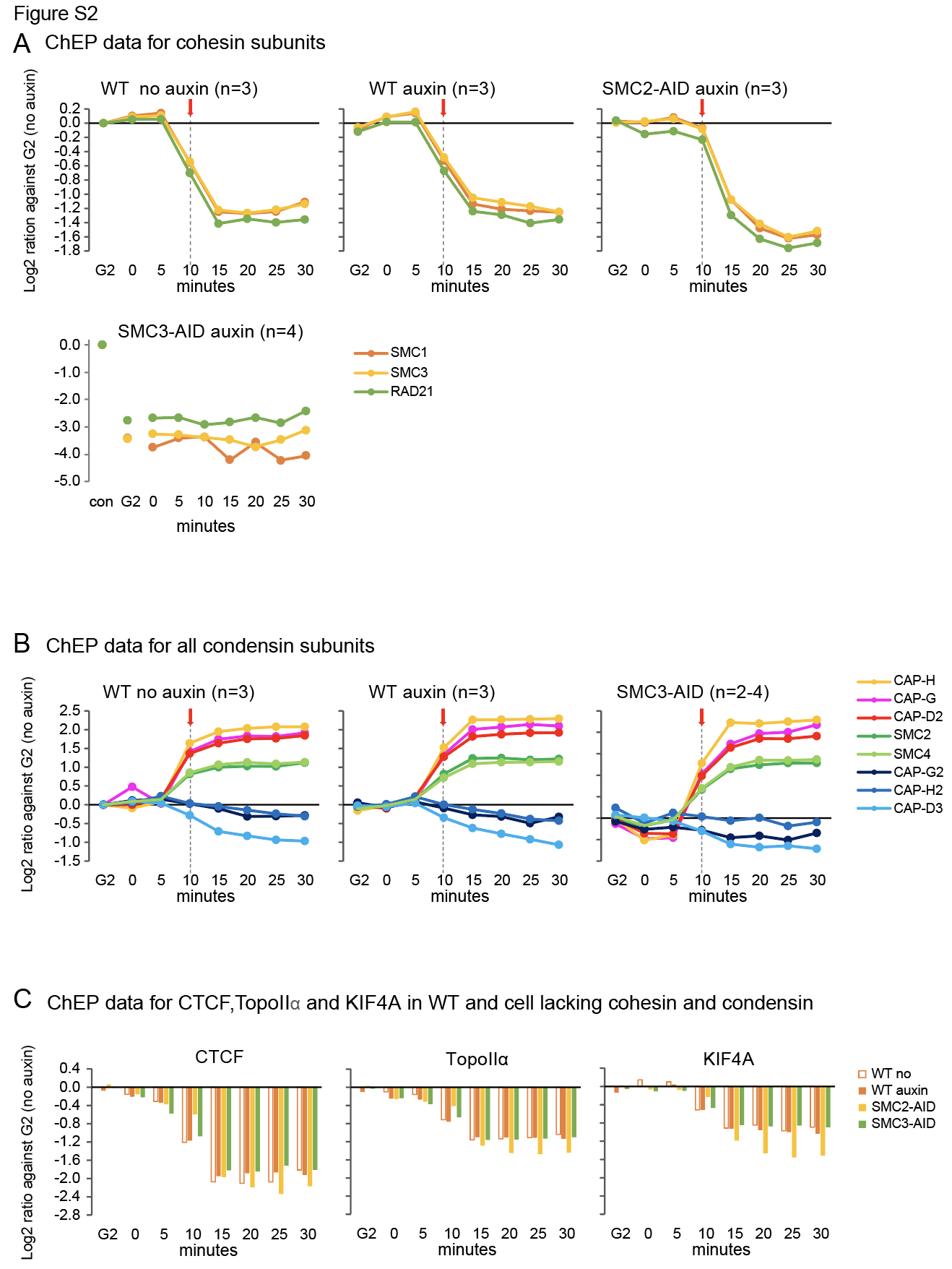

**Fig. S2. Quantification of key proteins associated with chromatin during mitotic entry.** (**A**) ChEP data for individual core cohesin subunits (SMC1, SMC3, Rad21) in WT cells no auxin (n=3), WT + auxin (n=3), SMC2-AID + auxin (n=3), and SMC3-AID + auxin cells (n=4). Curves show the Log2 ratio normalized against G_2_ without auxin for each cell line. Red arrows: timing of NEB. (**B**) ChEP data for individual condensin subunits in WT no auxin (n=3), WT + auxin (n=3), and SMC3-AID + auxin cells (n=2-4). Con (control) indicates G_2_ without auxin.  Four experiments were done, but as some condensin II subunits were not detected in some experiments, n=2-4 for SMC3-AID cells. Red arrows: timing of NEB. (**C**) ChEP data for CTCF, Topo IIα, and KIF4A, in WT cells with no auxin (n=3), WT + auxin (n=3), SMC2-AID + auxin (n=3), and SMC3-AID + auxin (n=4).

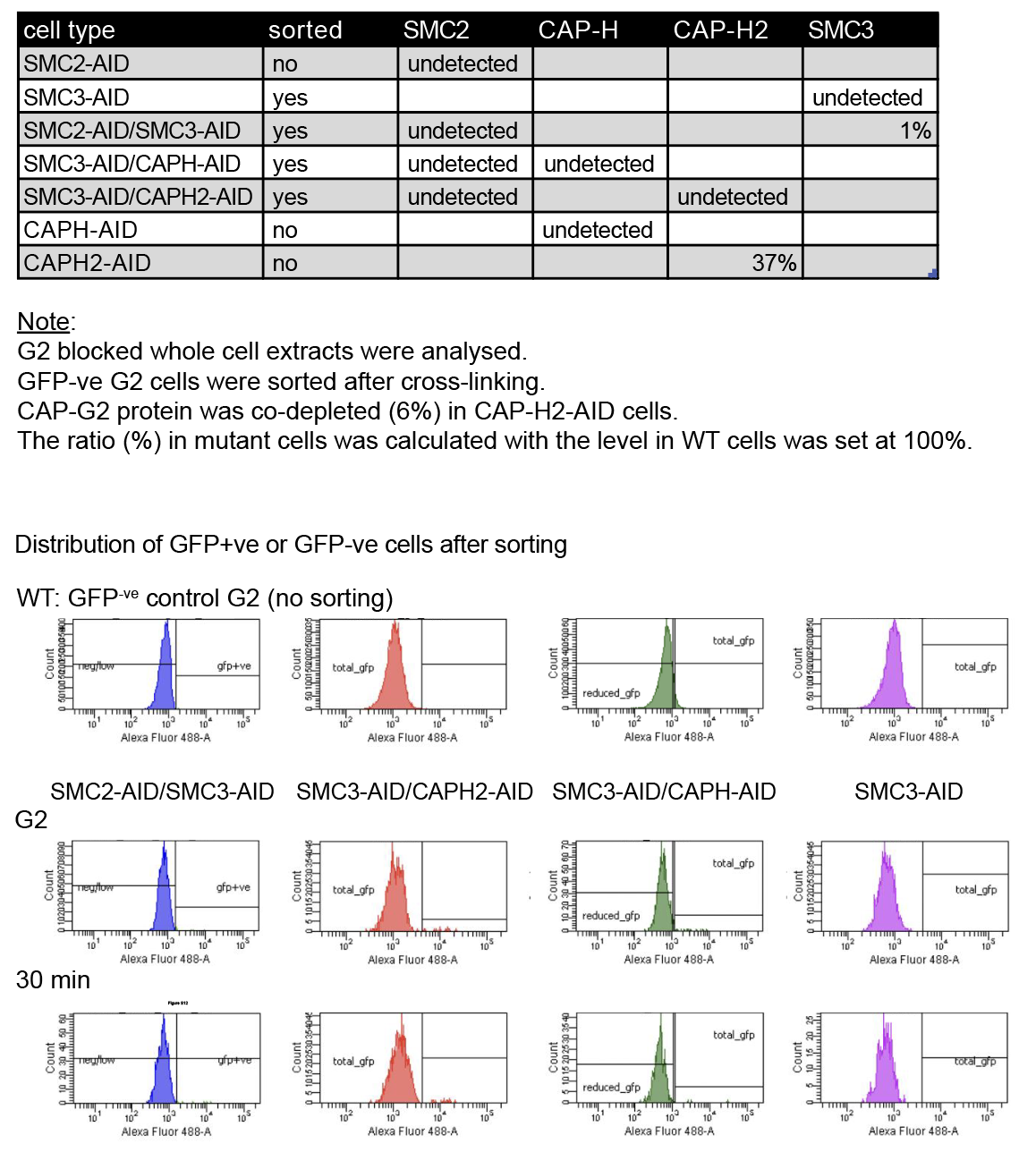

**Fig. S3. Level of target protein depletion in cells used to prepare Hi-C samples.** (**A**) Relative abundance of AID-GFP/clover tagged proteins in various mutant cells was compared to the abundance of the corresponding protein in wild type cells. Whole-cell mass spectrometry was performed on G_2_ cells used to prepare Hi-C samples for the following genotypes: WT, SMC2-AID, CAP-H-AID, CAP-H2-AID, SMC2-AID/SMC3-AID, SMC3-AID, SMC3-AID/CAP-H-AID, and SMC3-AID/CAP-H2-AID. Target proteins were fused with an mAID tag and a clover (GFP derivative) tag. Depending on the efficiency of depletion as determined from flow cytometry (next panels) some cells were sorted for GFP after auxin addition. (**B**) Distribution of GFP-positive and GFP-negative cells at G_2_ and at the 30 miniute time point after cell sorting. Control WT cells do not express GFP proteins, but the flow cytometer still detects auto fluorescence.

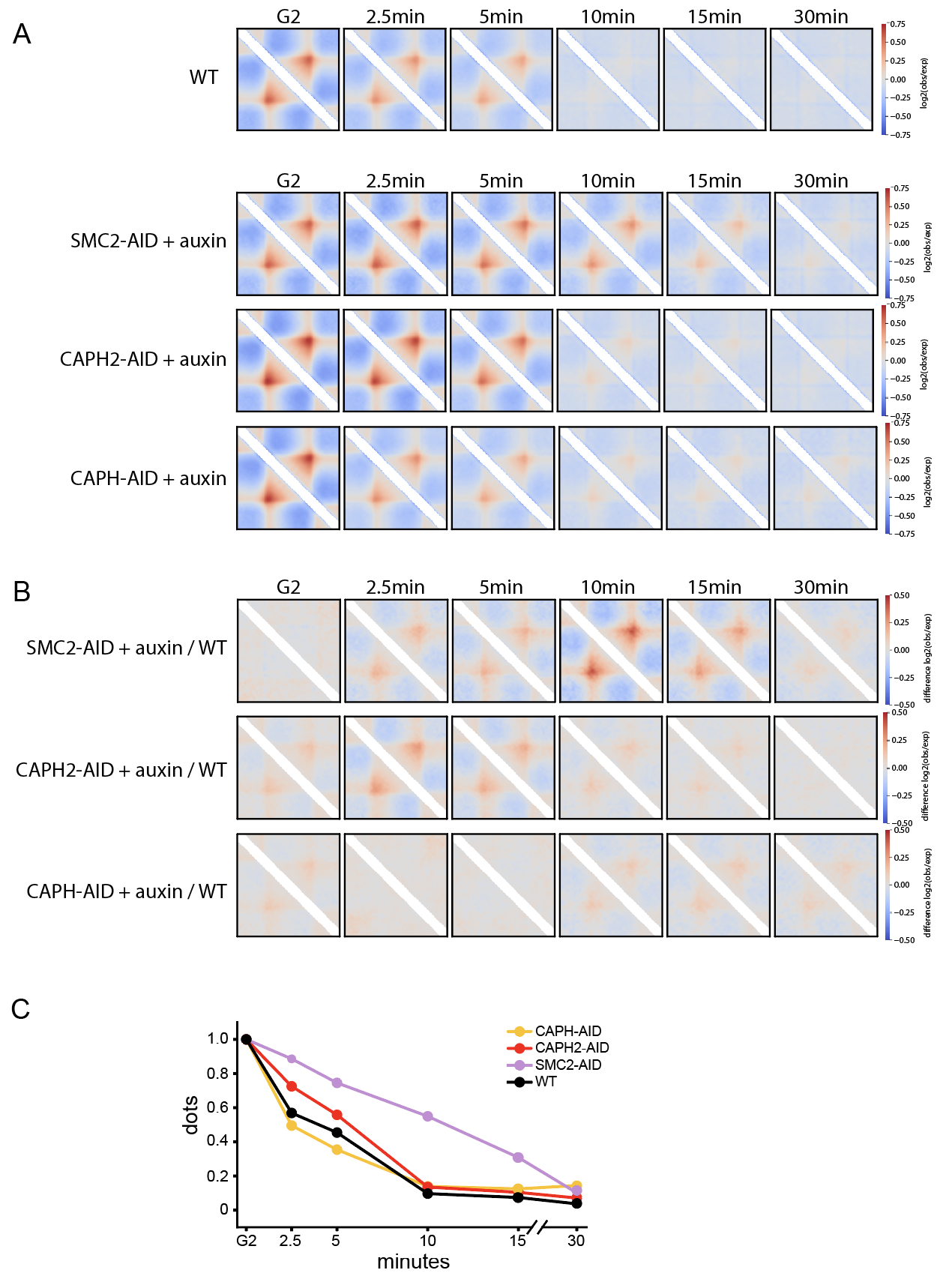

**Fig. S4. The dynamics of loop extrusion-related Hi-C features in SMC mutants.** (**A**) Top: Average contact footprints (pileups) of G_2_ -detected 100-700kb TADs throughout prophase. The contact maps are normalized by the distance-dependent decay of contact frequency (observed/expected). Bottom: as above, in the mutants of both condensins (top row), condensin II (middle row), and condensin I (bottom row). (**B**) The ratio of observed/expected pileups in each of the three mutants vs the WT pileup at the matching timepoints. (C) The average intensity of G_2_ -detected dots throughout prophase in WT and three condensin mutants.

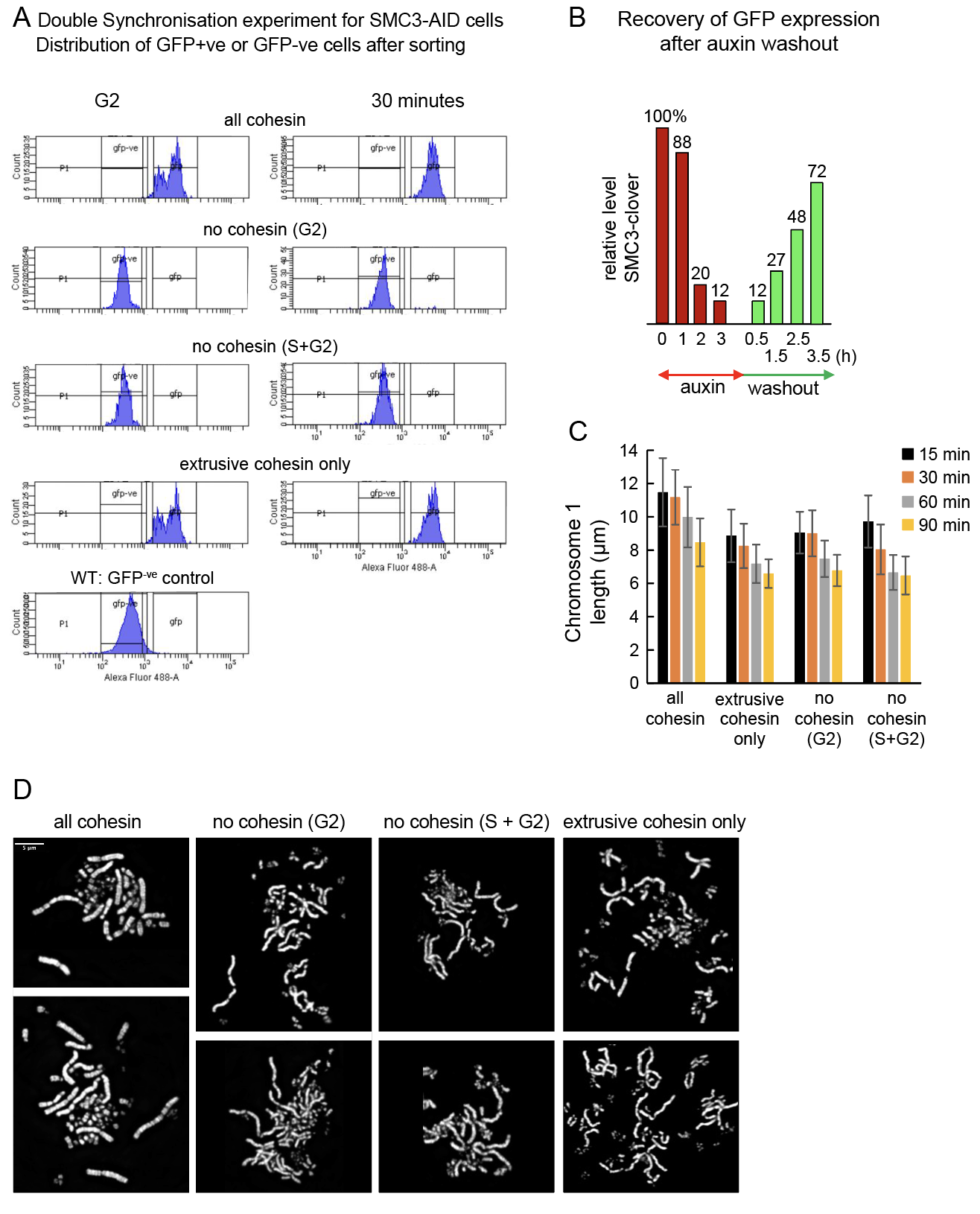

**Fig. S5. Double synchronization experiment with SMC3-AID cells to obtain populations with only loop-extruding cohesin.** (**A**) Level of SMC3-mAID-clover protein at G_2_ and 30-minute time points in various samples after cell sorting detected by flow cytometry. These cells were used for Hi-C and microscopy analysis. Control WT cells (bottom left) do not express GFP, and the signal is due to autofluorescence (**B**) Recovery of SMC3-mAID-Clover protein expression after auxin washout in an asynchronous cell culture detected by recovery of GFP fluorescence. The median fluorescence intensity at each time point was normalized against that at time 0. (**C**) Length of Chromosome 1 in chromosome spreads of double-synchronized SMC3-AID cells at 15, 30, 60, 90 minute time points. Measurements represent the average and STDEV from 16 cells at each time point (n=3). (**D**) Conventional chromosome spreads of SMC3-AID cells stained with DAPI. T=30 min. Scale bar: 5 µm.

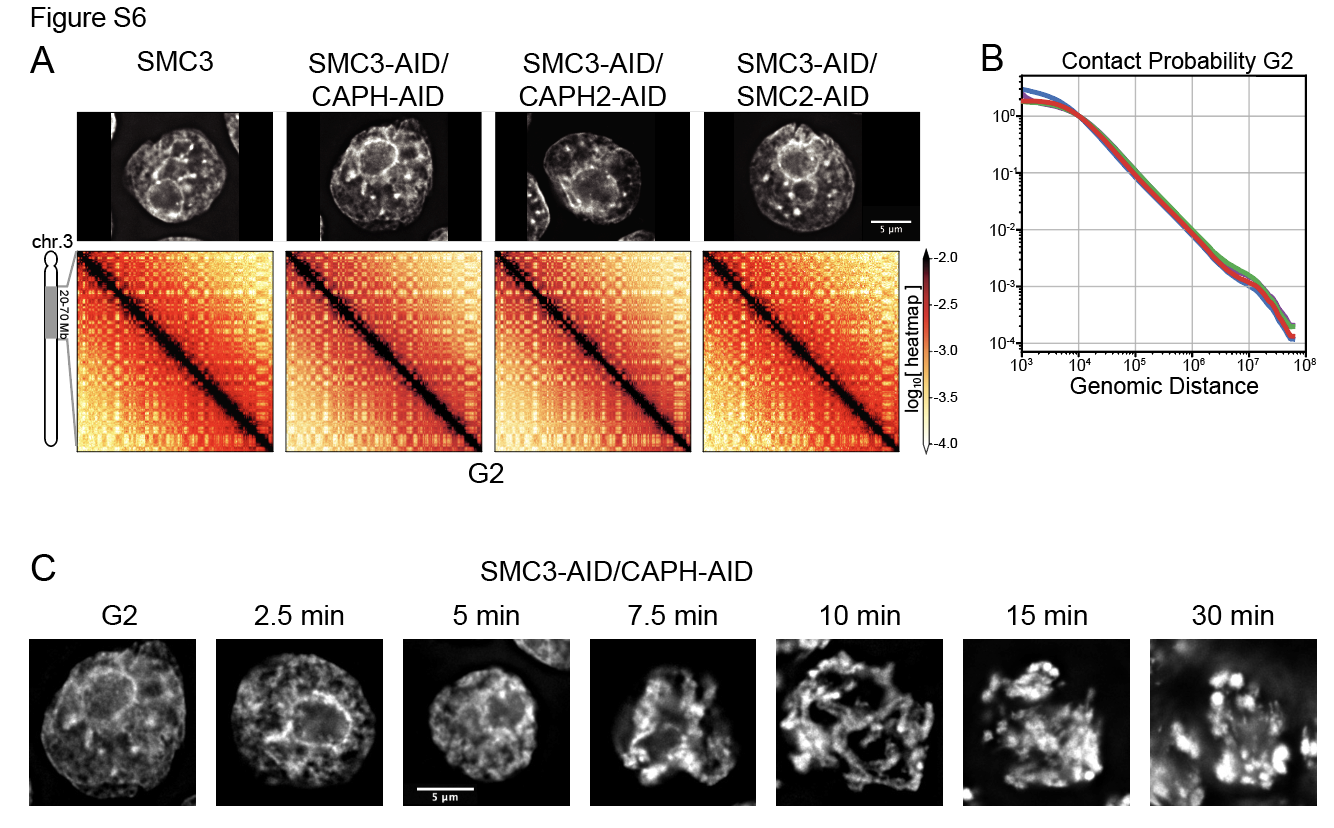

**Fig. S6. Organization of G_2_** **nuclei depleted of cohesin and condensin complexes.** (**A**) SMC3-AID, SMC3-AID/CAP-H-AID, SMC3-AID/CAP-H2-AID, SMC3-AID/SMC2-AID G_2_ cells plus auxin stained with DAPI and Hi-C heat maps of those cells. Scale bar 5 µm. (**B**) Contact frequency P(s) as a function of genomic separation (s) for Hi-C data shown in A. (**C**) Representative images of SMC3-AID/CAP-H-AID cells with auxin stained by DAPI at the indicated times after release from a G_2_ arrest. Cells were initially fixed with 1% formaldehyde and then with an ice-cold methanol/acetic acid solution. Scale bar: 5 µm.

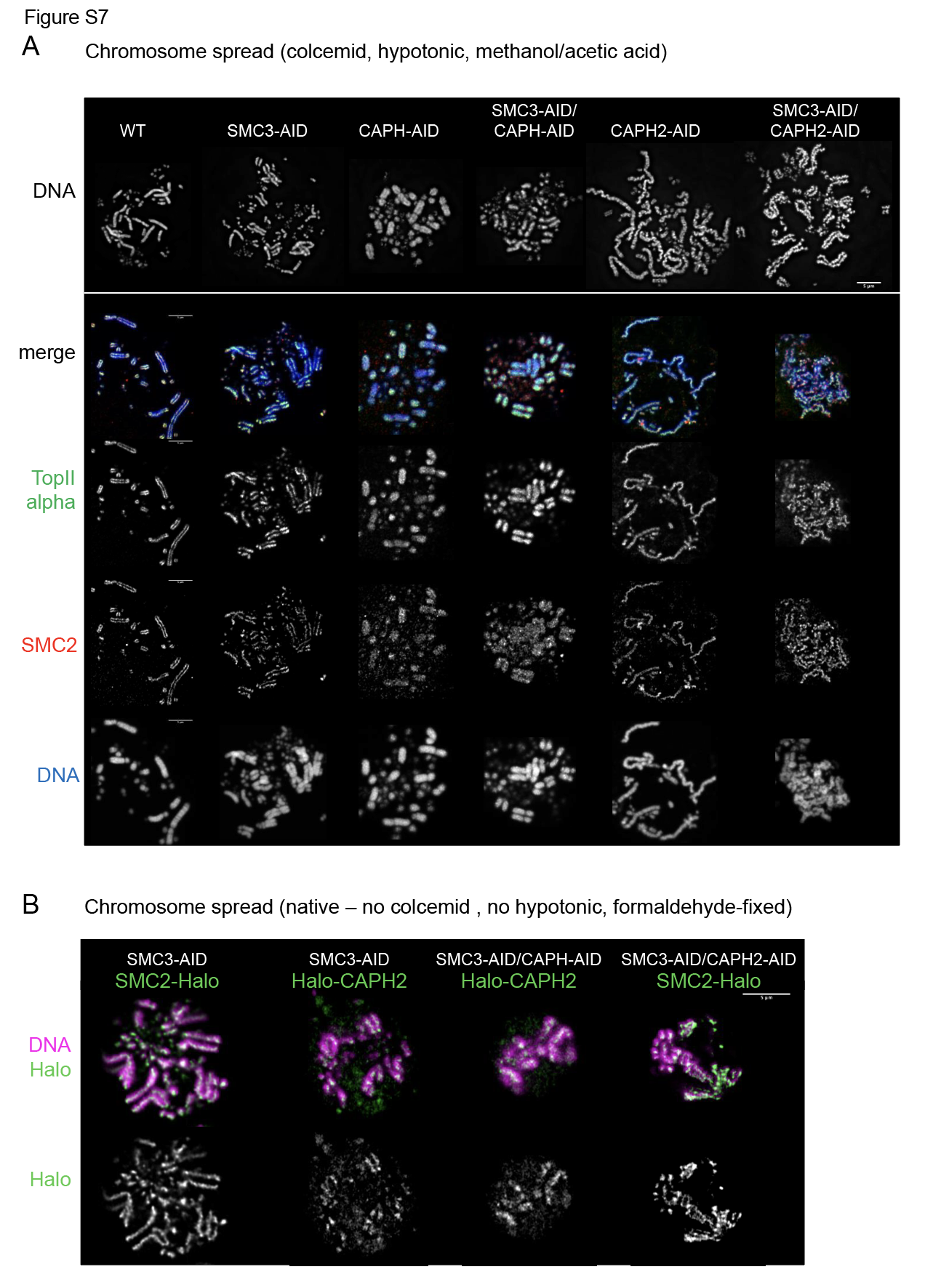

**Fig. S7. Axial localisation of chromosome scaffold proteins in WT and various mutant cells varies depending upon sample preparation method.** (**A**) Conventional chromosome spreads stained with DAPI and antibodies against Topo IIα and SMC2. Cells were treated with colcemid for 30 minutes after 1NM-PP1 washout (T=30 min), collected and then hypotonically swollen with 75 mM KCl for 10 minutes prior to fixation with cold methanol/acetic acid. A single Z section is shown. Note: Topo IIα and SMC2 localize diffusely throughout the entire chromosome in CAP-H-AID and SMC3-AID/CAP-H-AID cells after auxin treatment. Scale bar 5 µm. (**B**) For “Native” chromosome spreads, cells were rinsed with PBS and then fixed with 4% formaldehyde at T=30 min after release from G_2_ arrest. Halo-JFX549 (10,000) was added to the culture >30 minutes prior to 1NM-PP1 washout to label Halo-tagged proteins. A single Z section is shown. Note: In contrast to (A) Halo-CAPH2 is not diffuse throughout the chromosome in SMC3-AID/CAP-H-AID cells, but is confined to a more central region. Scale bar: 5 µm.

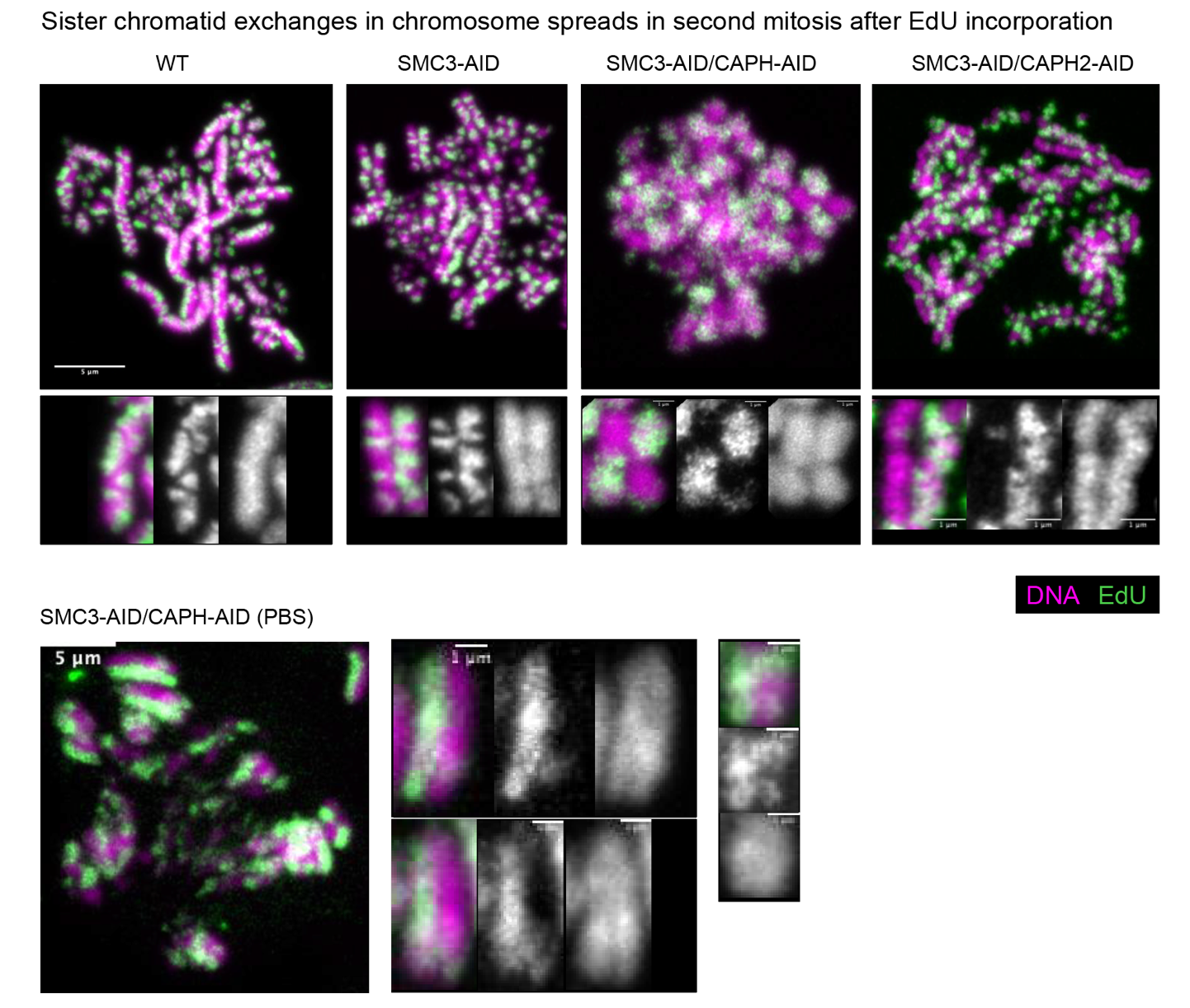

**Fig. S8. Sister chromatid exchanges in chromosome spreads in the second mitosis after EdU incorporation.** (**A**) Chromosome spreads of mitotic cells stained with DAPI to visualize DNA and azide-488 to visualize EdU. Cells were grown for one cell cycle in the presence of EdU, and then a second cycle without the drug. In the second mitosis, only one of each pair of sister chromatids is labeled with EdU. Conventional chromosome spreads were prepared as described in the legend to Figure S7A. Scale bar 5 µm (whole cell), 1 µm (enlarged image). (**B**) Similar to A, but “native” chromosome spreads were prepared by fixing cells depleted of SMC3 and CAPH with methanol/acetic acid solution in the absence of colcemid and hypotonic treatment. These chromosomes are longer than those prepared by conventional spreading, so some sister chromatid exchanges can be observed. In these chromosomes, which contain only much larger chromatin condensin II-extruded loops, the sister chromatid exchanges are diffuse and cannot be measured accurately.

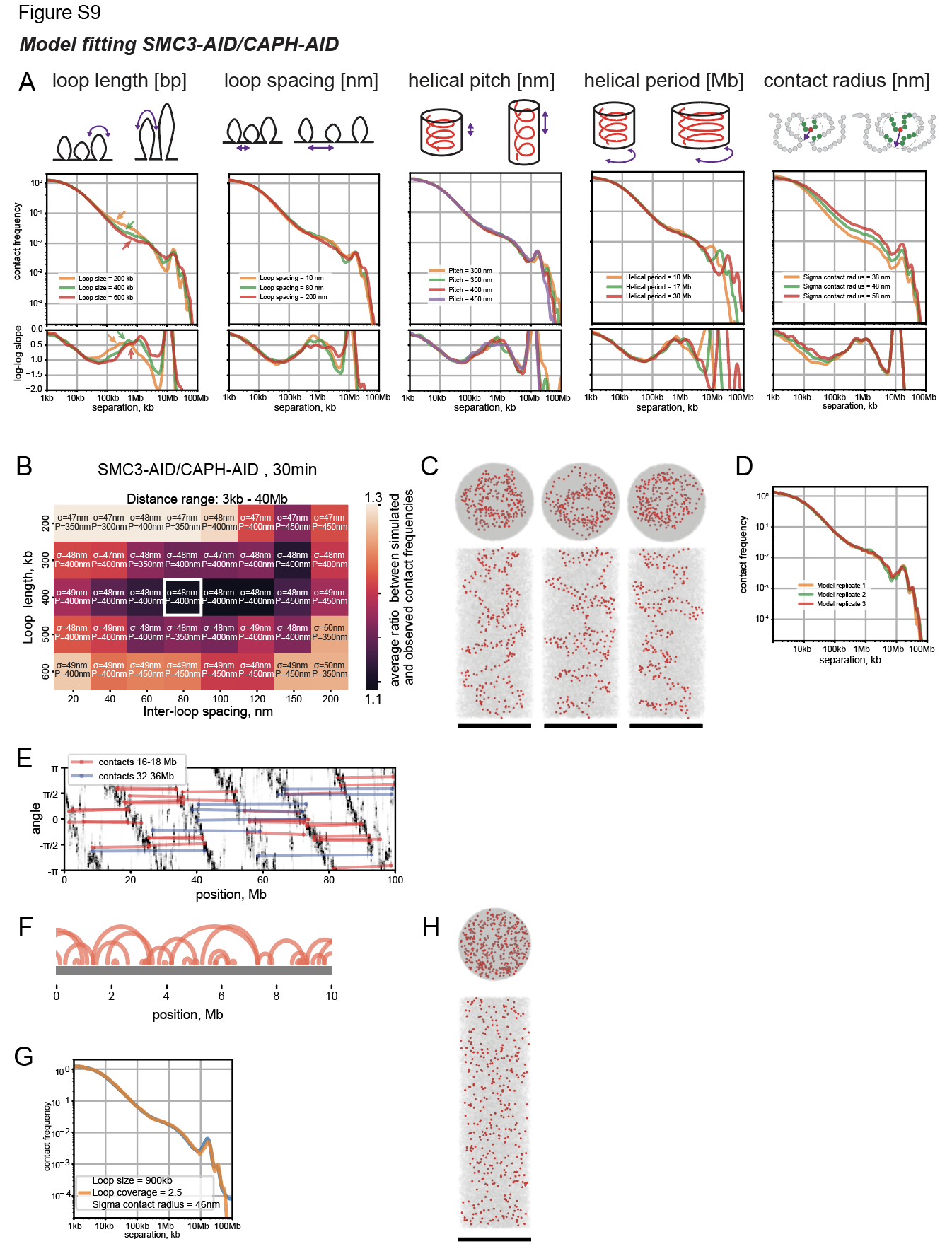

**Fig. S9. Fitting a polymer model of condensin II-only (SMC3-AID/CAPH-AID) chromosomes.** (**A**) The key five parameters of the model and their effect on the contact frequency P(s) vs genomic separation s curves. Top: the cartoon illustrating the effect of increasing the parameter on the polymer model. Middle: P(s) curves for several values of the parameter. Bottom: the slopes of P(s) curves in log-log space vs genomic separation s. In all models, unless specified, loop size=400kb, inter-loop spacing=80nm, helical pitch=400nm, helical period = 17Mb and sigma contact radius = 48nm. For slope calculation, P(s) was smoothed with sigma_log10 = 0.12. (**B**) The discrepancy between simulated and experimentally measured P(s) curves as a function of four model parameters. The color of each pixel shows the discrepancy of the best-fitting model from a group of models with two parameters (loop length and inter-loop spacing) specified by the x- and y- coordinates of the pixel. The annotation of each pixel shows the best-fitting values of the other two parameters (sigma contact radius and helical pitch) within the group specified by the x- and y- coordinates. The white square shows the parameters of the model selected for figure 7. (**C**) Chromosome conformation in three simulation replicates of the best-fitting model. Chromatin shown in gray, condensins II in red. (**D**) P(s) curves obtained from individual simulation replicates. (**E**) The 2D-histogram of the angular positions of chromatin particles (y-axis) vs their genomic position (x-axis) in the best-fitting model. The diagonal striped pattern originates from the helical organization of the chromosome. The colored lines indicate 30 randomly selected 3D contacts from the 2nd (red) and 3rd (blue) Hi-C diagonals. These lines illustrate that these diagonals are produced by interacting particles from consecutive gyres of the helix. (**F-H**) The best-fitting polymer model with overlapping condensin loops (F) can reproduce Hi-C P(s) (D), but lacks backbone-like staining of condensins (H).

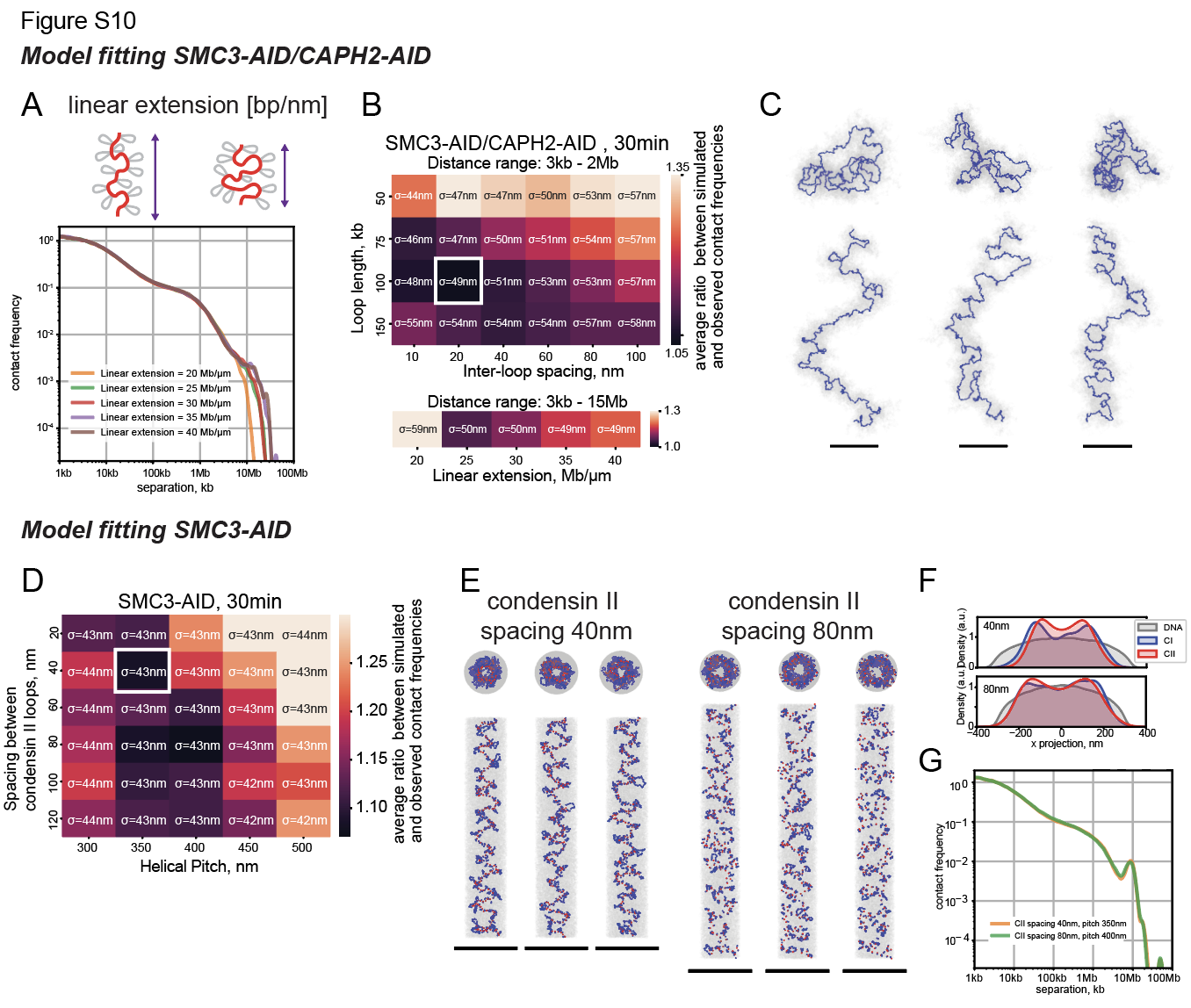

**Fig. S10. Fitting polymer models of condensin I-only (SMC3-AID/CAPH-AID) and condensin I+II (SMC3-AID) chromosomes.** (A-C) Fitting a model of condensin I-only chromosomes. (**A**) The effect of the linear extension of the chromosome on the contact frequency P(s) vs genomic separation s curve. In these simulations, loop size = 100kb, inter-loop spacing = 20nm, and sigma contact radius = 50nm. (**B**) The discrepancy between simulated and experimentally measured P(s) curves as a function of model parameters. Top: the discrepancy of P(s) in the 3kb-2Mb range in models without linear extension. The color of each pixel shows the discrepancy of the best-fitting model from a group of models with two parameters, loop length and inter-loop spacing, specified by the x- and y- coordinates of the pixel. The annotation of each pixel shows the best-fitting value of sigma contact radius. The white square shows the parameters of the model selected for figure 7. Bottom: the discrepancy of P(s) in the 3kb-15Mb range for models with fixed loop length and inter-loop spacing as a function of linear extension. (**C**) Chromosome conformation in three simulation replicates of the best-fitting model. Chromatin is shown in gray, condensins II in red. (**D-G**) Fitting a model of condensin I+II chromosomes. (D) The discrepancy between simulated and experimentally measured P(s) curves as a function of spacing between condensin II loops, helical pitch, and sigma contact radius. The white square shows the parameters of the model selected for figure 7. (E) Chromosome conformation in three simulation replicates of the best-fitting model with condensin II spacing of 40nm (left) and 80nm (right). (F, G) The cross-sectional distribution of condensins (F) and P(s) (G) in models as a function of condensin II spacing.

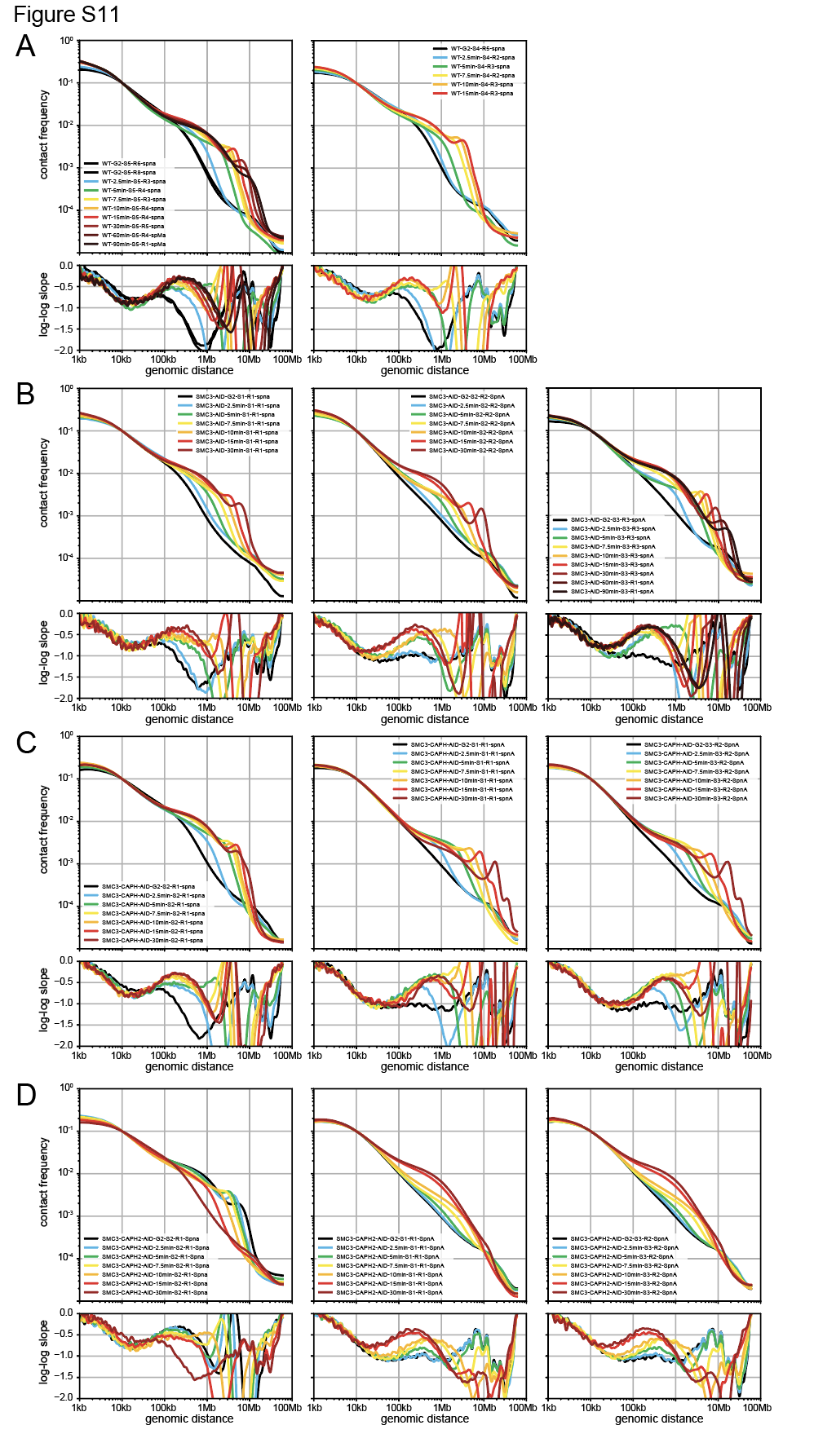

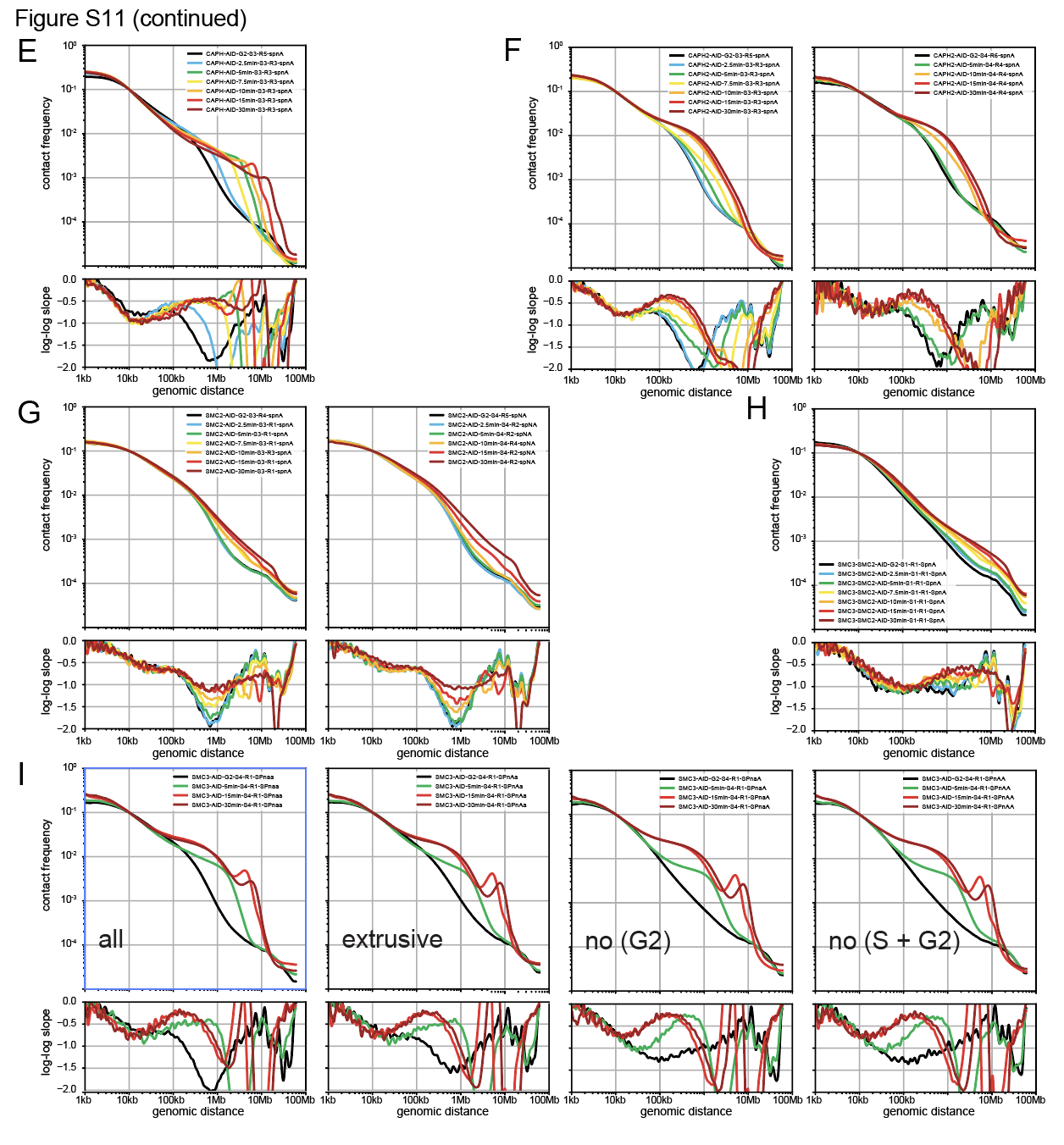

**Fig. S11. Plots of contact frequency vs genomic separation, P(s) for all Hi-C libraries used in this study.** (**A-I**) The contact frequency P(s) vs genomic separation (s) curves (upper) and their log-log derivatives (lower) for all cell lines.

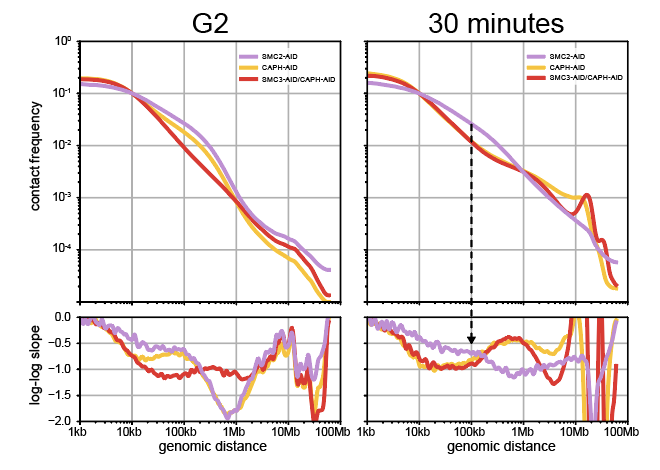

**Fig. S12. Condensin II removes cohesin loops.** The contact frequency *P(s)* vs genomic separation (*s)* curves for SMC2-AID cells (lacking both condensins, but having cohesins), CAPH-AID (having cohesins and condensin I), and SMC3-AID/CAPH-AID (having condensins II only) in G_2_ (left) and prometaphase (right). The characteristic bump on the P(s) at ~100kb (black arrow) reveals the presence of cohesin loops in G_2_ samples with intact cohesins (SMC2-AID and CAPH-AID), but not in the sample depleted of cohesin (SMC3-AID/CAPH-AID). In prometaphase, cohesin loops persist in the sample lacking both condensins (SMC2-AID), but cohesin loops are gone in the sample with active condensin II (CAPH-AID).

**Table S2. The size of DAPI stained area in SMC2-AID cells.** SMC2-AID cells were treated with auxin and nocodazole. DAPI stained areas (max projection) were measured using Fiji.

|  | No of cells | Area (cm^2^) |
| --- | --- | --- |
| G_2_ | **72** | **80.8** |
| 30 min | **96** | **41.8** |

**Table S3. Gyre size of chromosomes**

|  | SMC3-AID |  | WT |
| --- | --- | --- | --- |
| Experiment # | Mbp/gyre |  | Mbp/gyre |
| 1 | 8.99 |  | 7.27 |
| 2 | 9.84 |  | 6.75 |
| 3 | 10.6 |  |  |
| 4 | 12.2 |  |  |
| 5 |  |  | 8.2 |
| average | 10.41 |  | 7.41 |
| stdev | 1.36 |  | 0.73 |

**Table S4. Measurements vs modeling of chromosome length and width**

| Method | Measured (µm) | SMC3-AID | SMC3-AID/CAP-H-AID | SMC3-AID/CAPH2-AID |
| --- | --- | --- | --- | --- |
| LM | length | 7.7 ± 0.9 | 4.4 ± 0.5 | 13.6 ± 2.8 |
| EM | length | 6.3 | 5.3 | 12 |
| Simulations | end to end | 7 | 4.6 |  |
| LM | Tel — Tel | 6.9 ± 0.8 | 3.7 ± 0.5 | 8.6 ± 1.4 |
| Simulations | end to end |  |  | 8 |
| EM | width | 0.8 ± 0.12 | 1.2 ± 0.17 | 0.6 ± 0.15 |
| Simulations | width | 0.6 | 1.1 | 0.6 |

**Table S5: Second diagonal positions measured from P(s) derivative peaks (Mb)**

| **Time (min)** | **7.5** | **10** | **15** | **30** | **60** | **90** |
| --- | --- | --- | --- | --- | --- | --- |
| **WT** |  | 2.97 | 4.03 | 6.10 |  |  |
| **SMC3-AID** | 2.86 | 3.82 | 5.19 | 8.43 | 11.44 | 13.95 |
| **SMC3-CAPH-AID** |  |  | 6.55 | 16.40 |  |  |

**Table S6: List of CDK1^as^ cell lines.** All tags (mAID-Clover, clover, Halo) were knock-into the N-terminus or the C-terminus of the corresponding genes except CAP-H2-AID cells  which utilized a cDNA to express CAP-H2-mAID-GFP. CAP-H2 gene was knocked out in this case. Previous AID tagged cell lines were made over conditional knock out cell lines (see (*6*)).

| Cell line | knock-in tag | | random integration | |
| --- | --- | --- | --- | --- |
|  | N-terminus | C-terminus |  |  |
| wt |  |  |  |  |
| SMC2-AID (new) |  | mAID-Clover |  | OsTIR1 |
| SMC3-AID |  | mAID-Clover |  | OsTIR1 |
| SMC2-AID/SMC3-AID |  | mAID-Clover |  | OsTIR1 |
| CAP-H-AID (new) |  | mAID-Clover |  | OsTIR1 |
| CAP-H2-AID (new) |  |  | CAP-H2 cDNA-mAIDGFP | OsTIR1 |
| SMC3-AID/CAP-H2-AID |  | mAID-Clover | CAP-H2 cDNA-mAIDGFP | OsTIR1 |
| SMC3-AID/CAP-H-AID |  | mAID-Clover x2 |  | OsTIR1 |
| SMC3-Halo |  | Halo |  |  |
| CAP-H-Halo |  | Halo |  |  |
| Halo-CAP-H2 | Halo |  |  |  |
| wt 3XGFP-NES Halo-LaminB1 | Halo |  | 3XGFP-NES |  |
| SMC2-AID 3XCherry-NES |  |  | 3XCherry-NES | OsTIR1 |
| SMC3-AID 3XCherry-NES |  |  | 3XCherry-NES | OsTIR1 |
| SMC3-Clover/SMC2-Halo |  | Clover, Halo |  |  |
| SMC3-AID/SMC2-Halo |  | mAID-Clover, Halo |  | OsTIR1 |
| SMC3-AID/Halo-CAP-H2 | Halo | mAID-Clover |  | OsTIR1 |
| SMC3-AID/CAP-H2-AID/SMC2-Halo |  | mAID-Clover, Halo | CAP-H2 cDNA-mAIDGFP | OsTIR1 |
| SMC3-AID/CAP-H-AID/Halo-CAPH2 | Halo | mAID-Clover x2 |  | OsTIR1 |

**Table S7:List of oligos to construct guide RNA**. CAPH2 knock out guide RNA is cloned into CRIPSR-Cas9 vector from Thermofisher Scientific. All others are cloned into pX330 (addgene).

| gene | purpose | oligo |
| --- | --- | --- |
| SMC2 | knockin | CACCGAGTGAAGCCAGCAACAACA |
| SMC2 | knockin | AAACTGTTGTTGCTGGCTTCACTC |
| SMC3 | knockin | CACCGCAATTTCGCTTAACCATGCG |
| SMC3 | knockin | AAACCGCATGGTTAAGCGAAATTGC |
| CAPH | knockin | CACCGTCTGATGTCATTGTGAAACA |
| CAPH | knockin | AAACTGTTTCACAATGACATCAGAC |
| CAPH2 | knockin | CACCGCGTGACTCCACATCCTCCA |
| CAPH2 | knockin | AAACTGGAGGATGTGGAGTCACGC |
| CAPH2 | knockout | TTCTTGGTGAGGTCACGGATGTTTT |
| CAPH2 | knockout | ATCCGTGACCTCACCAAGAACGGTG |
| lamin B1 | knockin | CACCGTCCCCTACCATCACGTCACG |
| lamin B1 | knockin | AAACCGTGACGTGATGGTAGGGGAC |

**Table S1. Cell line characteristics for all libraries used in this study**

The name of the “Figure alias” (i.e. WT-2.5min-S4-R2-spna) is a combination of the following relevant information: cell type - timepoint - set - replicate - treatment. Sets of cells were derived from the same biological replicates and generally processed together. Set numbering continued from previously published sets (*6*). Treatments are sorting (S), palbociclib (P), nocodazole (N) and auxin (A), where capital letters indicate addition and small letters indicate omission thereof. Addition of MG132 was occasionally indicated with a capital M.

**Movies 1-4: 3D reconstructions of SBF-SEM datasets of all DT40 chromosomes.**

All datasets were modeled in Amira to differentiate chromosomes from the cytoplasm and then segmented to assign different colors to each individual chromosome. Movies are as follows: (**1**) Chromosome complement from wild-type DT40 cell; (**2**) Chromosome complement from SMC3-AID cell plus auxin; (**3**) Chromosome complement from SMC3-AID/CAP-H-AID cell plus auxin; (**4**) Chromosome complement from SMC3-AID/CAP-H2-AID cell plus auxin.

**Movies 5-8:  3D reconstructions of SBF-SEM datasets for chromosome 1.**

All datasets were modeled in Amira and segmented to assign different colors to each chromosome. Chromosome 1 was arbitrarily defined as here defined as the longest chromosome in each cell. Movies are as follows: (**5**) Chromosome 1 from wild-type DT40 cell; (**6**) Chromosome 1 from SMC3-AID cell plus auxin; (**7**) Chromosome 1 from SMC3-AID/CAP-H-AID cell plus auxin; (**8**) Chromosome 1 from SMC3-AID/CAP-H2-AID cell plus auxin.
